# Ciliary regulation of melanogenesis unveils moonlighting roles for Joubert Syndrome proteins

**DOI:** 10.64898/2026.09.27.754807

**Authors:** Brice Magne, Sara Forsey, Dawn C. Harper, Jiyu Zhang, Chloe E. Snider, Yuanqing Feng, Sarah A. Tishkoff, Alanna Strong, Theodore G. Drivas, Colin R. Goding, Robert J. Lee, Michael S. Marks

## Abstract

We previously found that skin tone correlates with intronic polymorphisms influencing the expression of the Joubert Syndrome (JS) ciliopathy gene *TMEM138*, suggesting that primary cilia might regulate pigment production in skin melanocytes. Herein, we show that many ciliary proteins regulate basal pigmentation in wild-type mouse and human melanocytes by facilitating primary cilium-dependent Melanocortin-1 receptor signaling, and downstream activation of the Microphthalmia transcription factor and melanogenic gene expression. Additionally, we identify an extra-ciliary moonlighting role for TMEM138 and other JS proteins in endolysosomal organization required for the biogenesis of melanosomes, the pigment-producing lysosome-related organelle in melanocytes. Our results provide new mechanistic insights into the physiological regulation of pigmentation and suggest that JS-specific pathology might be mediated by ciliary-independent mechanisms.

## Introduction

Pigmentation of the skin, hair and eyes provides natural protection against photodamage, vitamin B_9_ degradation, and cancer development (*1–4*). In the basal layer of the skin epidermis, melanin pigments are synthesized by melanocytes within specialized lysosome-related organelles (LROs), called melanosomes, and transferred to keratinocytes where they accumulate in nuclear caps to shield genomic DNA from ultraviolet (UV) radiation of the sun (*5–7*). Inactivating mutations in genes that regulate melanosome biogenesis or melanin synthesis (a.k.a. melanogenesis) underlie the pathogenesis of syndromic and non-syndromic forms of oculocutaneous albinism (*8*, *9*). In addition, non-pathogenic alleles of the same genes can influence the degree of physiological pigmentation and hence susceptibility to photodamage (*10*, *11*), underscoring complex genetic interactions regulating skin pigmentation and vision in humans. Despite the identification of several genetic variants driving physiological differences in human skin, hair, and eye color in genome wide association studies (GWAS) and other unbiased approaches (*12–23*), the full extent of genetic diseases that impact vision and skin pigmentation is not known. Identifying novel molecular components that control melanosome biogenesis or melanogenesis is thus essential to better understand the basic biology of skin and eye pigmentation, ethnic disparities that drive susceptibility to oculocutaneous diseases, and the etiology of rare diseases associated with pigmentary genodermatoses.

We recently showed that skin pigment variation in native African populations correlates with non-pathogenic polymorphisms in regulatory regions controlling *TMEM138* gene transcription (*12*, *18*), suggesting that *TMEM138* controls skin pigmentation in a dose-dependent manner. *TMEM138* encodes an ∼18 kDa transmembrane protein of unknown function that supports primary cilium biogenesis (i.e., ciliogenesis) in multiple cell types (*24*, *25*). Inactivating mutations in *TMEM138* or one of more than 35 other ciliogenic genes cause Joubert Syndrome (JS), a ciliopathy characterized by neurodevelopmental delay which can co-occur with ocular, hepatic, skeletal, and/ or renal abnormalities (*26*). Although ocular symptoms consistent with albinism such as retinal hypopigmentation and nystagmus are often observed in JS patients (*27*, *28*), clinical evidence linking JS and skin hypopigmentation has not yet been reported, perhaps because most patient cohorts are from light skin backgrounds in which hypopigmentation relative to close family members might not be obvious (*27–33*). Moreover, pigmentation has not been assessed in JS mouse models since most are lethal and/or not bred to a genetic background appropriate to assess coat color change (*34*). Thus, using new model systems to uncover why a link between TMEM138 and pigmentation was identified in our GWAS might reveal novel insights into JS pathogenesis.

In this study, we used highly pigmented cultured mouse and human melanocytes as tractable model systems to dissect novel cellular functions of *TMEM138* and other poorly understood ciliopathy gene products. We show that many ciliary proteins control basal pigmentation via G protein-coupled receptor (GPCR)-mediated signaling from primary cilia. We also uncover unprecedented moonlighting roles for TMEM138 and other JS proteins in endolysosomal regulation required for proper melanosome biogenesis. Our findings suggest that JS severity might reflect novel non-ciliary roles of JS gene products in the endolysosomal system and in LRO biogenesis.

### *TMEM138* and most other ciliopathy genes regulate basal melanogenesis in skin melanocytes

To assess the role of *TMEM138* in melanogenesis, we used either of two lentivirally encoded short hairpin RNAs (shRNAs) to silence its mouse ortholog (abbreviated here as *mTm138*) in stable lines of immortalized but non-cancerous melan-*Ink4a* melanocytes derived from C57BL/6J-*p16Ink4a*^null^ mice (*35*). These shRNAs reduced by 75-80% (sh*mTm138* #1) and 40% (sh*mTm138* #2) the levels of *mTm138* mRNA (by qRT-PCR; Suppl. Fig. 1A) and TMEM138 protein (by immunoblotting; Fig. 1A) relative to non-target shRNA (sh*NT*). Moreover, as assessed by immunofluorescence microscopy (IFM), *mTm138* depletion impaired both the formation and elongation of primary cilia in serum-starved mouse melanocytes commensurate with the degree of *mTm138* knockdown (Fig. 1B), extending findings in other cell types (*24*, *25*). By bright field microscopy and quantitative melanin content assays, cells expressing sh*mTm138* had reduced pigment content that also scaled with *mTm138* expression (Fig. 1C). Overexpression of HA-epitope tagged *human TMEM138* (HA-*hTM138*) had no significant effect in control cells, but restored pigmentation to near control levels in sh*mTm138* #1-knockdown cells (’rescue’; Fig. 1A, S1A, and 1C), confirming that hypopigmentation was caused by reduced *mTm138* expression. Consistently, reconstructed skin prepared from wild-type human fibroblasts and keratinocytes together with mouse melanocytes expressing either sh*NT*, sh*mTm138* #1, or sh*mTm138* #2 showed that visible pigment reduction scaled with *mTm138* knockdown levels (Fig. 1D), validating our findings in melan-*Ink4a* cell monocultures and echoing the phenotype observed in native African populations (*12*, *18*). Finally, depletion of *hTM138* mRNA in dark skin-derived primary human melanocytes by either of two lentivirally-encoded shRNAs (sh*hTM138*; Fig. S1B) impaired both ciliogenesis (following serum-starvation; Fig. 1E) and pigment content (Fig. 1F) commensurate with decreased *hTM138* expression, documenting that *TMEM138* is necessary for optimal ciliogenesis and melanogenesis in both mouse and human melanocytes.

**Figure 1.**
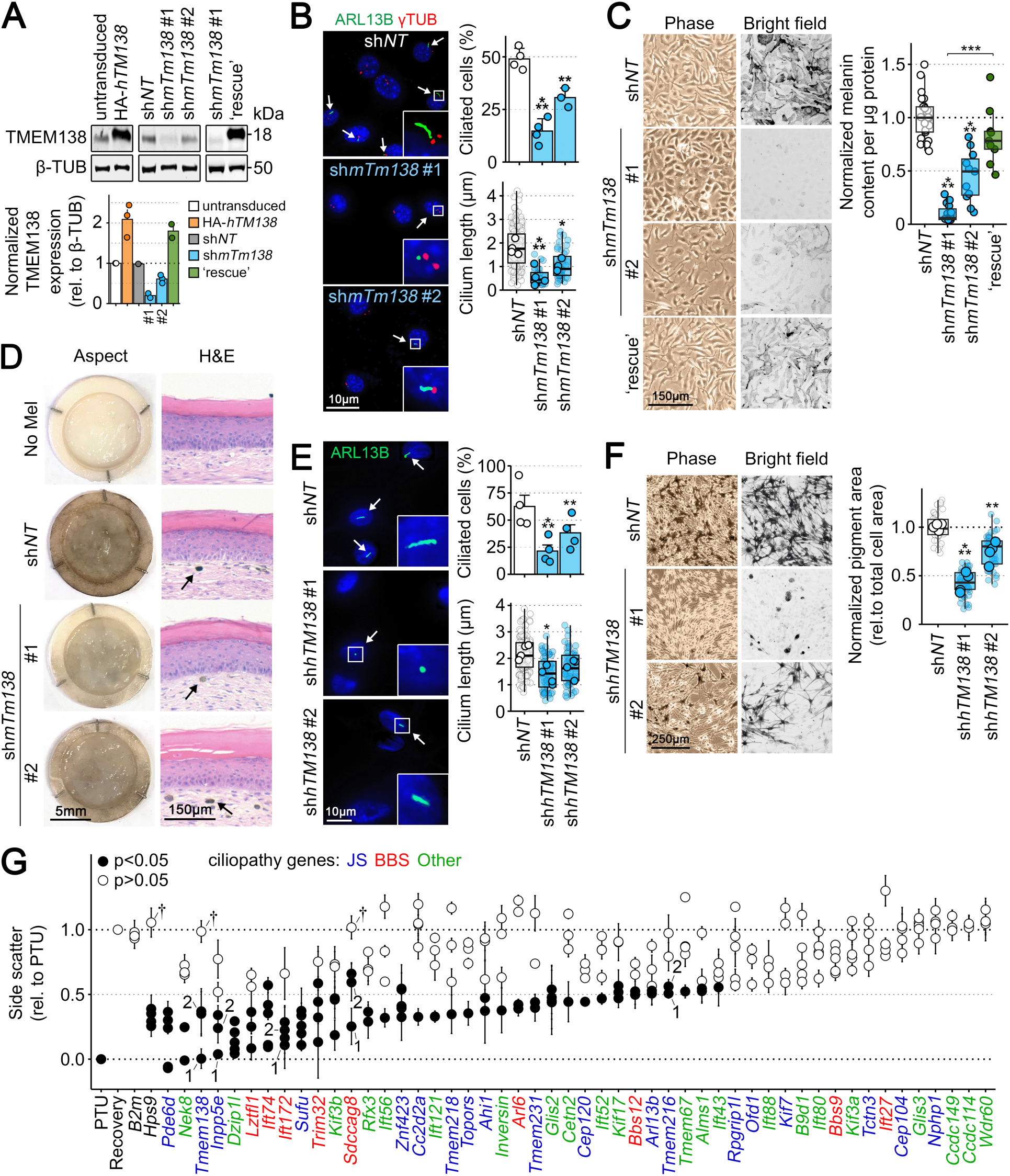
Melanogenesis is regulated by ciliary proteins. (**A** to **D**) Melan-*Ink4a* melanocytes were stably transduced with lentiviruses encoding a non-target shRNA (sh*NT*), or either of two *mTm138* shRNAs (sh*mTm138*). Overexpression and ‘rescue’ lines were generated by stable transduction of retroviruses encoding an shRNA resistant HA epitope-tagged, human *TMEM138* (HA-*hTM138*) in wild-type or sh*mTm138* #1-expressing cells, respectively. Untransduced melanocytes were used as control. (A) Top, immunoblotting of TMEM138 expression relative to β-Tubulin (β-TUB). Positions of nearby molecular weight markers are indicated at right in kDa. Bottom, quantification of TMEM138 band intensity normalized by untransduced cells across 1-3 experiments. (B) Left, representative IFM images of primary cilia. Arrows indicate ciliated cells. Right, proportion of ARL13B-labeled ciliated cells and cilium length among ciliated cells were quantified in 139-282 cells per condition across 3-4 experiments. (C) Left, representative images of phase and bright field contrast microscopy. Right, quantitative melanin content assay of stably transduced cells across 9-25 experiments. Values are normalized by melanin content of cells expressing sh*NT*. (D) Left, macroscopic aspect of reconstructed skin made from wild-type human skin fibroblasts and keratinocytes, containing either no melanocytes, or sh*NT*- or sh*mTm138*-expressing melan-Ink4a cells. Right, skin tissue sections were stained with Hematoxylin and Eosin (H&E) to highlight normal skin structure and melanocytes (arrows). Note that melan-Ink4a melanocytes were identified in the human skin dermis, likely because of species differences regulating melanocyte adhesion, migration, and proliferation. (**E** to **F**) Primary human melanocytes derived from four independent dark-skinned donors were transduced with lentiviruses encoding sh*NT* or either of two *hTMEM138* shRNAs (sh*hTM138*). (E) Left, representative IFM images of primary cilia. Right, graphs show proportion of ARL13B-labeled ciliated cells and cilium length in 26-87 cells per condition and per donor. (F) Left, representative images of phase contrast and bright field microscopy. Right, quantitative mean pixel intensity analysis over 10-15 bright field images per donor. Values are normalized by pigment area of cells expressing sh*NT*. (**G**) Flow cytometry-based shRNA screen analysis of pigment content. Graph shows mean values and error bars for each tested shRNA (circles) over 3-4 experiments. Statistical significance is color-coded for each shRNA, and black corresponds to p-values < 0.05. † highlights shRNAs known to have no impact on gene expression. Selected shRNAs used in the rest of this study are indicated with superscript numbers (1 or 2). Statistics: Data were analyzed using Fisher’s exact test (B and E, proportion of ciliated cells), or one-way ANOVA and Tukey’s multiple comparison test (all other datasets). *, p<0.05; **, p<0.01; ***, p<0.001.

To establish whether melanogenesis is regulated specifically by *mTm138* or more universally by ciliopathy genes, we designed an shRNA screen, adapted from a prior study (*36*), that uses the intrinsic light-scattering properties of melanin to assess pigment content. Melan-*Ink4a* cells were first transiently depleted of pre-existing melanin by a 2-week treatment with phenylthiourea (PTU), a reversible inhibitor of Tyrosinase (TYR; the rate-limiting enzyme in melanogenesis), and then transduced with lentiviruses encoding shRNAs to any of 50 ciliopathy genes or controls (note that knockdown efficiency of most shRNAs was not predetermined). After PTU removal and antibiotic selection, pigment recovery was assessed in live cells by flow cytometry using mean side-scatter as a proxy for melanin enrichment relative to cells transduced with sh*NT* remaining in PTU (set to 0) or following PTU removal (set to 1; Fig. S1C). Positive and negative controls included shRNAs to a known melanosome biogenesis gene (*Hps9*, encoding a subunit of Biogenesis of Lysosome-related Organelle Complex (BLOC)-1 required for cargo trafficking to maturing melanosomes (*37*, *38*)) or a pigment-unrelated gene (*B2m*), respectively. The results show that for most targeted ciliary genes at least one of several shRNAs caused a significant reduction in the recovery of side scatter, and hence pigment content (Fig. 1G). Particularly strong effects were observed using shRNAs targeting genes mutated in JS or Bardet-Biedl Syndrome (BBS), while little effect was observed for shRNAs targeting genes like *mCcdc114* that impact cilium motility. To validate these findings, we generated stable lines of melan-*Ink4a* cells depleted of two additional JS genes, *mTmem216* and *mInpp5e*, and two BBS genes, *mSdccag8* and *mIft172*, using effective shRNAs selected from the screen. We confirmed knockdown efficiency by RT-qPCR (40-80% reduction in mRNA expression; Fig. S1D) and observed various degrees of impaired ciliogenesis by IFM (Fig. S1E) and significant reduction in pigment content by quantitative melanin content assays (Fig. S1F) in all transduced cell lines. Of note, knockdown of *mTmem216* or *mSdccag8* in mouse melanocytes decreased melanin content but did not strongly impair ciliogenesis. While *mSdccag8* alters the function but not the biogenesis of primary cilia in mouse neurons (*39*), *hTMEM216* regulates both processes in kidney cells (*24*), suggesting that ciliogenic functions of *TMEM216* might vary across species or cell types (see Fig. 4 for evidence that shRNA knockdown was sufficient to interfere with non-ciliary functions of *mTmem216*). Altogether, these findings strongly support a role for JS, BBS, and other ciliopathy genes in constitutive melanogenesis within melanocytes.

### Ciliopathy genes regulate the expression of *MITF* and its target genes

Because primary cilia regulate signaling and downstream gene expression in most somatic cells (*40*, *41*), we next assessed whether reduced pigmentation in melanocytes depleted of JS or BBS genes was due to impaired melanogenic gene expression. By RT-qPCR, the expression of mRNAs for all pigment genes tested – *Tyr*, *Tyrp1*, *Pmel*, *Oca2*, *Dct*, *Mc1r*, and the master regulator of melanogenic gene transcription, *Mitf* (*42*) – was markedly reduced in stable melan-*Ink4a* lines depleted of *mTm138*, *mInpp5e*, *mSdccag8*, or *mIft172*, compared to untransduced or sh*NT*-expressing cells (Fig. 2A). Interestingly, mRNA expression of *Lamp2*, a non-melanogenic, lysosomal indirect target gene of *Mitf* (*43*), was also reduced albeit less dramatically than most other pigment gene, while mRNA expression of *Calr*, a non-melanogenic, non-target gene of *Mitf*, remained stable in ciliopathy gene depleted cell lines compared to controls (Fig. 2A). Melanocytes stably expressing shRNAs to *mTmem216* had no defect in melanogenic gene expression, correlating with intact ciliary function (Fig. S1E). These data suggest that primary cilia impact *Mitf* expression. By immunoblotting, we observed that melanocytes depleted of *mTm138* had reduced protein expression of TYRP1, PMEL (Fig. 2B), and TYR (Fig. S2A) compared to control cells, validating effects at both mRNA and protein levels.

**Figure 2.**
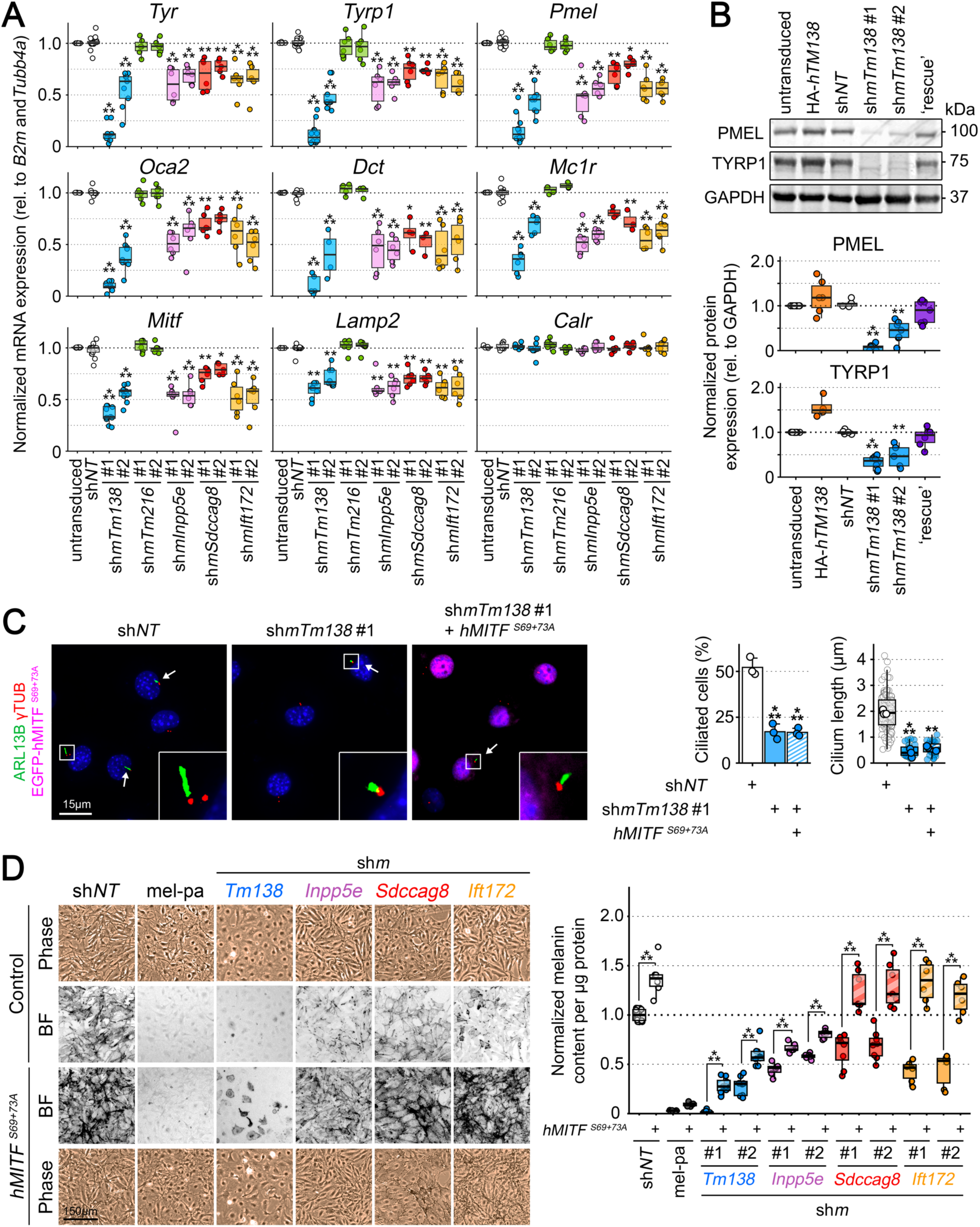
Optimal expression of *MITF* and its target genes requires ciliary proteins. Melan-*Ink4a* melanocytes that were untransduced or stably transduced with lentiviruses encoding a non-target shRNA (sh*NT*), or either of two shRNAs to *mTm138* (sh*mTm138*), *mTmem216* (sh*mTm216*), *mInpp5e* (sh*mInpp5e*), *mSdccag8* (sh*mSdccag8*) or *mIft172* (sh*mIft172*; see Fig. S1D for knockdown efficiency). **(A)** RT-qPCR analysis of mRNA expression levels for several melanogenic genes relative to *B2m* and *Tubb4a* housekeeping genes across 3-10 experiments. Values are normalized by mRNA expression levels of untransduced cells **(B)** Top, immunoblotting of PMEL and TYRP1 expression relative to GAPDH. Positions of nearby molecular weight markers are indicated at right in kDa. Bottom, quantifications of melanogenic protein expression relative to GAPDH over 4-9 experiments. Values are normalized by protein expression of untransduced cells. Overexpression and ‘rescue’ lines are described in Fig. 1A. **(C)** Left, IFM images of primary cilia. Cells expressing *hMITF^S69+73A^* show nuclear localization of active MITF; arrows point to ciliated cells. Insets are 4.25-fold magnifications of boxed regions. Right, percent ciliated cells and cilium length among ciliated cells were quantified in 126-212 cells per condition across 3 experiments. **(D)** A constitutively active form of human *MITF* (*hMITF^S69+73A^*) was expressed by stable lentiviral transduction, and BLOC-1-deficient melan-pa (mel-pa) melanocytes were used as an additional control. Left, representative images of phase and bright field contrast microscopy. Right, quantitative melanin content assay across 6-14 experiments. Values are normalized by melanin content of sh*NT*-expressing cells. Statistics: Data were analyzed using Fisher’s exact test (C, proportion of ciliated cells), or one-way ANOVA and Tukey’s multiple comparison test (all other datasets). *, p<0.05; **, p<0.01; ***, p<0.001.

To test whether ciliary proteins control melanogenic gene expression via *MITF* regulation, we expressed a constitutively active form of EGFP-tagged human *MITF* – *hMITF^S69+73A^* (*44*) – under the control of the weak hPGK promoter in stable lines of melan-*Ink4a* melanocytes depleted of *mTm138*, *mInpp5e*, *mSdccag8*, or *mIft172*. RT-qPCR analysis revealed that *hMITF^S69+73A^* expression restored mRNA abundance of melanogenic genes and *Lamp2* to the levels of control cells, without affecting *Calr* mRNA expression (Fig. S2B). However, *hMITF^S69+73A^* expression did not restore ciliogenesis in melanocytes expressing sh*mTm138* #1, demonstrating that *hMITF^S69+73A^*expression restores pigment gene expression independently of ciliary function (Fig. 2C). By bright field microscopy and quantitative melanin content assays, expression of *hMITF^S69+73A^* restored pigmentation completely in cells depleted of the BBS proteins *mSdccag8* or *mIft172*, but only partially in cells depleted of the JS proteins *mTm138 or mInpp5e* (Fig. 2D). This shows that pigment gene expression recovery is sufficient to reestablish pigmentation in BBS but not JS gene-deficient melanocytes. Moreover, consistent with previous findings (*45*), *hMITF^S69+73A^* expression in *Hps9*-deficient melan-pa melanocytes – which have reduced pigmentation due to defective BLOC-1-mediated trafficking to melanosomes (*38*, *46*) – only weakly increased pigment content (Fig. 2D) despite amplifying melanogenic gene expression as in control cells (Fig. S2B). This result shows that *MITF* activity minimally compensates for structural defects in melanosome biogenesis. Failure of *hMITF^S69+73A^* to fully restore pigmentation in *mTm138*- or *mInpp5e*-, but not *mSdccag8-* or *mIft172*-deficient melanocytes might thus reflect an additional non-transcriptional defect linked to JS but not BBS. Altogether, these findings support a role for primary cilia in basal melanogenesis through the control of *Mitf* and its target genes, and suggest a possible additional non-ciliary role specifically for JS genes in a post-transcriptional aspect of melanosome biogenesis.

### Activation of the Melanocortin-1 Receptor (MC1R)–MITF signaling axis requires ciliary function

To uncover how ciliary genes specifically regulate *MITF* expression and downstream pigment gene transcription, we asked whether defective ciliogenesis disrupts melanogenic signaling. One important stimulator of melanogenesis in melanocytes is the GPCR MC1R. MC1R stimulation activates adenylyl cyclase (ADCY) to produce cAMP, an activator of protein kinase A (PKA), which in turn phosphorylates the cAMP response element-binding protein (CREB), a transcription factor that drives *MITF* transcription (*47–49*). Under combined stimulation with UV light and the high-affinity MC1R ligand, α-Melanocyte-Stimulating Hormone (MSH), exogenously expressed GFP-tagged MC1R was shown to localize in part to primary cilia in melanoma and non-pigmented melanocytic cell lines (*50*). We thus hypothesized that ciliary function is required for physiological activation of endogenous MC1R–MITF signaling in pigmented, non-cancerous melanocytes.

To test this hypothesis, we first assessed the pigment response to a highly stable MSH analog (NDP-α-MSH) in serum-starved melan-*Ink4a* melanocytes with various degrees of cell ciliation (i.e., expressing either sh*NT*, sh*mTm138* #1, or sh*mTm138* #2; Fig. 1B). By bright field microscopy and quantitative melanin content assays, MSH stimulation increased pigmentation commensurate with the degree of cell ciliation (Fig. 3A), suggesting that primary cilia are required for effective MSH-induced pigmentation. Similar results were obtained after stimulating cells with Forskolin (FSK), a potent activator of ADCY (Fig. 3A). We confirmed by immunoblotting that CREB phosphorylation increased over time after MSH or FSK stimulation in ciliated (sh*NT*) but not unciliated (sh*mTm138* #1) cells (Fig. 3B). To rule out the possible effect of reduced *Mc1r* mRNA expression (Fig. 2A) on the MSH-induced pigment response of *mTm138*-depleted cells, we analyzed CREB phosphorylation in melan-*Ink4a* melanocytes expressing both sh*mTm138* #1 and *hMITF^S69+73A^* (sh*mTm138* #1/*hMITF^S69+73A^*cells) as these cells have defective ciliogenesis (Fig. 2C) but rescued *Mc1r* mRNA expression (Fig. S2B). MSH stimulation was unable to elevate CREB phosphorylation levels in these cells (Fig. 3B), demonstrating that the activation of the endogenous MC1R–MITF signaling axis specifically requires functional cilia.

**Figure 3.**
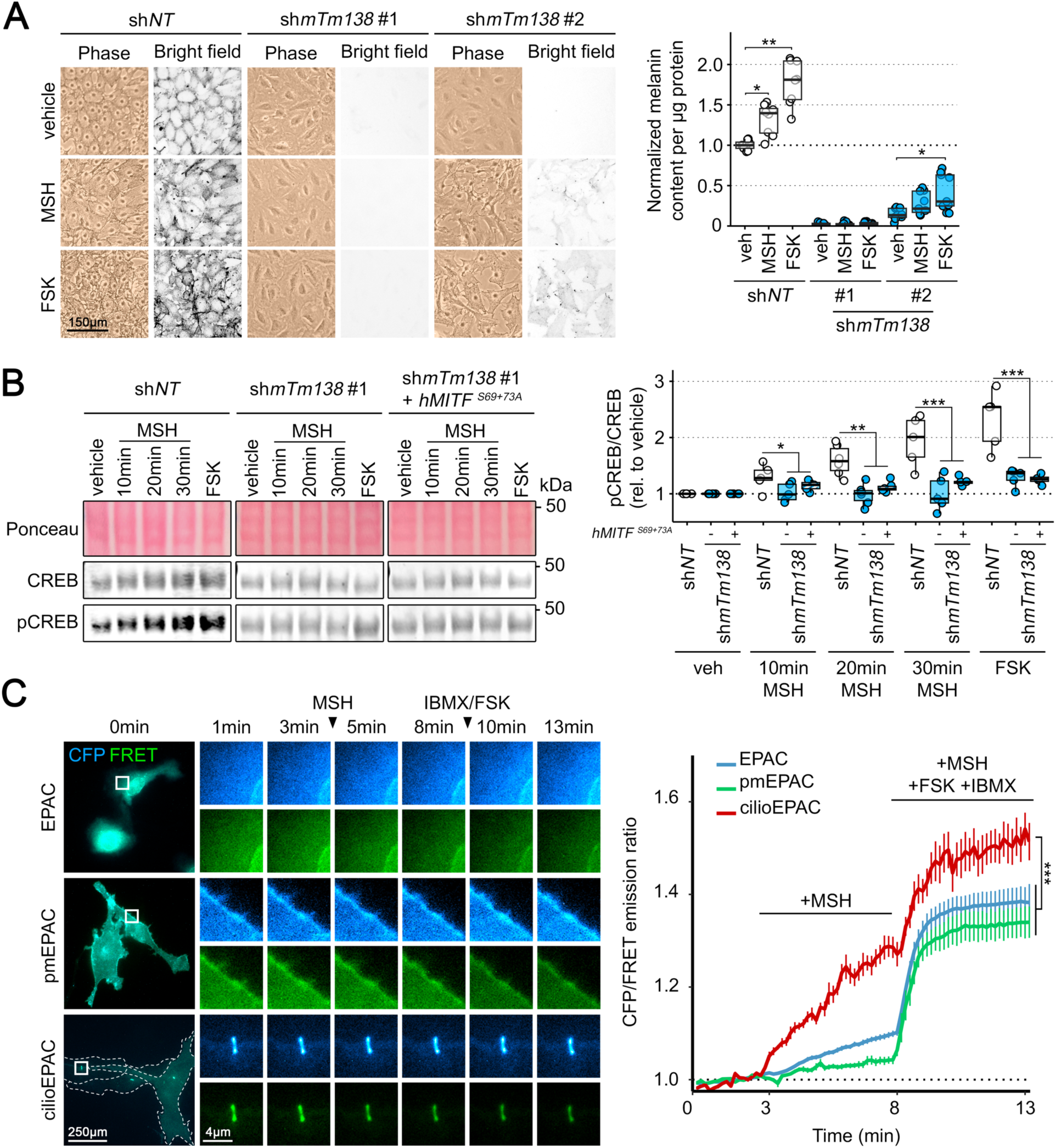
Primary cilia support MSH-induced pigmentation, CREB phosphorylation and cAMP production. (**A** to **B)** Melan-*Ink4a* cells stably expressing sh*NT*, sh*mTm138* #1, sh*mTm138*#2, or both sh*mTm138* #1 and *hMITF^S69+73A^* were serum-starved for 72 h. (A) Cells were then grown for three additional days in serum-containing medium. During and after starvation, cells were treated daily with either 0.6 µM Forskolin (FSK), 3 pM NDP-MSH (MSH), or a vehicle before imaging or harvesting. Left, representative images of phase and bright field contrast microscopy. Right, quantitative melanin content assay across 6 experiments. Values are normalized by melanin content of sh*NT*-expressing cells treated with vehicle. (B) After starvation, cells were stimulated with either 3 pM MSH for 10, 20 or 30 min, 0.6 µM FSK for 30 min, or a vehicle, and total CREB and pCREB expression was analyzed by immunoblotting. Left, representative immunoblots. Numbers to right show positions of molecular weight marker in kDa. Right, quantification of pCREB/CREB ratios relative to vehicle across 6 experiments. **(C)** Melan-*Ink4a* cells were transduced with retroviruses to express either the cytosolic EPAC, a plasma membrane resident EPAC (pmEPAC), or a primary cilium-targeted EPAC (cilioEPAC) and mCherry-ARL13B to mark the primary cilium (not shown). Cells were selected with blasticidin for 3 days, and serum-starved for 3 additional days before live cell fluorescence microscopy imaging at baseline for 3 min, after 0.5 nM MSH stimulation for an additional 5 min, and after 200 µM 3-Isobutyl-1-methylxanthine (IBMX)/20µM FSK for 5 min. Left, representative images of whole cells per condition at baseline and of boxed regions over time. Outlines of cells transduced with cilioEPAC are highlighted with white dotted lines. Right, graph of CFP/FRET emission ratios over time across 6 to 9 cells per condition. Statistics: Data were analyzed using one-way ANOVA and Tukey’s multiple comparison test (A, B), or Likelihood Ratio Chi-Square test (C). *, p<0.05; **, p<0.01; ***, p<0.001.

To definitively test whether endogenous MC1R signaling occurs from primary cilia in pigmented, non-melanoma cells, we next assessed the localized production of cAMP following treatment with NDP-α-MSH. Wild-type melan-*Ink4a* were transiently transduced with recombinant retroviruses to express EPAC-S^H187^ – a cytosolic cAMP biosensor that reduces its YFP/CFP FRET signal when bound to cAMP (*51*) – either alone (EPAC), fused to the plasma membrane-targeting sequence from Lyn tyrosine kinase (pmEPAC), or anchored to the C-terminus of the ciliary transmembrane protein Arl13b (cilioEPAC). Cells were co-transduced with retroviruses to express Arl13b-mCherry (to mark the primary cilium in live cells) and either of the three EPAC variants, serum starved for 72h to induce ciliogenesis, and then individual ciliated transductants were imaged in real-time at baseline, after MSH treatment, and after subsequent addition of both FSK and phosphodiesterase inhibitor 3-Isobutyl-1-methylxanthine (IBMX) to record maximal cAMP release. Analyses of single cells (Fig. 3C, left) or aggregate data from multiple cells (Fig. 3C, right) revealed that cAMP release was quicker and stronger in response to MSH at the primary cilium compared to the cell plasma membrane or cytosol, demonstrating that MC1R signals to ADCY most efficiently from the primary cilium.

In addition to its roles in melanogenesis, MC1R regulates melanocyte survival, proliferation, and oxidative stress-induced DNA repair through cAMP-independent transactivation of the receptor tyrosine kinase cKIT (*52*, *53*). To test whether primary cilia also regulate non-pigmentary functions of MC1R, we assessed how *mTm138* depletion affects MSH-induced proliferation. Serum-starved melan-*Ink4a* cells stably expressing either sh*NT*, sh*mTm138* #1, sh*mTm138* #2, or sh*mTm138* #1/*hMITF^S69+73A^* (to restore MC1R expression levels without rescuing ciliation) were treated with MSH, and live cells were counted 6 days later. MSH-induced proliferation was significantly impaired in cells expressing sh*mTm138* #1 (1.5-fold decrease), and sh*mTm138* #2 (1.2-fold decrease), but not sh*mTm138* #1/*hMITF^S69+73A^*compared to control cells (Fig. S3A). These data indicate that reduced proliferation in response to MSH is due to decreased *Mc1r* mRNA expression but not impaired ciliogenesis in *mTm138*-depleted cells. Consistent with these results, immunoblotting analysis showed that MSH stimulation induced the phosphorylation of ERK, a downstream target of cKIT, in melanocytes expressing either sh*NT*, sh*mTm138* #2, or sh*mTm138* #1/*hMITF^S69+73A^* (Fig. S3B). Together, these data demonstrate that MC1R-dependent pigmentation is specifically limited to a ciliary-resident cohort of endogenous MC1R, causing downstream elevation of cAMP, pCREB, and melanin content, but that MC1R-dependent proliferation and survival can be stimulated by a cohort of endogenous MC1R expressed outside of primary cilia in mouse melanocytes.

### JS but not BBS genes regulate TYRP1 localization

Because (i) *hMITF^S69+73A^* expression restores pigment gene expression in melanocytes depleted of both BBS- and JS-related genes, but only fully rescues pigmentation in BBS gene depleted cells (Fig. 2D), (ii) melanocytes depleted of the JS gene *mTm216* have reduced pigmentation (Fig. S1F) despite no obvious effect on ciliogenesis (Fig. S1E) or pigment gene expression (Fig. 2A), and (iii) some genes have been associated with both ciliogenesis and endosomal membrane trafficking (*54*, *55*), we assessed whether extra-ciliary functions of JS genes could influence melanosome protein localization. We first examined whether the steady-state localization of the melanosomal protein TYRP1 was affected by silencing JS or BBS genes in melan-*Ink4a* cells. In wild type melanocytes, TYRP1 localizes primarily to the limiting membrane of pigmented melanosomes found throughout the cell body, appearing in a ‘ring’-like pattern due to the light-absorbing properties of melanin (*38*, *56–58*). The same expression pattern was observed by deconvolution IFM (dIFM) and quantitative image analysis in cells either expressing sh*NT*, or depleted of BBS genes *mSdccag8* and *mIft172* (Fig. 4A). TYRP1 signal intensity in BBS gene-depleted cells was slightly reduced compared to control cells (Fig. 4D) as expected from weaker *Tyrp1* mRNA expression (Fig. 2A). However, in cells depleted of JS genes *mTm138*, *mTm216*, and *mInpp5e*, the low level of TYRP1 signal intensity accumulated in small punctate structures in the perinuclear region (Fig. 4A and B). This unusual distribution could not be explained by low mRNA expression levels, as TYRP1 distribution in *mSdccag8*- and *mIft172*-depleted cells was normal. Intriguingly, cell treatment for 3h with the protein synthesis inhibitor cycloheximide (CHX) significantly reduced TYRP1 expression in *mTm138*-depleted cells, but not control cells (Fig. S4A), suggesting that TYRP1 is rapidly degraded upon *mTm138* silencing.

**Figure 4.**
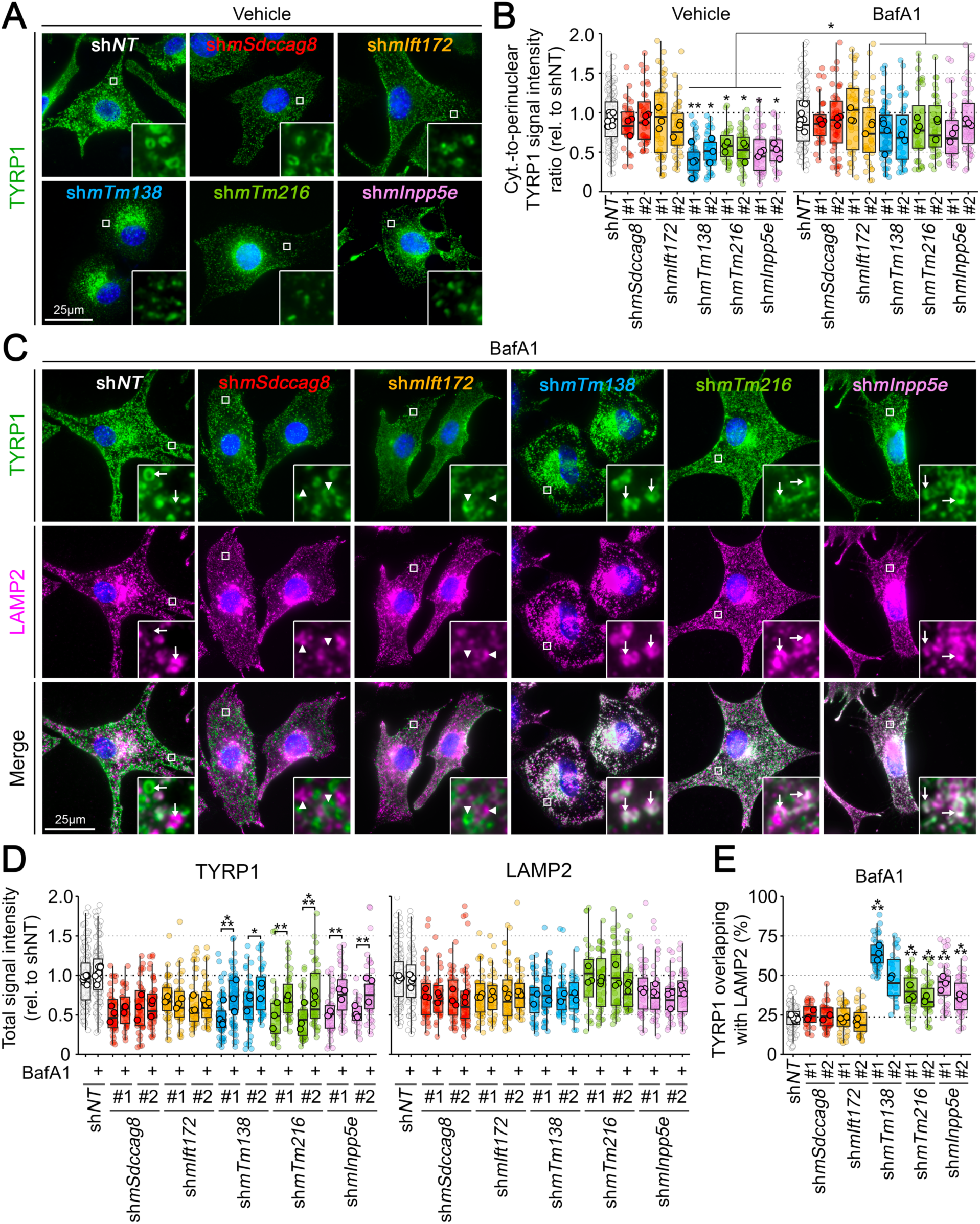
Depletion of JS but not BBS gene products mistargets TYRP1 to lysosomes for degradation. Melan-*Ink4a* melanocytes stably expressing sh*NT* or either of two shRNAs to *mTm138* (sh*mTm138*), *mTmem216* (sh*mTm216*), *mInpp5e* (sh*mInpp5e*), *mSdccag8* (sh*mSdccag8*) or *mIft172* (sh*mIft172*) were treated with vehicle or 25 nM bafilomycin A1 (BafA1) for 3h, fixed, and analyzed by dIFM for TYRP1 and LAMP2. **(A)** Representative dIFM images of cells treated with vehicle. **(B)** Cytoplasmic (cyt.)-to-perinuclear TYRP1 signal intensity ratio was quantified in 29-165 cells per condition across 3-8 experiments. **(C)** Representative dIFM images of cells treated with BafA1. Arrowheads indicate lack of overlap and arrows show overlap between TYRP1 and LAMP2. **(D)** Total signal intensities for TYRP1 and LAMP2 were quantified in 29-165 cells per condition across 3-8 experiments. **(E)** Percentage of TYRP1 area overlapping with LAMP2 total area in cells treated with BafA1 was quantified in 31 to 104 melanocytes per condition across 2-5 experiments. In (A) and (C), insets are 7-fold magnifications of boxed regions. Statistics: Data were analyzed using one-way ANOVA and Tukey’s multiple comparison test. *, p<0.05; **, p<0.01; ***, p<0.001.

To test if lysosomal degradation contributed to low TYRP1 protein expression in JS gene-depleted cells, shRNA transductants were analyzed by dIFM 3-5h after treatment with the vacuolar ATPase inhibitor bafilomycin A1 (BafA1). Whereas BafA1 had little effect on TYRP1 signal intensity or distribution in cells expressing sh*NT*, sh*mSdccag8*, or sh*mIft172*, it significantly restored TYRP1 peripheral signal in cells expressing sh*mTm138*, sh*mTm216*, or sh*mInpp5e* compared to vehicle-treated cells (Fig. 4B, C, and D). Moreover, the restored signal by dIFM overlapped with that of LAMP2, a membrane protein enriched on late endosomes and lysosomes (Fig. 4C and E), substantiating lysosomal degradation of TYRP1 in these cells. This effect was likely specific for melanosomal proteins like TYRP1, as BafA1 treatment did not affect the intensity or distribution of LAMP2 labelling in any of the cells analyzed (Fig. 4D). Together, these data reveal that while most ciliopathy proteins regulate pigment gene transcription, TMEM138 and other JS-related gene products are specifically required for proper targeting of at least one melanosomal protein.

### TMEM138 promotes melanosomal protein localization independently of MITF expression

Most melanogenic protein cargoes are delivered to maturing melanosomes using a well-known trafficking machinery (*38*, *46*, *59–62*) composed in part of potential transcriptional targets of MITF (*63*). Since defective ciliogenesis reduces expression of *MITF* and its target genes (Fig. 2A), we next tested whether TYRP1 mislocalization in JS gene-depleted cells could be rescued by *hMITF^S69+73A^* expression. Despite normalized *Tyrp1* mRNA levels relative to cells expressing sh*mTm138* #1 alone, TYRP1 was still mislocalized and degraded in sh*mTm138 #1*/*hMITF^S69+73A^* melanocytes as evidenced by dIFM and quantitative image analysis (Fig. 5A), and confirmed by immunoblotting (Fig. 5B). These data demonstrate that TYRP1 mislocalization and degradation in *Tm138*-depleted cells is independent of *MITF* expression and thus likely unrelated to defective ciliogenesis.

**Figure 5.**
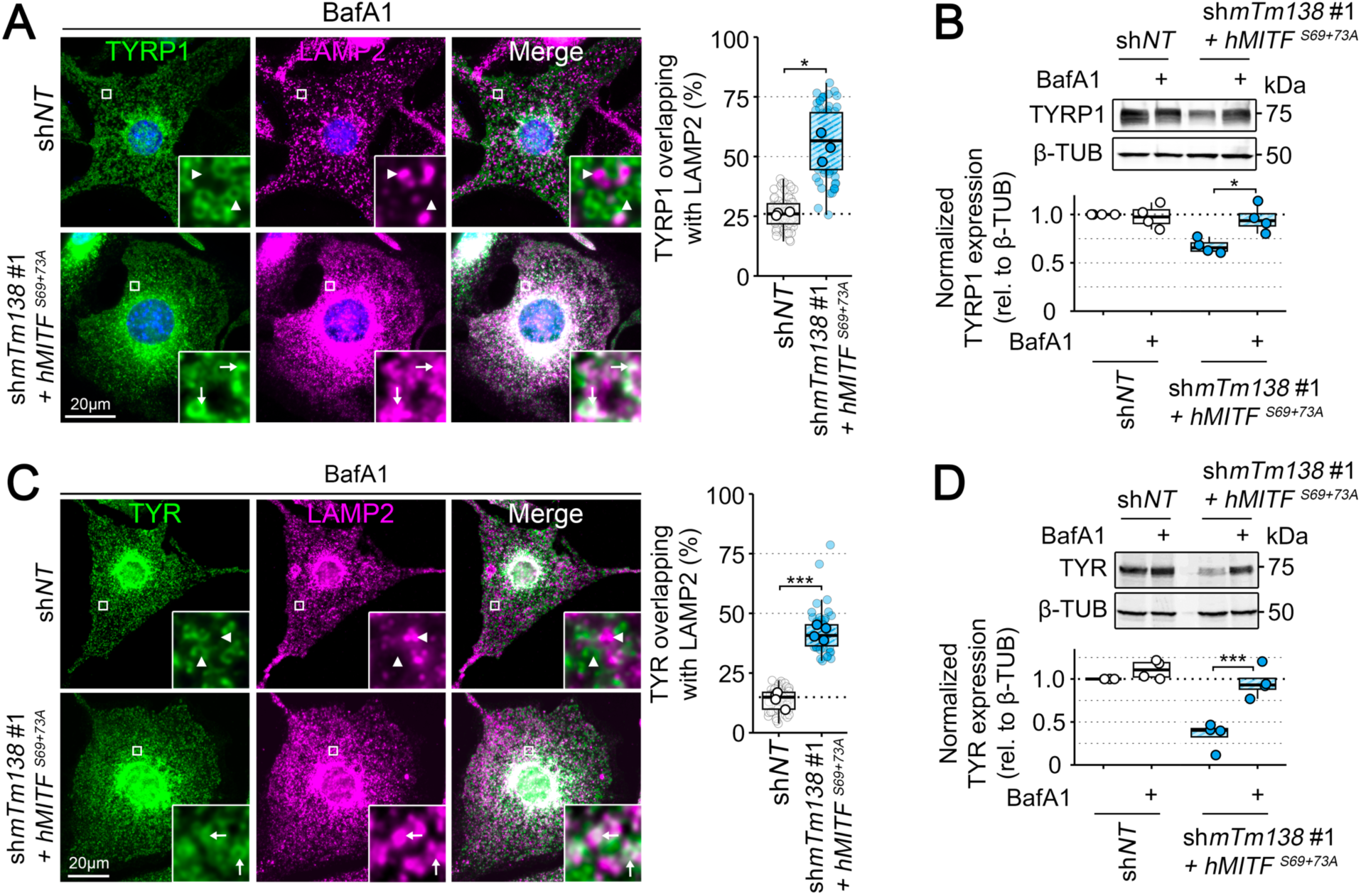
Active MITF expression does not rescue melanogenic cargo mislocalization in TMEM138-depleted melanocytes. Melan-*Ink4a* melanocytes stably expressing sh*NT* or both sh*mTm138* #1 and *hMITF^S69+73A^* were treated with vehicle or 25 nM BafA1 for 3 to 5 h, and then either fixed for dIFM imaging or harvested for immunoblotting. **(A** and **C)** Left, representative dIFM images of transduced cells labeled for TYRP1 and LAMP2 (A) or TYR and LAMP2 (C). Insets represent 7-fold magnifications of boxed regions; arrowheads indicate lack of overlap and arrows show overlap between both labels. Right, percentage of TYRP1 (A) or TYR (C) area overlapping with LAMP2 was quantified in 54-71 cells per condition across 3-4 experiments. **(B** and **D)** Immunoblotting of TYRP1 (B) or TYR (D) expression relative to β-Tubulin (β-TUB). Top, representative immunoblot; positions of nearby molecular weight markers are indicated to the right in kDa. Bottom, quantification of TYRP1 (B) or TYR (D) content relative to β-TUB across 4 experiments. Values are normalized by protein expression of cells expressing sh*NT* treated with vehicle. Statistics: Data were analyzed using paired t-tests (A, C) or one-way ANOVA and Tukey’s multiple comparison test (B, D). *, p<0.05; ***, p<0.001.

The increased melanogenic mRNA expression in *Tm138*-depleted cells expressing *hMITF^S69+73A^* allowed us to test whether TMEM138-dependent control over post-transcriptional trafficking of TYRP1 extended to other endogenous melanosomal protein cargoes. Like TYRP1, TYR was mislocalized and degraded in a BafA1-sensitive manner in sh*mTm138 #1*/*hMITF^S69+73A^*cells relative to sh*NT* melanocytes, as evidenced by both dIFM and quantitative image analysis (Fig. 5C), and by immunoblotting (Fig. 5D). These data support the conclusion that TMEM138 and likely other JS proteins play a moonlighting role, independent of their role in ciliary-dependent regulation of melanogenic signaling, in modulating melanosome maturation by ensuring proper cargo delivery.

### TMEM138 is required for the segregation of the melanosomal and endolysosomal systems

We initially interpreted the data above as evidence that TMEM138 functions to promote vesicular trafficking of melanogenic cargoes to immature melanosomes. Surprisingly, however, the early stage melanosome protein PMEL – a functional amyloid protein that segregates from the endosomal system prior to the delivery of other melanogenic protein cargoes (*57*, *64*) – also co-segregated with lysosomal markers by dIFM in sh*mTm138 #1*/*hMITF^S69+73A^-*but not sh*NT*-expressing melanocytes (Fig. 6A). Moreover, while immunoblotting showed that both sh*NT*- and sh*mTm138* #1/*hMITF^S69+73A^*-transduced cells expressed similar levels of immature unprocessed forms of PMEL (detected by Pep13h immunoblotting (*64*)), only lysates of sh*NT* cells expressed high levels of detergent-insoluble amyloidogenic PMEL fragments (detected by I51 (*65*); Fig. 6B). These data indicate that TMEM138 is required for proper pH-dependent biogenesis of PMEL amyloid in early stage melanosomes prior to melanogenic cargo delivery. Finally, while LAMP2 was completely segregated from melanin pigments in sh*NT* melanocytes, most pigment granules were surrounded by LAMP2 in sh*mTm138* #1/*hMITF^S69+73A^*cells (as shown by combined use of dIFM and bright field microscopy; Fig. 6C), suggesting that melanin was either synthesized in or engulfed by late endosomes and/ or lysosomes in these cells. These data are less consistent with a requirement for TMEM138 in the formation or delivery of melanogenic cargo transport carriers, and more reminiscent of defects in GPR143 or PIKFyve that promote the segregation of early stage melanosomes from maturing endosomes (*66–68*).

**Figure 6.**
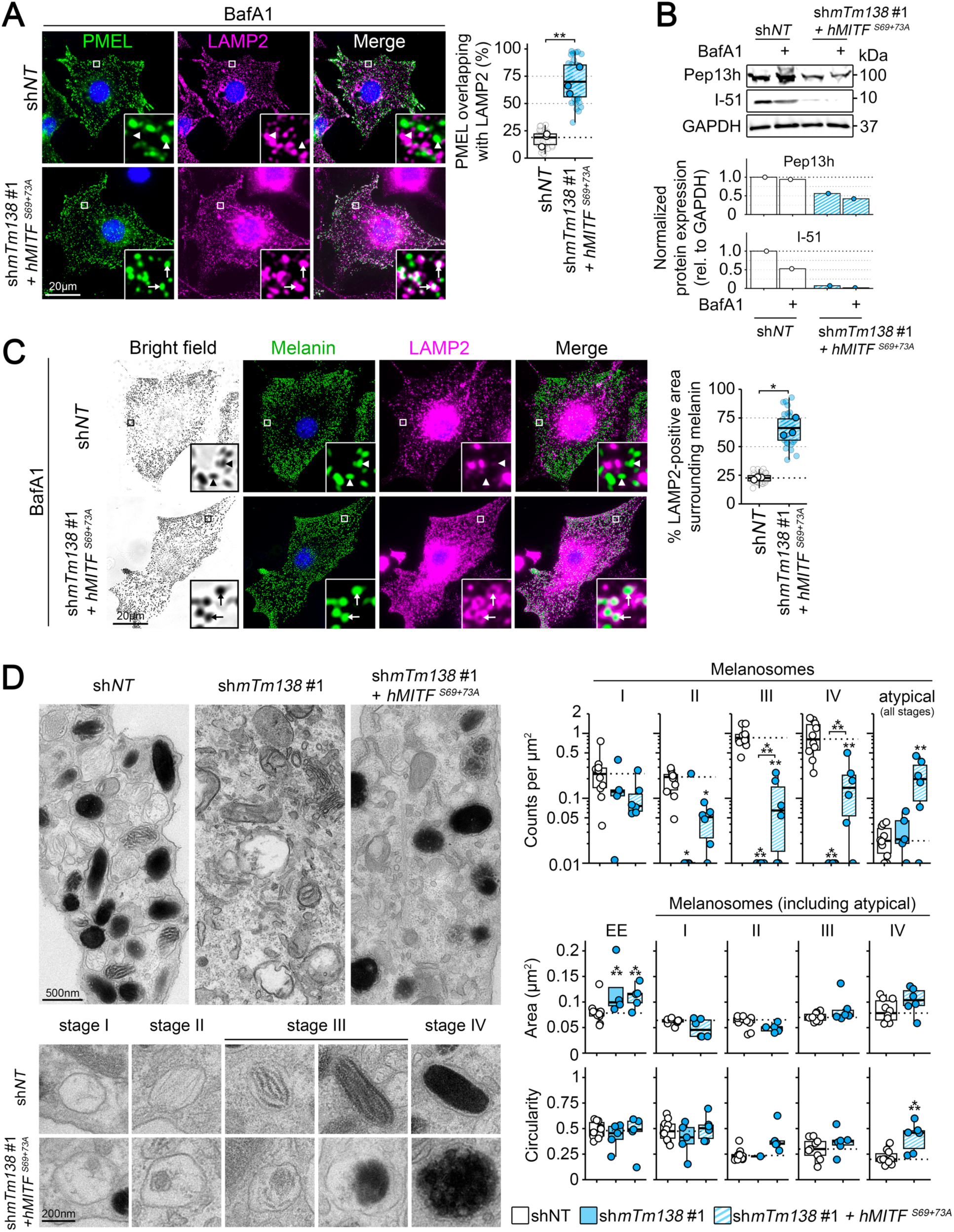
TMEM138 silencing disrupts early stages of melanosome biogenesis. Melan-*Ink4a* melanocytes stably expressing sh*NT*, sh*mTm138* #1 alone, or both sh*mTm138* #1 and *hMITF^S69+73A^* were treated with vehicle or 25 nM BafA1 for 3 to 5 h, and then fixed for immunoblotting or dIFM **(A** to **C)**, or untreated and fixed for TEM imaging **(D)**. (A) Left, representative dIFM images of transduced cells labeled for PMEL and LAMP2. Right, percentage of PMEL area overlapping with LAMP2 were quantified in 38 cells per condition across 3 experiments. (B) Top, representative immunoblots for PMEL unprocessed precursor protein (with Pep13h antibody) and core amyloid fragment (with I-51 antibody) relative to GAPDH; positions of nearby molecular weight markers are indicated to the right in kDa. Bottom, quantification of Pep13h (top) and I-51 (bottom) content relative to GAPDH in one experiment. Note that BafA1 has little effect on Pep13h immunoblotting as the unprocessed PMEL precursor protein is not expressed post-Golgi, but reduces I-51 immunoblotting as it blocks pH-dependent cleavage of PMEL required for amyloidogenic fiber formation (*64*, *87*, *88*). (C) Left, representative dIFM images of transduced cells labeled for LAMP2. Melanin is pseudocolored from bright field images to facilitate visualization of LAMP2. Right, percentage of LAMP2-positive area surrounding melanin was assessed in 35-41 cells per condition across 3 experiments. (A and C) Insets represent 7-fold magnifications of boxed regions; arrows show overlap and arrowheads indicate lack of overlap between both labels. (D) Left, representative TEM images of transduced cells showing cell bodies (top), and melanosome maturation stages (bottom). Right, organelle counts, areas, and circularities for all melanosome maturation stages and early endosomes were quantified in 5-10 cells per condition in a single experiment. Note that organelle counts are displayed on a log scale. Statistics: Data were analyzed using paired t-tests (A, C) or one-way ANOVA and Tukey’s multiple comparison test (D). *, p<0.05; **, p<0.01; ***, p<0.001.

To test this hypothesis, we analyzed melanocytes expressing sh*NT*, sh*mTm138* #1, or sh*mTm138* #1/*hMITF^S69+73A^* by thin section transmission electron microscopy (TEM; Fig. 6D). Cells expressing sh*NT* harbored numerous melanosomes of all stages with morphology typical of that commonly observed in wild-type melanocytes (*69*), including round vacuolar endosomes/ stage I melanosomes with intralumenal vesicles and disorganized PMEL fibrils, elliptical and striated stage II and III melanosomes containing, respectively, no or varying levels of melanin along the fibrous sheets of PMEL, and elliptical stage IV melanosomes completely filled with melanin. As expected, cells expressing sh*mTm138* #1 alone largely lacked any typical melanosome structures, consistent with the impaired expression of most melanogenic genes. Moreover, both sh*mTm138* #1 and sh*mTm138* #1/*hMITF^S69+73A^* cells displayed enlarged early endosomes compared to control cells. Interestingly, compared to sh*NT* melanocytes, sh*mTm138* #1/*hMITF^S69+73A^*cells had higher numbers of enlarged and rounded melanosomes with atypical maturation stages in which melanin deposition along PMEL sheets was nearly absent. These atypical melanosomes lacked the characteristic double membrane of autophagosomes, indicating that they were not the result of melanophagy, and often contained melanin in irregular clumps and membrane whorls reminiscent of lysosomes. Altogether, these data support that TMEM138, and perhaps other JS proteins, play a critical MITF-independent post-transcriptional role in the segregation of the melanosomal and endolyosomal lineages from early endosomes within melanocytes.

## Discussion

Our data show that ciliopathy proteins support basal pigmentation in epidermal melanocytes via a dual mechanism involving (i) endogenous ciliary GPCR-driven signal transduction regulated by most ciliopathy proteins to upregulate melanogenic gene transcription, and (ii) an extraciliary function in endolysosomal membrane dynamics specifically controlled by JS-related ciliopathy proteins to sustain melanosome biogenesis. Our observations link a wide range of ciliopathy genes to LRO biogenesis and unveil a yet undocumented role for JS genes in cilium-independent cell type-specific membrane dynamics required for LRO maturation. Our findings explain why genetic variation controlling the expression of *TMEM138* and other ciliopathy genes impacts skin pigmentation, suggest that hypopigmentation may be an as yet unrecognized consequence of JS, and point to extraciliary roles of JS proteins in cell type-specific membrane trafficking as a potential reason why JS is specifically associated with severe neuronal impairment.

Our GWAS analyses revealed a link between variations in human pigmentation and regulatory variants for several JS genes including *TMEM138*, but not for genes deficient in other ciliopathies such as BBS (*12*, *18*). Given our observations that most ciliopathy genes impact MC1R signaling but only JS genes also affect melanosome biogenesis, we propose that defects in melanogenic signaling from primary cilia might be more easily compensated at the skin tissue level than defects in intracellular membrane dynamics regulating melanosome biogenesis at the subcellular level. Consistent with this idea, we found that *TMEM138* silencing did not affect pigmentation as severely in reconstructed skin as in melanocyte monocultures, suggesting that pigment transfer to keratinocytes and keratinocyte- and fibroblast-induced stimulation of melanogenesis (*70–72*) do not require ciliary function in melanocytes. Dysfunction of the MC1R–MITF signaling axis in unciliated melanocytes might be overcome in the skin epidermis by stimulation of other cilium-independent melanogenic pathways, such as those driven by the Endothelin type B or cKIT receptors (*2*). These compensatory mechanisms may also explain why reduced skin pigmentation has not been described generally in ciliopathy patients. Nevertheless, our unquantified clinical observations suggest that many JS patients have fairer skin and hair than their unaffected siblings and parents.

Our shRNA screen showed that at least one shRNA to most tested ciliopathy genes decreased melanogenesis in mouse melanocytes, an effect possibly underestimated as we do not know the knockdown efficiency of all tested shRNAs. Of the one third of genes for which no shRNA reduced pigment content significantly, most are linked to diseases that disrupt cell type- or tissue-specific processes unrelated to pigment cells (e.g. *Kif3a*, atopic dermatitis; *Glis3*, diabetes mellitus; *Ccdc114*, primary ciliary dyskinesia; *Wdr60*, short-rib thoracic dysplasia; etc.). *Tmem138* depletion had among the strongest effects on pigmentation, which might be explained if its protein product is the limiting factor of a putative protein complex with other JS proteins including *Ahi1* and *Tmem231* (*25*). Interestingly, unlike observations in human kidney cells (*24*), silencing of *Tmem216* in mouse melanocytes by ∼ 80% did not impact ciliogenesis, suggesting either that a low level of *Tmem216* is sufficient to support ciliogenesis in melanocytes or that there exist cell type-specific differences in required ciliogenesis components.

Although a link between ciliary function and MC1R signaling has been previously reported under very specific activating conditions combining both UV light and MSH in unpigmented human melanoma cell lines (*50*), our study shows in non-transformed melanocytes that primary cilia regulate constitutive pigmentation and are required to activate endogenous, basal MC1R-induced melanogenic signaling. This requirement might reflect any or all of three possible mechanisms: (i) as MC1R is endogenously expressed at very low copy numbers (*73*), its enrichment within primary cilia likely strengthens signaling ability, as previously proposed for the related MC4R in neurons (*74*); (ii) ciliary enrichment would promote MC1R interaction with specific ADCY isoforms, such as ADCY-3 (*75*), which might be required for transmitting MC1R signaling to cAMP production; and (iii) inhibitory GPCRs expressed at the plasma membrane but excluded from the primary cilium, such as OPN3 (*76*), might inhibit MC1R-induced cAMP production elsewhere at the plasma membrane in primary human melanocytes, making the ciliary membrane a privileged space for MC1R signaling. Intriguingly, our experiments indicate that MC1R-induced proliferation is independent of primary cilium function, suggesting that primary cilia have a spatial control over separate axes of GPCR signaling.

TMEM138, and likely other JS proteins, seem to be necessary to segregate the endosomal and melanosomal lineages from early sorting endosomes (*57*). Segregation of these lineages is still a mysterious aspect in the biogenesis of melanosomes and other LROs, for which only a few genes have been proposed to participate (*77*). One of these genes is GPR143, a GPCR that localizes at steady state throughout the endolysosomal and melanosomal systems and for which a functional defect is associated with ocular albinism type I (*66*, *78*, *79*). Thus, TMEM138 might promote endosomal “splitting” by chaperoning GPR143 to the endosomal system, as it has been proposed to chaperone other GPCRs to the primary cilium like Rhodopsin in ocular photoreceptors (*25*), and perhaps MC1R in melanocytes. Alternatively, as a predicted tetraspan protein, TMEM138 might function by promoting formation of membrane microdomains on endosomes to permit segregation of stage II melanosomes from maturing late endosomes, similar to the role of CD63 in promoting inward budding for formation of intralumenal membranes (*80*). We predict that a cohort of TMEM138 and other JS proteins localize to early endosomes to facilitate these functions, but thus far we have been unable to visualize functionally tagged TMEM138 in melanocytes.

LROs, including melanosomes in melanocytes, originate from cell type-specific adaptations of the endolysosomal system (*81*). Using melanocytes as a tractable model system to study TMEM138 and other JS proteins, we uncovered unexpected extraciliary roles for these proteins in LRO biogenesis that would have unlikely been observed otherwise in cell types such as fibroblasts or HeLa cells. This novel association between JS proteins and LRO biogenesis might underlie several pathological features of JS, especially in the skin and brain. Skin lamellar bodies are LROs produced exclusively in highly differentiated skin keratinocytes and are essential for skin barrier formation (*82*). Defects in lamellar body biogenesis might explain why skin rashes, allergies, and recurrent infections have been reported in a subset of JS patients (*29*). Importantly, synaptic vesicle biogenesis in brain neurons also shares features with LRO biogenesis (*83*), and defects in specific components of the LRO machinery compromise synaptic vesicle maturation (*84*), transport (*85*), and release (*86*). Hence, defects in synaptic vesicle biogenesis might at least partly account for significant brain abnormalities found in JS patients compared to other ciliopathy patients.

## Acknowledgments

We thank Elena Oancea (Brown University; Providence, RI, USA), and Graça Raposo (Institut Curie, Paris, France) for their input into the project, Michael Hast (University of Pennsylvania; Philadelphia, PA, USA) for designing and fabricating custom pedestals used for the biofabrication of reconstructed skin, Duarte Barral (NOVA University; Lisbon, Portugal) for providing the pcDNA-ENTR-ARL13B-mCherry construct, Shuixing Li (Children’s Hospital of Philadelphia Research Institute; Philadelphia, PA, USA) for helping with cloning, Alex Simon (University of Pennsylvania; Philadelphia, PA, USA) for help with FRET imaging, Chris Marshall (Skin Biology and Disease Resource-based Center (SBDRC) of University of Pennsylvania; Philadelphia, PA, USA) for providing all the human skin samples, Inna Martynyuk and Biao Zuo (Electron Microscopy Resource Laboratory of University of Pennsylvania; Philadelphia, PA, USA) for sample preparation and imaging by TEM, and Roseanne Davila-Rivera, Rachel Tocci, Matias Schmukler, Rachel Welles, Christopher Davis, and Alexandre Lacourtiade for technical assistance, feedback, and support.

## Funding

This work was supported by:

National Institutes of Health grant R01EY015625 (M.S.M., G.R.)

National Institutes of Health grant R01AR076241 (M.S.M., S.A.T., E.O.)

National Institutes of Health grant 1R01AR085585-01 (S.A.T.)

National Institutes of Health grant 1R35GM161902-01 (S.A.T.)

LEO foundation grant LF-OC-25-002553 (M.S.M.)

Penn Skin Biology and Diseases Resource-based Center grant P30-AR069589 (B.M.)

Children’s Hospital of Philadelphia Research Institute, Bridge-to-faculty program fellowship GRT-00006135 (B.M.)

The content is solely the responsibility of the authors and does not necessarily represent the official views of the National Institutes of Health.

## Author contributions

Conceptualization: B.M., E.O., G.R., M.S.M.

Methodology: B.M., D.C.H., R.J.L., G.R., C.R.G., M.S.M.

Investigation: B.M., D.C.H., S.F., J.Z., C.E.S., Y.F.

Visualization: B.M.

Funding acquisition: B.M., S.A.T., E.O., G.R., M.S.M.

Project administration: B.M., M.S.M.

Supervision: B.M., M.S.M.

Writing – original draft: B.M., M.S.M.

Writing – review & editing: B.M., D.C.H., S.F., C.E.S., Y.F., S.A.T., E.O., G.R., C.R.G., R.J.L., M.S.M.

## Competing interests

Authors declare that they have no competing interests.

## Data, code, and materials availability

The data and detailed procedures needed to support the conclusions, and ensure reproducibility of this work are available within this manuscript and supplementary materials. Custom-made code for data analysis and other materials (e.g., plasmids, cell lines, antibodies) generated in lab will be made available upon request.

## Notes

### Competing Interest Statement

The authors have declared no competing interest.

## References and Notes

1. M. Brenner, V. J. Hearing, The protective role of melanin against UV damage in human skin. Photochem. Photobiol. 84, 539–549 (2008). doi:10.1111/j.1751-1097.2007.00226.x; pmid:18435612.

2. J. Y. Lin, D. E. Fisher, Melanocyte biology and skin pigmentation. Nature 445, 843–850 (2007). doi:10.1038/nature05660; pmid:17314970.

3. N. G. Jablonski, G. Chaplin, The evolution of human skin coloration. J. Hum. Evol. 39, 57–106 (2000). doi:10.1006/jhev.2000.0403; pmid:10896812.

4. N. Kobayashi, A. Nakagawa, T. Muramatsu, Y. Yamashina, T. Shirai, M. W. Hashimoto, Y. Ishigaki, T. Ohnishi, T. Mori, Supranuclear melanin caps reduce ultraviolet induced DNA photoproducts in human epidermis. Journal of Investigative Dermatology 110, 806­–810 (1998). doi:10.1046/j.1523-1747.1998.00178.x; pmid:9579550.

5. S. Benito-Martinez, L. Salavessa, G. Raposo, M. S. Marks, C. Delevoye, Melanin transfer and fate within keratinocytes in human skin pigmentation. Integr. Comp. Biol., 1–23 (2021). doi:10.1093/icb/icab094; pmid:34021340.

6. H. Moreiras, M. C. Seabra, D. C. Barral, Melanin transfer in the epidermis: The pursuit of skin pigmentation control mechanisms. Int. J. Mol. Sci. 22 (2021). doi:10.3390/ijms22094466; pmid:33923362.

7. R. E. Boissy, Melanosome transfer to and translocation in the keratinocyte. *Experimental Dermatology*, Supplement 12, 5–12 (2003). doi:10.1034/j.1600-0625.12.s2.1.x; pmid:14756517.

8. L. Montoliu, K. Grønskov, A. H. Wei, M. Martínez-García, A. Fernández, B. Arveiler, F. Morice-Picard, S. Riazuddin, T. Suzuki, Z. M. Ahmed, T. Rosenberg, W. Li, Increasing the complexity: New genes and new types of albinism. Pigment Cell Melanoma Res. 27, 11–18 (2014). doi:10.1111/pcmr.12167; pmid:24066960.

9. K. Grønskov, J. Ek, K. Brondum-Nielsen, Oculocutaneous albinism. Orphanet J. Rare Dis. 2 (2007). doi:10.1186/1750-1172-2-43; pmid:17980020.

10. D. J. Green, V. Michaud, E. Lasseaux, C. Plaisant, T. Fitzgerald, E. Birney, G. C. Black, B. Arveiler, P. I. Sergouniotis, The co-occurrence of genetic variants in the TYR and OCA2 genes confers susceptibility to albinism. Nature Communications 15 (2024). doi:10.1038/s41467-024-52763-y; pmid:39349469.

11. J. L. Rees, The melanocortin 1 receptor (MC1R): More than just red hair. Pigment Cell Res. 13, 135–140 (2000). doi:10.1034/j.1600-0749.2000.130303.x; pmid:10885670.

12. Y. Feng, N. Xie, F. Inoue, S. Fan, J. Saskin, C. Zhang, F. Zhang, M. E. B. Hansen, T. Nyambo, S. W. Mpoloka, G. G. Mokone, C. Fokunang, G. Belay, A. K. Njamnshi, M. S. Marks, E. Oancea, N. Ahituv, S. A. Tishkoff, Integrative functional genomic analyses identify genetic variants influencing skin pigmentation in Africans. Nat. Genet. 56, 258–272 (2024). doi:10.1038/s41588-023-01626-1; pmid:38200130.

13. K. Batai, Z. Cui, A. Arora, E. Shah-Williams, W. Hernandez, M. Ruden, C. M. P. Hollowell, S. E. Hooker, M. Bathina, A. B. Murphy, C. Bonilla, R. A. Kittles, Genetic loci associated with skin pigmentation in African Americans and their effects on Vitamin D deficiency. PLoS Genet. 17 (2021). doi:10.1371/journal.pgen.1009319; pmid:33600456.

14. F. Lona-Durazo, N. Hernandez-Pacheco, S. Fan, T. Zhang, J. Choi, M. A. Kovacs, S. K. Loftus, P. Le, M. Edwards, C. A. Fortes-Lima, C. Eng, S. Huntsman, D. Hu, E. J. Gómez-Cabezas, L. C. Marín-Padrón, J. Grauholm, O. Mors, E. G. Burchard, H. L. Norton, W. J. Pavan, K. M. Brown, S. Tishkoff, M. Pino-Yanes, S. Beleza, B. Marcheco-Teruel, E. J. Parra, Meta-analysis of GWA studies provides new insights on the genetic architecture of skin pigmentation in recently admixed populations. BMC Genet. 20 (2019). doi:10.1186/s12863-019-0765-5; pmid:31315583.

15. K. Adhikari, J. Mendoza-Revilla, A. Sohail, M. Fuentes-Guajardo, J. Lampert, J. C. Chacón-Duque, M. Hurtado, V. Villegas, V. Granja, V. Acuña-Alonzo, C. Jaramillo, W. Arias, R. B. Lozano, P. Everardo, J. Gómez-Valdés, H. Villamil-Ramírez, C. C. Silva de Cerqueira, T. Hunemeier, V. Ramallo, L. Schuler-Faccini, F. M. Salzano, R. Gonzalez-José, M. C. Bortolini, S. Canizales-Quinteros, C. Gallo, G. Poletti, G. Bedoya, F. Rothhammer, D. J. Tobin, M. Fumagalli, D. Balding, A. Ruiz-Linares, A GWAS in Latin Americans highlights the convergent evolution of lighter skin pigmentation in Eurasia. Nat. Commun. 10 (2019). doi:10.1038/s41467-018-08147-0; pmid:30664655.

16. I. Galván-Femenía, M. Obón-Santacana, D. Piñeyro, M. Guindo-Martinez, X. Duran, A. Carreras, R. Pluvinet, J. Velasco, L. Ramos, S. Aussó, J. M. Mercader, L. Puig, M. Perucho, D. Torrents, V. Moreno, L. Sumoy, R. De Cid, Multitrait genome association analysis identifies new susceptibility genes for human anthropometric variation in the GCAT cohort. J. Med. Genet. 55, 765–778 (2018). doi:10.1136/jmedgenet-2018-105437; pmid:30166351.

17. A. R. Martin, M. Lin, J. M. Granka, J. W. Myrick, X. Liu, A. Sockell, E. G. Atkinson, C. J. Werely, M. Möller, M. S. Sandhu, D. M. Kingsley, E. G. Hoal, X. Liu, M. J. Daly, M. W. Feldman, C. R. Gignoux, C. D. Bustamante, B. M. Henn, An Unexpectedly Complex Architecture for Skin Pigmentation in Africans. Cell 171, 1340–1353.e14 (2017). doi:10.1016/j.cell.2017.11.015; pmid:29195075.

18. N. G. Crawford, D. E. Kelly, M. E. B. Hansen, M. H. Beltrame, S. Fan, S. L. Bowman, E. Jewett, A. Ranciaro, S. Thompson, Y. Lo, S. P. Pfeifer, J. D. Jensen, M. C. Campbell, W. Beggs, F. Hormozdiari, S. W. Mpoloka, G. G. Mokone, T. Nyambo, D. W. Meskel, G. Belay, J. Haut, H. Rothschild, L. Zon, Y. Zhou, M. A. Kovacs, M. Xu, T. Zhang, K. Bishop, J. Sinclair, C. Rivas, E. Elliot, J. Choi, S. A. Li, B. Hicks, S. Burgess, C. Abnet, D. E. Watkins-Chow, E. Oceana, Y. S. Song, E. Eskin, K. M. Brown, M. S. Marks, S. K. Loftus, W. J. Pavan, M. Yeager, S. Chanock, S. A. Tishkoff, Loci associated with skin pigmentation identified in African populations. Science (1979). 358 (2017). doi:10.1126/science.aan8433; pmid:29025994.

19. F. Liu, M. Visser, D. L. Duffy, P. G. Hysi, L. C. Jacobs, O. Lao, K. Zhong, S. Walsh, L. Chaitanya, A. Wollstein, G. Zhu, G. W. Montgomery, A. K. Henders, M. Mangino, D. Glass, V. Bataille, R. A. Sturm, F. Rivadeneira, A. Hofman, W. F. J. van IJcken, A. G. Uitterlinden, R. J. T. S. Palstra, T. D. Spector, N. G. Martin, T. E. C. Nijsten, M. Kayser, Genetics of skin color variation in Europeans: genome-wide association studies with functional follow-up. Hum. Genet. 134, 823–835 (2015). doi:10.1007/s00439-015-1559-0; pmid:25963972.

20. S. Beleza, N. A. Johnson, S. I. Candille, D. M. Absher, M. A. Coram, J. Lopes, J. Campos, I. I. Araújo, T. M. Anderson, B. J. Vilhjálmsson, M. Nordborg, A. Correia e Silva, M. D. Shriver, J. Rocha, G. S. Barsh, H. Tang, Genetic Architecture of Skin and Eye Color in an African-European Admixed Population. PLoS Genet. 9 (2013). doi:10.1371/journal.pgen.1003372; pmid:23555287.

21. W. Branicki, U. Brudnik, J. Draus-Barini, T. Kupiec, A. Wojas-Pelc, Association of the SLC45A2 gene with physiological human hair colour variation. J. Hum. Genet. 53, 966–971 (2008). doi:10.1007/s10038-008-0338-3; pmid:18806926.

22. J. Han, P. Kraft, H. Nan, Q. Guo, C. Chen, A. Qureshi, S. E. Hankinson, F. B. Hu, D. L. Duffy, Z. Z. Zhen, N. G. Martin, G. W. Montgomery, N. K. Hayward, G. Thomas, R. N. Hoover, S. Chanock, D. J. Hunter, A genome-wide association study identifies novel alleles associated with hair color and skin pigmentation. PLoS Genet. 4 (2008). doi:10.1371/journal.pgen.1000074; pmid:18483556.

23. R. L. Lamason, M. A. P. K. Mohideen, J. R. Mest, A. C. Wong, H. L. Norton, M. C. Aros, M. J. Jurynec, X. Mao, V. R. Humphreville, J. E. Humbert, S. Sinha, J. L. Moore, P. Jagadeeswaran, W. Zhao, G. Ning, I. Makalowska, P. M. McKeigue, D. O’Donnell, R. Kittles, E. J. Parra, N. J. Mangini, D. J. Grunwald, M. D. Shriver, V. A. Canfield, K. C. Cheng, Genetics: SLC24A5, a putative cation exchanger, affects pigmentation in zebrafish and humans. Science (1979). 310, 1782–1786 (2005). doi:10.1126/science.1116238; pmid:16357253.

24. J. H. Lee, J. L. Silhavy, J. E. Lee, L. Al-Gazali, S. Thomas, E. E. Davis, S. L. Bielas, K. J. Hill, M. Iannicelli, F. Brancati, S. B. Gabriel, C. Russ, C. V. Logan, S. M. Sharif, C. P. Bennett, M. Abe, F. Hildebrandt, B. H. Diplas, T. Attié-Bitach, N. Katsanis, A. Rajab, R. Koul, L. Sztriha, E. R. Waters, S. Ferro-Novick, C. G. Woods, C. A. Johnson, E. M. Valente, M. S. Zaki, J. G. Gleeson, Evolutionarily assembled cis-regulatory module at a human ciliopathy locus. Science (1979). 335, 966–969 (2012). doi:10.1126/science.1213506; pmid:22282472.

25. D. Guo, J. Ru, L. Xie, M. Wu, Y. Su, S. Zhu, S. Xu, B. Zou, Y. Wei, X. Liu, Y. Liu, C. Liu, Tmem138 is localized to the connecting cilium essential for rhodopsin localization and outer segment biogenesis. Proc. Natl. Acad. Sci. U. S. A. 119 (2022). doi:10.1073/pnas.2109934119; pmid:35394880.

26. A. Radha Rama Devi, S. M. Naushad, L. Lingappa, Clinical and Molecular Diagnosis of Joubert Syndrome and Related Disorders. Pediatr. Neurol. 106, 43–49 (2020). doi:10.1016/j.pediatrneurol.2020.01.012; pmid:32139166.

27. F. Morelli, F. Toni, E. Saligari, F. D’Abrusco, V. Serpieri, E. Ballante, G. Ruberto, R. Borgatti, E. M. Valente, S. Signorini, Visual function in children with Joubert syndrome. Dev. Med. Child Neurol. 66, 379–388 (2024). doi:10.1111/dmcn.15732; pmid:37593819.

28. B. P. Brooks, W. M. Zein, A. H. Thompson, M. Mokhtarzadeh, D. A. Doherty, M. Parisi, I. A. Glass, M. C. Malicdan, T. Vilboux, M. Vemulapalli, J. C. Mullikin, W. A. Gahl, M. Gunay-Aygun, Joubert Syndrome: Ophthalmological Findings in Correlation with Genotype and Hepatorenal Disease in 99 Patients Prospectively Evaluated at a Single Center. Ophthalmology 125, 1937–1952 (2018). doi:10.1016/j.ophtha.2018.05.026; pmid:30055837.

29. Y. Dong, K. Zhang, H. Yao, T. Jia, J. Wang, D. Zhu, F. Xu, M. Cheng, S. Zhao, X. Shi, Clinical and genetic characteristics of 36 children with Joubert syndrome. Front. Pediatr. 11 (2023). doi:10.3389/fped.2023.1102639; pmid:37547106.

30. T. Aksu Uzunhan, B. Ertürk, K. Aydın, A. Ayaz, U. Altunoğlu, M. H. Yarar, A. Gezdirici, D. F. İçağasıoğlu, E. Gökpınar İli, B. Uyanık, M. Eser, Y. B. Kutbay, Y. Topçu, B. Kılıç, G. Bektaş, A. Arduç Akçay, B. Ekici, A. Chousein, Ş. Avcı, A. Yüksel, H. Kayserili, Clinical and genetic spectrum from a prototype of ciliopathy: Joubert syndrome. Clin. Neurol. Neurosurg. 224 (2023). doi:10.1016/j.clineuro.2022.107560; pmid:36580738.

31. T. Vilboux, D. A. Doherty, I. A. Glass, M. A. Parisi, I. G. Phelps, A. R. Cullinane, W. Zein, B. P. Brooks, T. Heller, A. Soldatos, N. L. Oden, D. Yildirimli, M. Vemulapalli, J. C. Mullikin, M. C. V. Malicdan, W. A. Gahl, M. Gunay-Aygun, Molecular genetic findings and clinical correlations in 100 patients with Joubert syndrome and related disorders prospectively evaluated at a single center. Genetics in Medicine 19, 875–882 (2017). doi:10.1038/gim.2016.204; pmid:28125082.

32. H. Y. Kroes, G. R. Monroe, B. Van Der Zwaag, K. J. Duran, C. G. De Kovel, M. J. Van Roosmalen, M. Harakalova, I. J. Nijman, W. P. Kloosterman, R. H. Giles, N. V. A. M. Knoers, G. Van Haaften, Joubert syndrome: Genotyping a Northern European patient cohort. European Journal of Human Genetics 24, 214–220 (2016). doi:10.1038/ejhg.2015.84; pmid:25920555.

33. R. Bachmann-Gagescu, J. C. Dempsey, I. G. Phelps, B. J. O’Roak, D. M. Knutzen, T. C. Rue, G. E. Ishak, C. R. Isabella, N. Gorden, J. Adkins, E. A. Boyle, N. de Lacy, D. O’Day, A. Alswaid, A. Radha Ramadevi, L. Lingappa, C. Lourenço, L. Martorell, Garcia-Cazorla, H. Ozyürek, G. Haliloglu, B. Tuysuz, M. Topçu, P. Chance, M. A. Parisi, I. A. Glass, J. Shendure, D. Doherty, Joubert syndrome: A model for untangling recessive disorders with extreme genetic heterogeneity. J. Med. Genet. 52, 514–522 (2015). doi:10.1136/jmedgenet-2015-103087; pmid:26092869.

34. D. P. Norris, D. T. Grimes, Mouse models of ciliopathies: the state of the art. Dis. Model. Mech. 5, 299–312 (2012). doi:10.1242/dmm.009340; pmid:22566558.

35. E. V. Sviderskaya, S. P. Hill, T. J. Evans-Whipp, L. Chin, S. J. Orlow, D. J. Easty, S. C. Cheong, D. Beach, R. A. DePinho, D. C. Bennett, p16Ink4a in melanocyte senescence and differentiation. J. Natl. Cancer Inst. 94, 446–454 (2002). doi:10.1093/jnci/94.6.446; pmid:11904317.

36. V. K. Bajpai, T. Swigut, J. Mohammed, S. Naqvi, M. Arreola, J. Tycko, T. C. Kim, J. K. Pritchard, M. C. Bassik, J. Wysocka, A genome-wide genetic screen uncovers determinants of human pigmentation. Science (1979). 381 (2023). doi:10.1126/science.ade6289; pmid:37561850.

37. J. M. Falcón-Pérez, M. Starcevic, R. Gautam, E. C. Dell’Angelica, BLOC-1, a novel complex containing the pallidin and muted proteins involved in the biogenesis of melanosomes and platelet-dense granules. Journal of Biological Chemistry 277, 28191­–28199 (2002). doi:10.1074/jbc.M204011200; pmid:12019270.

38. S. R. G. Setty, D. Tenza, S. T. Truschel, E. Chou, E. V. Sviderskaya, A. C. Theos, M. L. Lamoreux, S. M. Di Pietro, M. Starcevic, D. C. Bennett, E. C. Dell’Angelica, G. Raposo, M. S. Marks, BLOC-1 is required for cargo-specific sorting from vacuolar early endosomes toward lysosome-related organelles. Mol. Biol. Cell 18, 768–780 (2007). doi:10.1091/mbc.E06-12-1066; pmid:17182842.

39. R. Insolera, W. Shao, R. Airik, F. Hildebrandt, S. H. Shi, SDCCAG8 Regulates Pericentriolar Material Recruitment and Neuronal Migration in the Developing Cortex. Neuron 83, 805–822 (2014). doi:10.1016/j.neuron.2014.06.029; pmid:25088364.

40. V. Singla, J. F. Reiter, The primary cilium as the cell’s antenna: Signaling at a sensory organelle. Science (1979). 313, 629–633 (2006). doi:10.1126/science.1124534; pmid:16888132.

41. J. L. Tobin, P. L. Beales, The nonmotile ciliopathies. [Preprint] (2009). 10.1097/GIM.0b013e3181a02882.

42. C. R. Goding, H. Arnheiter, Mitf—the first 25 years. Genes Dev. 33, 983–1007 (2019). doi:10.1101/gad.324657.119; pmid:31123060.

43. J. Ballesteros-Álvarez, R. Dilshat, V. Fock, K. Möller, L. Karl, L. Larue, M. H. Ögmundsdóttir, E. Steingrímsson, MITF and TFEB cross-regulation in melanoma cells. PLoS One 15 (2020). doi:10.1371/journal.pone.0238546; pmid:32881934.

44. K. C. Ngeow, H. J. Friedrichsen, L. Li, Z. Zeng, S. Andrews, L. Volpon, H. Brunsdon, G. Berridge, S. Picaud, R. Fischer, R. Lisle, S. Knapp, P. Filippakopoulos, H. Knowles, E. Steingrímsson, K. L. B. Borden, E. E. Patton, C. R. Goding, BRAF/MAPK and GSK3 signaling converges to control MITF nuclear export. Proc. Natl. Acad. Sci. U. S. A. 115, E8668–E8677 (2018). doi:10.1073/pnas.1810498115; pmid:30150413.

45. P. S. Goff, S. Patel, D. C. Harper, T. Carter, M. S. Marks, E. V. Sviderskaya, Reprogramming of endolysosomes for melanogenesis in BLOC-1-deficient melanocytes. Current Biology 35, 3570–3586.e7 (2025). doi:10.1016/j.cub.2025.06.031.

46. A. Sitaram, M. K. Dennis, R. Chaudhuri, W. De Jesus-Rojas, D. Tenza, S. R. G. Setty, C. S. Wood, E. V. Sviderskaya, D. C. Bennett, G. Raposo, J. S. Bonifacino, M. S. Marks, Differential recognition of a dileucine-based sorting signal by AP-1 and AP-3 reveals a requirement for both BLOC-1 and AP-3 in delivery of OCA2 to melanosomes. Mol. Biol. Cell 23, 3178–3192 (2012). doi:10.1091/mbc.e11-06-0509; pmid:22718909.

47. S. Guida, G. Guida, C. R. Goding, MC1R Functions, Expression, and Implications for Targeted Therapy. Journal of Investigative Dermatology 142, 293–302.e1 (2022). doi:10.1016/j.jid.2021.06.018; pmid:34362555.

48. V. B. Swope, Z. A. Abdel-Malek, MC1R: Front and center in the bright side of dark eumelanin and DNA repair. MDPI AG [Preprint] (2018). 10.3390/ijms19092667.

49. E. M. Wolf Horrell, M. C. Boulanger, J. A. D’Orazio, Melanocortin 1 receptor: Structure, function, and regulation. Frontiers Media S.A. [Preprint] (2016). 10.3389/fgene.2016.00095.

50. X. Tian, H. Wang, S. Liu, W. Liu, K. Zhang, X. Gao, Q. Li, H. Zhao, L. Zhang, P. Liu, M. Liu, Y. Wang, X. Zhu, R. Cui, J. Zhou, Melanocortin 1 receptor mediates melanin production by interacting with the BBSome in primary cilia. PLoS Biol. 22, 1–26 (2024). doi:10.1371/journal.pbio.3002940.

51. J. Klarenbeek, J. Goedhart, A. Van Batenburg, D. Groenewald, K. Jalink, Fourth-generation Epac-based FRET sensors for cAMP feature exceptional brightness, photostability and dynamic range: Characterization of dedicated sensors for FLIM, for ratiometry and with high affinity. PLoS One 10 (2015). doi:10.1371/journal.pone.0122513; pmid:25875503.

52. M. Castejón-Griñán, C. Herraiz, C. Olivares, C. Jiménez-Cervantes, J. C. García-Borrón, CAMP-independent non-pigmentary actions of variant melanocortin 1 receptor: AKT-mediated activation of protective responses to oxidative DNA damage. Oncogene 37, 3631–3646 (2018). doi:10.1038/s41388-018-0216-1; pmid:29622793.

53. C. Herraiz, F. Journé, Z. Abdel-Malek, G. Ghanem, C. Jiménez-Cervantes, J. C. García-Borrón, Signaling from the human melanocortin 1 receptor to ERK1 and ERK2 mitogen-activated protein kinases involves transactivation of cKIT. Molecular Endocrinology 25, 138–156 (2011). doi:10.1210/me.2010-0217; pmid:21084381.

54. W. J. Monis, V. Faundez, G. J. Pazour, BLOC-1 is required for selective membrane protein trafficking from endosomes to primary cilia. Journal of Cell Biology 216, 2131–2150 (2017). doi:10.1083/jcb.201611138; pmid:28576874.

55. D. C. Barral, S. Garg, C. Casalou, G. F. M. Watts, J. L. Sandoval, J. S. Ramalho, V. W. Hsu, M. B. Brenner, Arl13b regulates endocytic recycling traffic. Proc. Natl. Acad. Sci. U. S. A. 109, 21354–21359 (2012). doi:10.1073/pnas.1218272110; pmid:23223633.

56. S. T. Truschel, S. Simoes, S. R. G. Setty, D. C. Harper, D. Tenza, P. C. Thomas, K. E. Herman, S. D. Sackett, D. C. Cowan, A. C. Theos, G. Raposo, M. S. Marks, ESCRT-I function is required for Tyrp1 transport from early endosomes to the melanosome limiting membrane. Traffic 10, 1318–1336 (2009). doi:10.1111/j.1600-0854.2009.00955.x; pmid:19624486.

57. G. Raposo, D. Tenza, D. M. Murphy, J. F. Berson, M. S. Marks, Distinct protein sorting and localization to premelanosomes, melanosomes, and lysosomes in pigmented melanocytic cells. Journal of Cell Biology 152, 809–823 (2001). doi:10.1083/jcb.152.4.809.

58. S. Vijayasaradhi, Y. Xu, B. Bouchard, A. N. Houghton, “Intracellular Sorting and Targeting of Melanosomal Membrane Proteins: Identification of Signals for Sorting of the Human Brown Locus Protein, GP75.”

59. S. L. Bowman, L. Le, Y. Zhu, D. C. Harper, A. Sitaram, A. C. Theos, E. V. Sviderskaya, D. C. Bennett, G. Raposo-Benedetti, D. J. Owen, M. K. Dennis, M. S. Marks, A BLOC-1–AP-3 super-complex sorts a cis-snare complex into endosome-derived tubular transport carriers. Journal of Cell Biology 220 (2021). doi:10.1083/jcb.202005173; pmid:33886957.

60. J. J. Bultema, A. L. Ambrosio, C. L. Burek, S. M. Di Pietro, BLOC-2, AP-3, and AP-1 proteins function in concert with Rab38 and Rab32 proteins to mediate protein trafficking to lysosome-related organelles. Journal of Biological Chemistry 287, 19550–19563 (2012). doi:10.1074/jbc.M112.351908; pmid:22511774.

61. C. Delevoye, I. Hurbain, D. Tenza, J. B. Sibarita, S. Uzan-Gafsou, H. Ohno, W. J. C. Geerts, A. J. Verkleij, J. Salamero, M. S. Marks, G. Raposo, AP-1 and KIF13A coordinate endosomal sorting and positioning during melanosome biogenesis. Journal of Cell Biology 187, 247–264 (2009). doi:10.1083/jcb.200907122; pmid:19841138.

62. S. M. Di Pietro, J. M. Falcón-Pérez, D. Tenza, S. R. G. Setty, M. S. Marks, G. Raposo, E. C. Dell’Angelica, BLOC-1 interacts with BLOC-2 and the AP-3 complex to facilitate protein trafficking on endosomes. Mol. Biol. Cell 17, 4027–4038 (2006). doi:10.1091/mbc.E06-05-0379; pmid:16837549.

63. T. Strub, S. Giuliano, T. Ye, C. Bonet, C. Keime, D. Kobi, S. Le Gras, M. Cormont, R. Ballotti, C. Bertolotto, I. Davidson, Essential role of microphthalmia transcription factor for DNA replication, mitosis and genomic stability in melanoma. Oncogene 30, 2319–2332 (2011). doi:10.1038/onc.2010.612; pmid:21258399.

64. J. F. Berson, D. C. Harper, D. Tenza, G. Raposo, M. S. Marks, Pmel17 Initiates Premelanosome Morphogenesis within Multivesicular Bodies. Mol. Biol. Cell 12, 3451–3464 (2001). doi:10.1091/mbc.12.11.3451.

65. J. S. Hee, S. M. Mitchell, X. Liu, R. M. Leonhardt, Melanosomal formation of PMEL core amyloid is driven by aromatic residues. Sci. Rep. 7 (2017). doi:10.1038/srep44064; pmid:28272432.

66. F. Giordano, C. Bonetti, E. M. Surace, V. Marigo, G. Raposo, The ocular albinism type 1 (OA1) G-protein-coupled receptor functions with MART-1 at early stages of melanogenesis to control melanosome identity and composition. Hum. Mol. Genet. 18, 4530–4545 (2009). doi:10.1093/hmg/ddp415; pmid:19717472.

67. C. Bissig, P. Croisé, X. Heiligenstein, I. Hurbain, G. M. Lenk, E. Kaufman, R. Sannerud, W. Annaert, M. H. Meisler, L. S. Weisman, G. Raposo, G. van Niel, The PIKfyve complex regulates the early melanosome homeostasis required for physiological amyloid formation. J. Cell Sci. 132 (2019). doi:10.1242/jcs.229500; pmid:30709920.

68. M. C. Liggins, J. L. Flesher, S. Jahid, P. Vasudeva, V. Eby, S. Takasuga, J. Sasaki, T. Sasaki, R. E. Boissy, A. K. Ganesan, PIKfyve regulates melanosome biogenesis. PLoS Genet. 14, e1007290 (2018). doi:10.1371/journal.pgen.1007290; pmid:29584722.

69. M. Seiji, T. B. Fitzpatrick, M. S. Birbeck, The melanosome: a distinctive subcellular particle of mammalian. J. Invest. Dermatol. 36, 243–252 (1961). doi:10.1038/jid.1961.42; pmid:13749804.

70. A. Lo Cicero, C. Delevoye, F. Gilles-Marsens, D. Loew, F. Dingli, C. Guéré, N. André, K. Vié, G. Van Niel, G. Raposo, Exosomes released by keratinocytes modulate melanocyte pigmentation. Nat. Commun. 6, 7506 (2015). doi:10.1038/ncomms8506; pmid:26103923.

71. Y. Yamaguchi, V. J. Hearing, Physiological factors that regulate skin pigmentation. BioFactors 35, 193–199 (2009). doi:10.1002/biof.29; pmid:19449448.

72. G. Cardinali, G. Bolasco, N. Aspite, G. Lucania, L. V. Lotti, M. R. Torrisi, M. Picardo, Melanosome transfer promoted by keratinocyte growth factor in light and dark skin-derived keratinocytes. Journal of Investigative Dermatology 128, 558–567 (2008). doi:10.1038/sj.jid.5701063; pmid:17882267.

73. P. D. Donatien, G. Hunt, C. Pieron, J. Lunec, A. Taieb, A. J. Thody, “The expression of functional MSH receptors on cultured human melanocytes” (Springer-Verlag, 1992).

74. M. E. Truong, S. Bilekova, S. P. Choksi, W. Li, L. J. Bugaj, K. Xu, J. F. Reiter, Vertebrate cells differentially interpret ciliary and extraciliary cAMP. Cell 184, 2911–2926.e18 (2021). doi:10.1016/j.cell.2021.04.002; pmid:33932338.

75. J. E. Siljee, Y. Wang, A. A. Bernard, B. A. Ersoy, S. Zhang, A. Marley, M. Von Zastrow, J. F. Reiter, C. Vaisse, Subcellular localization of MC4R with ADCY3 at neuronal primary cilia underlies a common pathway for genetic predisposition to obesity. Nat. Genet. 50, 180–185 (2018). doi:10.1038/s41588-017-0020-9; pmid:29311635.

76. R. N. Ozdeslik, L. E. Olinski, M. M. Trieu, D. D. Oprian, E. Oancea, Human nonvisual opsin 3 regulates pigmentation of epidermal melanocytes through functional interaction with melanocortin 1 receptor. Proc. Natl. Acad. Sci. U. S. A. 166, 11508–11517 (2019). doi:10.1073/pnas.1902825116; pmid:31097585.

77. L. Le, J. Sirés-Campos, G. Raposo, C. Delevoye, M. S. Marks, Melanosome Biogenesis in the Pigmentation of Mammalian Skin. Integr. Comp. Biol. 61, 1517–1545 (2021). doi:10.1093/icb/icab078; pmid:34021746.

78. M. V. Schiaffino, C. Tacchetti, The ocular albinism type 1 (OA1) protein and the evidence for an intracellular signal transduction system involved in melanosome biogenesis. [Preprint] (2005). 10.1111/j.1600-0749.2005.00240.x.

79. B. Bueschbell, P. Manga, A. C. Schiedel, The Many Faces of G Protein-Coupled Receptor 143, an Atypical Intracellular Receptor. Frontiers Media S.A. [Preprint] (2022). 10.3389/fmolb.2022.873777.

80. G. van Niel, S. Charrin, S. Simoes, M. Romao, L. Rochin, P. Saftig, M. S. Marks, E. Rubinstein, G. Raposo, The Tetraspanin CD63 Regulates ESCRT-Independent and - Dependent Endosomal Sorting during Melanogenesis. Dev. Cell 21, 708–721 (2011). doi:10.1016/j.devcel.2011.08.019; pmid:21962903.

81. C. Delevoye, M. S. Marks, G. Raposo, Lysosome-related organelles as functional adaptations of the endolysosomal system. Curr. Opin. Cell Biol. 59, 147–158 (2019). doi:10.1016/j.ceb.2019.05.003; pmid:31234051.

82. S. Mahanty, Skin lamellar bodies: a unique set of lysosome-related organelles. Front. Cell Dev. Biol. 13 (2025). doi:10.3389/fcell.2025.1597696.

83. S. L. Bowman, J. Bi-Karchin, L. Le, M. S. Marks, The road to lysosome-related organelles: Insights from Hermansky-Pudlak syndrome and other rare diseases. Traffic 20, 404–435 (2019). doi:10.1111/tra.12646; pmid:30945407.

84. K. Newell-Litwa, G. Salazar, Y. Smith, V. Faundez, S. L. Schmid, Roles of BLOC-1 and Adaptor Protein-3 Complexes in Cargo Sorting to Synaptic Vesicles. Mol. Biol. Cell 20, 1441–1453 (2009). doi:10.1091/mbc.E08.

85. S. Niwa, L. Tao, S. Y. Lu, G. M. Liew, W. Feng, M. V. Nachury, K. Shen, BORC Regulates the Axonal Transport of Synaptic Vesicle Precursors by Activating ARL-8. Current Biology 27, 2569–2578.e4 (2017). doi:10.1016/j.cub.2017.07.013; pmid:28823680.

86. A. Evstratova, S. Chamberland, V. Faundez, K. Tóth, Vesicles derived via AP-3-dependent recycling contribute to asynchronous release and influence information transfer. Nat. Commun. 5 (2014). doi:10.1038/ncomms6530; pmid:25410111.

87. R. M. Leonhardt, N. Vigneron, J. S. Hee, M. Graham, P. Cresswell, Critical residues in the PMEL/Pmel17 N-terminus direct the hierarchical assembly of melanosomal fibrils. Mol. Biol. Cell 24, 964–981 (2013). doi:10.1091/mbc.E12-10-0742; pmid:23389629.

88. J. F. Berson, A. C. Theos, D. C. Harper, D. Tenza, G. Raposo, M. S. Marks, Proprotein convertase cleavage liberates a fibrillogenic fragment of a resident glycoprotein to initiate melanosome biogenesis. Journal of Cell Biology 161, 521–533 (2003). doi:10.1083/jcb.200302072; pmid:12732614.

