## Supplementary Methods and Figures for "Ciliary regulation of melanogenesis unveils moonlighting roles for Joubert Syndrome proteins"

**This file includes:**

Materials and Methods
Figs. S1 to S4
References

### Materials and Methods

#### Antibodies

Monoclonal antibodies used in this study include: anti-β-tubulin (clone D3U1W, 1:1,000 for immunoblotting; Cell Signaling, cat# 86298S), rabbit anti-GAPDH (clone 14C10, 1:1,000 for immunoblotting; Cell Signaling, cat# 2118S), mouse anti-γ-tubulin (clone GTU-88, 1:500 for IFM; Sigma-Aldrich, cat# T5326), mouse anti-TYRP1 (clone TA99, 1:50 for IFM; antibody produced in lab using a hybridoma cell line purchased from American Type Culture Collection, cat# HB-8704), rabbit anti-TYRP1 (clone H-90, 1:500 for immunoblotting; Santa Cruz, cat# sc-25543), rabbit anti-TYRP1 (clone αPep1, 1:500 for immunoblotting; gift from Vincent J. Hearing), rabbit anti-PMEL (clone αPep13h (*1*), 1:200 for immunoblotting), mouse anti-PMEL (clone HMB45, 1:200 for IFM; Enzo Life Sciences, cat# C34930), rabbit anti-core amyloid fragment (clone I-51, 1:200 for immunoblotting; kind gift from Ralf M. Leonhardt), rabbit anti-TYR (clone αPep7h-msm (*2*), 1:50 for immunoblotting, 1:500 for IFM); mouse anti-CREB (clone 86B10, 1:1,000 for immunoblotting; Cell Signaling, cat# 9104S), rat anti-LAMP2A (clone GL2A7, 1:250 for IFM; Developmental Studies Hybridoma Bank, cat# GL2A7-s), mouse anti-LAMP2A (clone H4B4, 1:100 for IFM; Developmental Studies Hybridoma Bank, cat# H4B4-c). Polyclonal antibodies used in this study include: rabbit anti-TMEM138 (1:50 for immunoblotting; Sigma-Aldrich, cat# HPA042373), rabbit anti-Arl13b (1:500 for IFM; Proteintech, cat# 17711-1-1-AP), rabbit anti-pCREB$^{Ser133}$ (1:1,000 for immunoblotting; Sigma-Aldrich, cat# 06-519), rabbit anti-ERK1/2 (1:1,000 for immunoblotting; Thermo Fisher, cat# 44-654G), rabbit anti-pERK1/2$^{Thr202/Tyr204}$ (1:1,000 for immunoblotting; Cell Signaling, cat# 9101S), and rabbit anti-LAMP2A (1:250 for IFM; Abcam, cat# ab18528). Extensively cross-adsorbed species-specific secondary antibodies from donkey conjugated either to Alexa Fluor 488, Alexa Fluor 594, or Alexa Fluor 647 were used for IFM at 1:500, or to IRDye-790CW, or IRDye-680LT were used for immunoblotting at 1:5,000 (Jackson ImmunoResearch Laboratories, Inc.).

#### Cloning and plasmid generation

PCR-amplification of DNA fragments from existing vector backbones was carried out on a C1000 Touch thermal cycler (Biorad) using 10-50 ng plasmid DNA, 1X Phusion high-fidelity DNA polymerase master mix (New England Biolabs), and 0.5 µM forward and reverse primers (Table S1) with matching annealing temperature (1-5 °C below highest melting temperature); when primers' GC content exceeded 60%, 5-10% (v/v) molecular biology-grade DMSO (New England Biolabs) was added. Restriction enzymes and buffers were all purchased from New England Biolabs, and used according to the manufacturer's instructions. DNA fragments were isolated using the Zymoclean gel DNA recovery kit (Zymo Research) after migration on 0.8 % ultra-pure agarose (Thermo Fisher) gels. For each Gibson assembly reaction, 50-100 ng plasmid backbone was used, and various backbone-to-fragment molar ratios (1:3, 1:5, or 1:8) were tested. Gibson assembly master mix was composed of 1.2 µl T5 exonuclease (New England Biolabs), 20 µl Phusion high-fidelity DNA polymerase (New England Biolabs), 160 µl Taq DNA ligase (New England Biolabs), 700 µl nuclease-free water (Thermo Fisher), and 320 µl of isothermal reaction buffer containing 0.5 M Tris-HCl, pH 7.4 (Thermo Fisher), 5 mM NAD (Applichem), 0.05 M Magnesium Chloride (Sigma-Aldrich), 0.05 M DTT (Fermentas), 1mM dNTP mix (Fermentas), and 25% (v/w) PEG-8000 (Sigma-Aldrich). Standard ligation was carried out using the T4 DNA ligase master mix (New England Biolabs) according to the manufacturer's instructions. Ligation or Gibson assembly products were transformed into 10-beta (New England Biolabs) or DH5α (Thermo Fisher) competent cells, and Ampicilin- or Kanamycin-resistant clones were subcultured for DNA isolation using the PureLink HiPure plasmid filter Mini-, Midi-, or MaxiPrep kits (Thermo Fisher). Sequences were validated by restriction digest, and whole plasmid sequencing.

Retroviral plasmids for recombinant protein expression

Plasmid pBMN-HA-hTMEM138-IRES-Hygro was created by Gibson Assembly using the hTMEM138 cassette amplified by PCR from pEGFP-C1-hTMEM138-EGFP (*3*) with primers 1 (containing an overhang with a start codon and an HA-epitope tag upstream of hTMEM138) and 2 in the NotI restriction site of pBMN-IRES-Hygro (*4*) (see Table S1 for primers). This plasmid was further edited using the Q5 site-directed mutagenesis kit (New England Biolabs) with primers 3 and 4 to increase hTMEM138 resistance to sh*mTmem138* #1 without altering its amino acid composition (Table S2). Plasmid pBMN-IRES-Puro was generated by Gibson Assembly using the Puromycin resistance cassette amplified by PCR from plasmid pLKO.1-Puro (High-throughput Screening Core of the University of Pennsylvania) with primers 5 and 6 into the BstXI restriction sites flanking the Hygromycin B resistance cassette of pBMN-IRES-Hygro. Plasmid pBMN-ARL13B-mCherry-IRES-Puro was generated by Gibson Assembly using the ARL13B-mCherry fragment amplified by PCR from pcDNA-ENTR-ARL13B-mCherry (a kind gift from Duarte Barral, NOVA University, Portugal) with primers 7 and 8 into the NotI/ XhoI restriction sites of pBMN-IRES-Puro. Plasmid pMXs-IRES-Blast was purchased from Cell Biolabs. Plasmid pMXs-EPAC-$S^{H187}$-IRES-Blast was generated by Gibson assembly using EPAC-$S^{H187}$ cassette amplified by PCR from pcDNA3-EPAC-$S^{H187}$ (provided by Robert J. Lee, University of Pennsylvania, PA, USA; Addgene#170348) with primers 9 and 10 into the NotI/ XhoI restriction sites of pMXs-IRES-Blast to monitor the cytosolic activity of EPAC-$S^{H187}$. To generate plasma membrane-localized EPAC (pmEPAC), pMXs-EPAC-$S^{H187}$-IRES-Blast was edited to insert the plasma membrane targeting sequence ('GCIKSKRKDKD') derived from the Lyn kinase (*5*) at the N terminus of EPAC using the Q5 site-directed mutagenesis kit (New England Biolabs) with primers 11 and 12. To generate ciliary membrane-localized EPAC (cilioEPAC), ARL13B was fused to the N-terminus of EPAC with a 19-amino acid linker by Gibson assembly using the ARL13B cassette amplified by PCR from pBMN-ARL13B-mCherry-IRES-Puro with primers 13 and 14 into the PacI/ SbfI restriction sites of pMXs-EPAC-$S^{H187}$-IRES-Blast.

Lentiviral plasmids for recombinant protein expression

Plasmid pLenti-hPGK-IRES-Hygro was created from the pGW-EGFP-WPRE vector (a kind gift of Stefano Rivella, CHOP Research Institute, Philadelphia, PA) after removal of the EGFP-WPRE cassette using the Q5 site-directed mutagenesis kit (New England Biolabs) with primers 17 and 18, and insertion of the IRES-Hygromycin B fragment amplified by PCR from plasmid pBMN-IRES-Hygro using primers 19 and 20 within NotI/ PacI restriction sites. Plasmid pLenti-hPGK-EGFP-$hMITF^{S69+73A}$-IRES-Hygro was generated by Gibson assembly using the EGFP-$hMITF^{S69+73A}$ fragment amplified by PCR from pEGFP-C1-EGFP-$hMITF^{S69+73A}$ (*6*) using primers 21 and 22 into the AgeI restriction site of pLenti-hPGK-IRES-Hygro. Lentiviral packaging plasmids psPAX2 and pMD2.G encoding the vesicular stomatitis virus glycoprotein were gifts from Didier Trono (University of Geneva, Switzerland; Addgene# 12260 and 12259, respectively).

Short hairpin RNA (shRNA) constructs for gene knockdown

All shRNAs were generated in and expressed from the lentiviral vector pLKO.1 (Addgene# 10878), which confers puromycin resistance. Most constructs were from the TRC library (*7*), and obtained from the High-throughput Screening Core of the University of Pennsylvania. Sequences are documented in Table S3. Non-target shRNA construct ('GCGCGATAGCGCTAATAATTT', Sigma-Aldrich) and pLKO.1-Puromycin empty backbone were used as controls. Additional lentiviral vectors encoding shRNAs to mouse *Tmem138*, mouse *Ccdc114*, and mouse *Ccdc149* were made in the lab using a described protocol (*8*). Briefly, shRNA sequences were either obtained from the Broad Institute TRC shRNA library (https://www.broadinstitute.org/rnai-consortium/rnai-consortium-shrna-library), or custom designed according to the Broad Institute TRC shRNA design process guidelines (https://portals. broadinstitute.org/gpp/public/resources/ rules). 5'-end phosphorylated sense and anti-sense oligonucleotides containing the shRNA original and reverse complement sequences, loop domain, transcriptional terminator sequences, and AgeI and EcoRI restriction sites overhangs were generated for each construct, and are listed in Table S4. Sense and anti-sense oligonucleotides were annealed at a 1:1 ratio in pH 7.4 aqueous buffer containing 10 mM Potassium Acetate (Fisher Scientific), 200 µM Magnesium Acetate (Sigma-Aldrich), and 30 mM HEPES (Fisher Scientific), by successively incubating at 95 °C for 4 min, 70 °C for 10 min, and progressively cooling down at 1 °C/min to 10 °C. The annealed oligonucleotides were subcloned into the AgeI and EcoRI restriction sites of the pLKO.1-Puromycin empty backbone by overnight ligation at 4 °C with T4 ligase (New England Biolabs).

#### Cell culture

Mouse melanocyte cell lines

Immortalized, non-cancerous mouse melan-*Ink4a* melanocytes derived from the C57BL/6J-*Ink4a*$^{-/-}$ mice (*9*) were used as wild-type cells, or transduced with retro- or lentiviruses to either transiently or stably express recombinant proteins or shRNAs. Cell lines were maintained at 37 °C and 10 % $CO_2$ in RPMI 1640 medium supplemented with 10 % fetal bovine serum (FBS; Atlanta Biologicals), 2 mM L-glutamine (Corning), and 200 nM 12-O-tetradecanoylphorbol-13-acetate (TPA; Sigma-Aldrich), and split 1:4-1:5 twice a week up to 30 passages. All cell lines were authenticated by both changes in pigmentation using bright field microscopy, and expression of melanocyte-specific cargoes (e.g. TYRP1, PMEL, etc.) using immunofluorescence microscopy (IFM). Cells were also routinely tested for mycoplasma contamination using the MycoAlert Mycoplasma Detection Kit (Lonza).

Primary human skin cells

Primary human skin cells were isolated from deidentified foreskin samples provided by the Skin Biology and Disease Resource-based Center (SBDRC) at the University of Pennsylvania after circumcision surgeries of neonates with dark skin phototypes (4-6 on Fitzpatrick's scale). Briefly, foreskins were rinsed in phosphate-buffered saline (PBS; Corning) supplemented with 100 U/ml penicillin (Sigma-Aldrich) and 25 µg/ml gentamycin (Gemini Bio), minced into small pieces (2 mm thick x 5-10 mm long), and incubated for 16 h at 4 °C in digestion solution [500 µg/ml thermolysin (Thermo Fisher), 5mM $CaCl_2$ (Sigma-Aldrich), 33.5 mM KCl (Sigma-Aldrich), 710 mM NaCl (VWR Scientific) and 50 mM HEPES (Thermo Fisher)]. After digestion, dermis and epidermis were separated by mechanical peel-off, and further digested under constant agitation at 37 °C in 0.125 U/ml collagenase H (Sigma-Aldrich) for 3 h (for dermis), or 0.025% trypsin-EDTA (Thermo Fisher) for 25 min (for epidermis). Dermal and epidermal cell lysates were passed through a 100-µm cell strainer, and cells were collected after centrifugation at 300 g for 10 min.

For primary human fibroblast isolation and expansion, dermal cells were seeded at 60,000 cells/cm$^2$ at P0, then 8,000 cells/cm$^2$ at following passages, and maintained at 37 °C and 5 % $CO_2$

in DMEM medium (Thermo Fisher) supplemented with 10 % FBS (Thermo Fisher), 100 U/ml penicillin (Sigma-Aldrich) and 25 μg/ml gentamycin (Gemini Bio) for up to 15 passages. Cells were authenticated by both morphology using bright field microscopy, and by their ability to produce extracellular matrix upon addition of 50 μg/ml ascorbic acid (Sigma-Aldrich) to the culture media.

For primary human melanocyte isolation and expansion, epidermal cells were seeded at 300,000 cells/cm$^2$ at P0, and cultured at 37 °C and 10 % $CO_2$ in RPMI 1640 medium supplemented with 10 % FBS (Atlanta Biologicals), 2 mM L-glutamine (Corning), 200 nM TPA (Sigma-Aldrich), 100 μg/ml geneticine (Sigma-Aldrich), 0.2 nM FGF-2 (R&D), 100 U/ml penicillin and 25 μg/ml gentamycin. After P0, melanocytes were seeded at 10,000 cells/cm$^2$, and maintained in Medium 254 with HMGS supplements (Thermo Fisher), 100 U/ml penicillin (Sigma-Aldrich), and 25 μg/ml gentamycin (Gemini Bio) for up to 8 passages. Cells were authenticated by pigment content using bright field microscopy, and expression of TYRP1 by IFM.

For primary keratinocyte isolation, epidermal cells were seeded at 40,000 cells/cm$^2$ at P0, and 6,500 cells/cm$^2$ at following passages, onto sub-confluent layers of mitotically-inactive primary human fibroblasts irradiated at 60 Gy using an X-RAD 320ix irradiator (Precision Xray). Keratinocytes were maintained at 37 °C and 10 % $CO_2$ in DMEM-Ham F12 (3:1) medium (Thermo Fisher) supplemented with 5 % fetal clone II serum (Hyclone), 36.5 mM sodium bicarbonate (Sigma-Aldrich), 24 μg/ml adenine (Sigma-Aldrich), 10 ng/ml EGF (Thermo Fisher), 5 μg/ml insuline (Sigma-Aldrich), 0.4 μg/ml hydrocortisone (Sigma-Aldrich), 0.212 μg/ml isoprenaline (Sigma-Aldrich), 100 U/ml penicillin, and 25 μg/ml gentamycin for up to 10 passages. Keratinocytes were authenticated by both morphology using bright field microscopy, and by their ability to form pluristratified epithelia when cultured at the air-liquid interface.

**Virus production and transductions**

For retrovirus production, Plat-E cells (*10*) were seeded at 95,000 cells/cm$^2$ in a 10-cm Cell-Bind Petri dish (Corning), and maintained at 37 °C and 5 % $CO_2$ in DMEM high glucose medium (Thermo Fisher) supplemented with 10 % FBS (Thermo Fisher), 2 mM L-glutamine (Corning), and 1mM sodium pyruvate (Thermo Fisher). After 24h, the cells were transfected by adding drop-by-drop 3 ml of Opti-MEM medium (Thermo Fisher) containing 4 % (v/v) Lipofectamine 2000 (Thermo Fisher) and 24 μg of retroviral DNA. For lentivirus production (*11*), HEK293T cells were seeded at 95,000 cells/cm$^2$ in a 10-cm Cell-Bind Petri dish (Corning), and maintained at 37 °C and 5 % $CO_2$ in DMEM high glucose medium (Thermo Fisher) supplemented with 10 % FBS (Thermo Fisher), and 2 mM L-glutamine (Corning). After 24h, the cells were transfected by adding drop-by-drop 3 ml of Opti-MEM medium (Thermo Fisher) containing 4 % (v/v) Lipofectamine 2000 (Thermo Fisher), 21 μg of lentiviral DNA, 15.75 μg of psPAX2, and 5.25 μg of pMD2.G (see section on Lentiviral plasmids).

One to two days after Plat-E or HEK293T transfection, medium was replaced with 10 ml of fresh media, and the cells were maintained in culture for 1 to 3 additional days. Virus-containing supernatants were centrifuged at 300 g for 5 min and passed through a 45-μm membrane filter. Viral particles were used freshly or aliquoted and frozen at -80°C for later use.

Melan-*Ink4a* cells and primary human melanocytes were infected with viral particles in a 1:1 ratio with fresh medium. One to two days after infection, supernatants were discarded, and fresh medium was added to the transduced cells. In most experiments, stable transductants were selected for 2 weeks using 500 µg/ml hygromycin B, 2 µg/ml puromycin, or 5 µg/ml blasticidin depending on antibiotic resistance conferred by plasmids expression.

**Reconstructed organotypic skin**

Organotypic skin was produced from primary human keratinocytes and fibroblasts, and melan-*Ink4a* melanocytes using the 'self-assembly' method as reported previously (*12–15*). Melan-*Ink4a* cells were chosen over primary human melanocytes as they divide faster, produce only dark/brown eumelanins, and are genetically homogeneous. Briefly, primary human fibroblasts were seeded at 8,000 cells/$cm^2$ in 6-well plates containing pre-cut paper rings, and grown in fibroblast medium supplemented with 50 µg/ml ascorbic acid for 4-5 weeks to form cohesive fibroblast sheets. Three fibroblast sheets were then stacked on top of each other to build a thicker dermal tissue, onto which primary keratinocytes and melanocytes were seeded at 100,000 cells/$cm^2$ and 50,000 cells/$cm^2$ respectively. These organotypic tissues were grown for 7 days immersed in keratinocyte medium containing 50 µg/ml ascorbic acid, then raised to the air-liquid interface using custom-made pedestals (designed and 3D-printed by Michael Hast at University of Pennsylvania, PA, USA) for 14 more days in keratinocyte medium containing 50 µg/ml ascorbic acid but no EGF to promote epidermal stratification.

**shRNA screen**

Melan-*Ink4a* melanocytes were transiently depigmented using 300 µM N-Phenylthiourea (PTU; Sigma-Aldrich) for 2 weeks, then seeded into 24-well plates in duplicates. Cells were transduced with recombinant lentiviruses to express sh*NT* or each of 3-5 shRNAs to various ciliopathy genes, the BLOC-1 subunit gene *Hps9* (positive control), or *B2m* (as a pigment-unrelated negative control; Tables S3-4). Transductants were selected with 2 µg/ml puromycin, and after one day PTU was removed to enable pigment recovery. After 3-7 days, melanocytes were stained with 1 µg/ml DAPI (Sigma-Aldrich) for 20 min at 4 °C, washed with PBS, trypsinized, transferred to 96-well plates, fixed in 3.7 % formaldehyde (VWR Scientific) for 10 min, and washed three times in PBS. The cells were then resuspended in PBS + 1% FBS, and analyzed by flow cytometry. Side scatter was measured in live (DAPI-negative) cells, and 20,000 events were recorded for each sample. Pigment recovery was expressed on a scale from 0 (no recovery) to 1 (full recovery) after averaging side scatter across duplicates, and normalizing sample mean values. Normalization was achieved after subtracting the mean side scatter value of unpigmented sh*NT* cells remaining in PTU, and dividing by the mean side scatter value of pigment-recovered sh*NT* cells off of PTU.

**Proliferation assay**

Melan-*Ink4a* cells stably transduced to express sh*NT*, sh*mTmem138* #1, sh*mTmem138* #2, or sh*mTmem138* #1 and EGFP-$hMITF^{S69+73A}$ were seeded at 50,000 cells per well in 6-well plates in duplicates. Cells were treated daily for 6 days with 100 pM [$Nle^4$, D-$Phe^7$]-α-MSH (Tocris) or vehicle diluted in normal media, trypsinized, and live cells were counted after Trypan blue (Corning) exclusion. Live cell counts were averaged across replicates, and expressed as fold change between vehicle and MSH treatment.

**Sample collection for pigment, protein, or mRNA content assays**

<u>Cultured cells</u>

Melanocytes were grown in 10-cm Petri dishes until they reached 85-100 % confluency. In some experiments, cells were either treated for 3-5 h with vehicle (DMSO; Sigma-Aldrich), or 25 nM bafilomycin A1 (BafA1; Sigma-Aldrich), or stimulated for various periods of time with vehicle, 3-100 pM [Nle$^4$, D-Phe$^7$]-α-MSH (Tocris), or 0.6 µM Forskolin (FSK; Cayman Chemical Company). Prior to harvest, cell shape and pigmentation were assessed qualitatively by phase contrast and bright field microscopy, respectively, using a PrimoVert inverted microscope (Zeiss). Cells were then rinsed with ice-cold PBS, and incubated on ice for 5 min in ice-cold PBS containing 5 mM EDTA (Sigma-Aldrich). Cells were then gently harvested using a cell lifter, and collected by centrifugation at 500 g for 5 min. Cell pellets were resuspended in PBS supplemented with 1 % FBS, and centrifuged again to eliminate EDTA. Pellets were then lysed directly, or frozen at -80°C for later use.

<u>Organotypic skin</u>

Organotypic skin samples were rinsed in PBS, photographed, minced into small pieces (0.5 cm x 0.1 cm), and fixed in 3.7 % formaldehyde (VWR Scientific). Samples were dehydrated and embedded in paraffin for histological processing by the SBDRC at the University of Pennsylvania.

**Melanin content assays**

Melanin content in cell lysates was quantified as previously described (*16*). Specifically, cell pellets were resuspended in ice-cold 50 mM Tris-HCl, pH 7.4 (Thermo Fisher), 2 mM EDTA (Sigma-Aldrich), 150 mM NaCl (VWR Scientific), 1 mM DTT (Sigma-Aldrich), 1X protease inhibitor cocktail (Roche), and sonicated on ice 3 x 3 s using the Sonic Dismembrator Model 100 (Thermo Fisher). Sonicates were centrifuged at 20,000 g for 15 min at 4 °C, and supernatants were isolated for protein concentration assay using a BCA kit (Thermo Fisher). Melanin-containing pellets were then rinsed in equal parts of pure ethanol (VWR Scientific) and diethyl ether (VWR Scientific), and incubated in 2 M NaOH (Thermo Fisher) containing 20 % (v/v) DMSO (Sigma-Aldrich) at 60 °C until complete melanin dissolution. Absorbance was measured by spectrophotometry at 492 nm, and divided by protein concentration in each sample to normalize for protein content. Samples were analyzed in duplicates for each experimental condition in each experiment. Results were normalized by dividing the mean value of every condition by the mean value of the untransduced, vehicle-treated, or sh*NT*-expressing cells.

**Cell lysate preparation and immunoblotting**

<u>Soluble protein lysates</u>

Cell pellets were resuspended in ice-cold 62.5 mM Tris-HCl, pH 7.4 (Thermo Fisher), 2 % (v/v) SDS (Thermo Fisher), 10 % (v/v) Glycerol (Amresco), 1X protease inhibitor cocktail (Roche), 5 mM Sodium Fluoride (Sigma-Aldrich), and 1 mM Sodium Orthovanadate (Sigma-Aldrich), and sonicated on ice using the Sonic Dismembrator (Thermo Fisher, Model 100). Sonicates were clarified by centrifugation at 20,000 g for 15 min at 4°C, and supernatants were collected for protein concentration assay using a BCA kit (Thermo Fisher), and frozen at -80°C for later use.

<u>Insoluble protein lysates</u>

Cell pellets were resuspended in ice-cold 10 mM Tris-HCl, pH 7.4 (Thermo Fisher), 140 mM Sodium Chloride (VWR Scientific), 1 mM EDTA (Sigma-Aldrich), 1 % (v/v) Triton-X-100

(Thermo Fisher), 1 % (v/v) Sodium Deoxycholate (Sigma-Aldrich), 1 % (v/v) SDS (Thermo Fisher), and 1X protease inhibitor cocktail (Roche), and sonicated on ice using the Sonic Dismembrator (Thermo Fisher, Model 100). Sonicates were clarified by centrifugation at 20,000 g for 15 min at 4°C, and supernatants were collected for protein concentration assay using a BCA kit (Thermo Fisher). Insoluble protein pellets were resuspended in 50 mM Tris-HCl, pH 7.4 (Thermo Fisher), 8 M Urea (Sigma-Aldrich), and 5 mM DTT (Sigma-Aldrich), shaken at 4 °C for 2h, and clarified by centrifugation at 20,000 g for 15 min at 4°C. Supernatants were collected and frozen at -80°C for later use.

Immunoblotting

Samples were thawed on ice, diluted in concentrated Laemmli buffer with 2 % (v/v) 2-mercaptoethanol, and boiled at 95 °C for 10 min. Samples were then run on 10 or 15 % polyacrylamide gels, and transferred to reinforced nitrocellulose membranes (Sigma-Aldrich). Membranes were stained with Ponceau Red (Thermo Fisher) to validate protein transfer, blocked for 1h at room temperature in blocking buffer [TBS/0.2 % Tween-20 (Sigma-Aldrich; TBS-T) supplemented with 5 % (w/v) Milk or 5 % (w/v) BSA (Sigma-Aldrich; for phosphorylated proteins)], and incubated overnight at 4°C with primary antibodies diluted in blocking buffer. Membranes were rinsed 5 x 10 min in TBS-T, and incubated with secondary antibodies diluted in blocking buffer for 1 h at room temperature. Membranes were rinsed 5 x 10 min in TBS-T, and then imaged using a LICOR Odyssey Clx imaging system. Image analysis was carried out using ImageJ [v1.53j, (*17*)]. Briefly, lanes were delineated with the rectangle tool using the 'Select Lane' function, and intensity profile was generated using the 'Plot Lane' function. Peaks of interest and background were outlined from the intensity profile using the straight-line tool, and area under the curve was measured for each selected peak and background using the wand tool. Background values were subtracted from peak values in each lane, and data were expressed as the ratio of background-subtracted peak area values for proteins of interest over background-subtracted peak area values for loading control (GAPDH or β-Tubulin). Results were then normalized by dividing the mean value for every condition by the mean value of the control condition (untransduced, vehicle-treated, or sh*NT*-expressing cells).

**Real-time quantitative Polymerase Chain Reaction (RT-qPCR)**

RT-qPCR was carried out as previously described (*18*). Briefly, cell pellets were thawed on ice, lysed, sonicated, and processed for RNA extraction using the PureLink RNA mini kit (Thermo Fisher) as per the manufacturer's instructions. RNA concentration and purity were assessed using a Nanodrop (Thermo Fisher). For each sample, 1µg RNA was treated with 1 U/µl DNAse I (Thermo Fisher) at room temperature for 15 min. DNAse was inactivated by addition of 2 mM EDTA and incubation at 65 °C for 10 min. Samples were then reverse transcribed into cDNA using the high capacity RNA-to-cDNA kit (Applied Biosystems). RT-qPCR was carried out in a 384-well plate on a QuantStudio 7 Flex Real-Time PCR system (Thermo Fisher) by adding 50 to 100 ng cDNA, 10 µl fast SYBR green master mix (Applied biosystems), 375 nM forward and reverse primers (see Table S5), and RNAse-free water to a final volume of 20 µl in each well. Products were amplified for 45 cycles as follows: denaturation at 95 °C for 20 sec; annealing at 56 °C for 20 sec, and extension at 72 °C for 20 sec. Cycle threshold (Ct) values were averaged across three replicates, and differences between mean Ct values of each gene of interest (GOI) and housekeeping genes (HKG; *B2m*, *Tubb4a*, *Gapdh*) were computed as $\Delta Ct = Ct^{GOI} - Ct^{HKG}$, and relative quantities were determined using $2^{-\Delta Ct}$ values. Results were normalized by dividing the $2^{-\Delta Ct}$ values of every condition by the $2^{-\Delta Ct}$ value of the untransduced or sh*NT*-expressing cells.

**Electron Transmission Microscopy**

Subconfluent melan-*Ink4a* cells were rinsed in PBS, and pre-fixed for 2h at room temperature in 5 % paraformaldehyde (Polysciences), 72 mM Sodium Cacodylate, pH 7.4 (Sigma-Aldrich), 4 mM Sodium Chloride (Sigma-Aldrich), and 0.5 % glutaraldehyde (Polysciences). Cells were then fixed overnight at 4 °C in 5 % paraformaldehyde (Polysciences), 72 mM Sodium Cacodylate, pH 7.4 (Sigma-Aldrich), 4 mM Sodium Chloride (Sigma-Aldrich), and 2 % glutaraldehyde (Polysciences). Cells were harvested on ice using a cell lifter, centrifuged at 500 g for 5 min at 4°C, post-fixed in 2 % osmium tetroxide and 1.5 % potassium ferrycyanide for 1 hour at room temperature, and rinsed in distilled water prior to en bloc staining with 2 % uranyl acetate. After dehydration through a graded ethanol series, cells were infiltrated and embedded in EMbed-812 (Electron Microscopy Sciences). Thin sections were stained with uranyl acetate and SATO lead and examined with a Talos L120C electron microscope fitted with a Ceta 16M digital camera at the Electron Microscopy Resource Laboratory of University of Pennsylvania.

**Immunofluorescence, bright field, and phase contrast microscopy on fixed cells**

Melanocytes were seeded on phenol-free Matrigel (Corning)-coated coverslips, and grown in serum-containing or serum-deprived medium for 48-72 h. In some experiments, cells were treated for 3-5 h with vehicle (DMSO; Sigma-Aldrich), 25 nM bafilomycin A1 (BafA1; Sigma-Aldrich), or 25 µg/ml cycloheximide (CHX; Sigma-Aldrich) prior to fixation and staining as previously described (*19*). Specifically, coverslips were then gently washed in PBS, and then fixed with 3.7 % formaldehyde (VWR Scientific) in PBS at room temperature for 15 min. After three washes in PBS, coverslips were incubated for 1 h at room temperature with primary antibodies diluted in PBS/ 0.01 % BSA (Sigma-Aldrich)/ 0.02 % saponin (Sigma-Aldrich). Coverslips were washed three times in PBS, then incubated for 45 min at room temperature with secondary antibodies diluted in PBS/ 0.01 % BSA (Sigma-Aldrich)/0.02 % saponin (Sigma-Aldrich). Coverslips were washed three times in PBS and mounted onto slides using VectaShield with DAPI (Vector Laboratories). Three-color images (DAPI, AlexaFluor-488, AlexaFluor-594) were collected on a DMI6 inverted widefield fluorescence microscope (Leica Biosystems) equipped with an EL6000 mercury metal halide bulb lamp (Leica Biosystems), an Orca Flash4.0v2 camera (Hamamatsu), and Plan Apo 100X oil immersion objective lenses (NA 1.47) using the LAS X software (Leica Biosystems). Four-color images (DAPI, AlexaFluor-488, AlexaFluor-546, AlexaFluor-647) were acquired on an AxioObserver 7 inverted widefield fluorescence microscope (Zeiss) equipped with a multichannel LED light source, an Axiocam 807 mono sCMOS camera (Zeiss) and Plan Apo 63X oil immersion objective lenses (NA 1.4) using the Zen Blue software (Zeiss). Cell morphology and pigmentation were assessed by phase contrast and bright field microscopy, respectively, using the DMI6 inverted microscope, or the AxioVert A1 microscope equipped with an Axiocam 305 color camera (Zeiss) using the Zen Lit software (Zeiss). Images were further processed and analyzed using ImageJ [v1.53j, (*17*)].

**cAMP live cell imaging**

Melan-*Ink4a* melanocytes were retrovirally transduced to express the mTurquoise2 (CFP variant) and Venus (YFP variant) biosensor EPAC-$S^{H187}$ (EPAC), plasma membrane targeted EPAC-$S^{187}$ (pmEPAC), or cilia-targeted EPAC-$S^{187}$ (cilioEPAC), and selected for 3 days with 5 μg/ml blasticidin. Transductants were then split, and 20,000 cells were seeded into each well of phenol-free Matrigel (Corning)-coated live-cell imaging 8-well chambers (Cellvis). Cells were then retrovirally transduced again to express mCherry-Arl13b, and cultured for 48-72 h in serum-free media. Transduced cells were imaged in a pH 7.4 imaging buffer containing Hank's Balanced Salt Solution (HBSS) supplemented with 20 mM HEPES (Thermo Fisher), 5.6 mM D-Glucose (Sigma-Aldrich), and 1.8 mM $CaCl_2$ (Sigma-Aldrich). Imaging was performed using an IX83 microscope (Olympus, Tokyo Japan) equipped with a 60X Plan Apo 1.4 NA oil immersion objective, a spinning disk attachment (Olympus DSU), and an Orca Flash 4.0 sCMOS camera (Hamamatsu, Tokyo Japan), using MetaFluor software (Molecular Devices, Sunnyvale, CA). Illumination was provided by a Xcite 120 LED Boost system (Excelitas Technologies, Pittsburgh, PA) at 30% power using standard CFP and YFP filters (Chroma Technologies, Bellows Falls, VT). CFP excitation used a 436/20 band-pass excitation filter with 455 long-pass dichroic. YFP excitation used a 500/20 band-pass excitation filter with 515 long-pass dichroic. Emission filters, controlled separately (Sutter Lamda 10-2), allowed both CFP (470/24 band-pass filter) and YFP (535/30 for YFP filter) to be collected with CFP excitation. Visualization of mCherry was done at the same time using a standard TRITC filter set (554/23 nm excitation, 609/54 emission, 573 long-pass dichroic, LED-TRITC-A-OMF from Semrock, Rochester, NY). CFP, FRET and mCherry images were collected every 10 seconds for 18 min in selected mCherry-Arl13b positive cells at baseline for 3 minutes, and then after stimulation with 500 pM [$Nle^4$, D-$Phe^7$]-α-MSH (Tocris) for 5 minutes, and with 20 μM Forskolin (FSK; Cayman Chemical Company) and 200 μM 3-Isobutyl-1-methylxanthine (IBMX; Sigma-Aldrich) for the final 5 minutes. FRET signals are expressed as CFP/YFP emission ratios after CFP excitation.

### Quantitative image analyses

Colocalization quantification

Colocalization between pairs of fluorescently-labeled proteins was quantified in fixed cells as the percentage of area of overlap using a custom-made semi-automatic macro on ImageJ [v1.53j, (*17*)] adapted from an image analysis pipeline previously described (*20*). Briefly, IFM pictures were first deconvolved with the Microvolution deconvolution plug-in on ImageJ [v1.53j, (*17*)] using the least number of iterations to avoid creating artifacts. Single z-plane images were then generated from deconvolved IFM (dIFM) z-stack series, and contrast and brightness values associated with each fluorescent channel in a given experiment were adjusted identically for all images. Local background was subtracted using the "rolling ball" algorithm with "sliding paraboloid" using a 10-pixel rolling ball radius. Contour masks were drawn to delineate both perinuclear and cytoplasmic surface areas of selected cells. After excluding the perinuclear region from each selected cell, images were converted to binary images using the Bernson algorithm of the "Auto Local Threshold" function with a 15-pixel radius. Structures of <5 pixels in area were considered background, and filtered out using the "Analyze Particles" function. To generate an overlap image, two binary images were multiplied using the "Image Calculator" function, and the areas of the individual and overlap binary images were computed with the "Measure" function. Percentage of overlap was calculated as the percentage of area overlap relative to total area of the individual labels.

Cytoplasmic-to-perinuclear signal intensity ratio quantification

Cytoplasmic-to-perinuclear signal intensity ratio was quantified in fixed cells using a custom-made semi-automatic macro on ImageJ [v1.53j, (*17*)]. Briefly, IFM pictures were deconvolved, converted into single z-planes, adjusted for brightness, contrast, and background subtraction, and contour masks were generated to delineate perinuclear and cytoplasmic surface areas, as described above. Mean fluorescence intensities over each perinuclear and cytoplasmic areas were collected using the "Measure" function after selecting each defined mask. Cytoplasmic-to-perinuclear signal intensity ratios were calculated for each cell by dividing cytoplasmic mean fluorescence intensity over perinuclear mean fluorescence intensity. Results were further normalized by dividing all cytoplasmic-to-perinuclear signal intensity ratios by the mean ratio value of the control.

Pigment content in brightfield images

Pigment content in primary human melanocytes was assessed by quantitative image analysis using ImageJ [v1.53j, (*17*)]. Briefly, bright field images were first denoised using a difference of Gaussians algorithm as follows: (i) images were transformed using the "Gaussian blur" function with a 30-pixel radius, and (ii) Gaussian-blurred duplicates were subtracted from original images using the "Image Calculator" function. Mean pixel value of selected background areas was then measured, and subtracted from denoised images using the "Subtract background" function. Mean pixel value for pigment content was then computed from each background-subtracted denoised images using the "Measure" function. To quantify area of pigment coverage, images were further converted into binary images using the "Threshold" function with the "Moments" method, and total pigment area was computed after inverting the images using the "Measure" function. Pigment content in each picture was expressed as the ratio of total pigment area over total cell area. Results were further normalized after dividing all pigment content values by the mean pigment content value of the control for each given experiment.

Proportion of ciliated cells, and length of primary cilia in ciliated cells

Number of ciliated cells, and primary cilium length among ciliated cells were quantified manually from IFM images of melanocytes stained for ARL13B (ciliary axoneme), and nuclei (DAPI) on ImageJ [v1.53j, (*17*)]. Briefly, the total number of cells ($DAPI^+$ nuclei), and total number of ciliated cells (i.e., expressing any $ARL13B^+$ projections) were counted manually. The percentage of ciliated cells was expressed as the ratio of number of ciliated cells over total number of cells across all quantified pictures for each experiment. Primary cilium length was measured in each $ARL13B^+$ cell using the line tool, and mean length was computed over all ciliated cells.

cAMP production in live cells expressing the CFP/YFP FRET biosensor EPAC-$S^{H187}$

Live cell imaging was performed using CFP excitation and collecting either CFP emission (CFP signal) or YFP emission (FRET signal). Changes in CFP/FRET emission ratio over time was assessed by quantitative image analysis on ImageJ [v1.53j, (*17*)]. Specifically, background signal was measured from an off-cell area in both CFP and FRET stacks, and subtracted from each stack slice using the "Subtract background" function. Regions of interest were defined manually with the "Polygon selection" tool to delineate the primary cilium, plasma membrane at the edges of the cells, or cytoplasm. Mean signal intensities were measured within these defined areas in both background-subtracted FRET and CFP stacks using the "Measure" function. CFP/FRET emission ratio was calculated and plotted for each time point as the ratio of background-subtracted CFP over FRET emission signal intensity.

Quantification of LAMP2 area surrounding melanin

Percentage of LAMP2-positive area surrounding melanin was quantified in fixed cells using a custom-made macro on ImageJ [v1.53j, (*17*)]. Briefly, IFM pictures were first deconvolved with the Microvolution deconvolution plug-in on ImageJ [v1.53j, (*17*)] using the least number of iterations to avoid creating artifacts. Single z-plane images were then generated from deconvolved IFM (dIFM) z-stack series, and contrast and brightness values associated with LAMP2 and bright field channels in a given experiment were adjusted identically for all images. Bright field channel was then pseudocolored using the "Image Inverter" and "Green" functions, and debris and small particles were filtered out using two iterations of the "Analyze particle" function with 0 to 0.4 circularity to remove debris, and 0- to 5-pixel size to remove small particles. A mask highlighting melanin granules was then generated using a third iteration of the "Analyze particle" function with 0.85 to 1 circularity, and converted to binary image using the "Make binary" function. This mask was duplicated, and the boundaries of each object in the duplicate mask were expanded by 1 pixel using the "Dilate" function. A mask of the expanded boundaries was then created using the "Image calculator" function by subtracting the original melanin mask from the duplicate mask with dilated objects, and melanin boundary selection was saved using both the "Create Selection" and "ROI manager" functions. The LAMP2 channel was then converted to binary image using the Bernson algorithm of the "Auto Local Threshold" function with a 30-pixel radius. The percentage of LAMP2-positive area ("%Area") in the melanin boundary selection was then computed using the "Measure" function. %Area measurements were collected for individual cells and averaged across experimental conditions for each experiment.

**Graphical presentation of data and statistical analyses**

At least two technical replicates of each testing conditions were used in every experiment, and sample sizes are indicated in each figure legend. Most data are presented as "super-plots" (*21*) in which values for individual cells within separate experiments are represented by small dots, mean values for each experimental repeat are depicted with bigger dots, and overall distribution of the entire data set is outlined with a box plot showing median, quartiles, maximum and minimum values. Bar plots were sometimes used instead of box plots, and show mean values, and standard error of the mean (SEM) across all experimental repeats. Colors and patterns were used to highlight separate experimental conditions. Data representation and statistical analyses were carried out in RStudio (version 2025.05.1+513) using open source packages including 'ggplot2' (*22*), 'ggbreak' (*23*), 'ggpattern' (*24*), 'emmeans' (*25*), and 'RVAideMemoire' (*26*). Most experiments were designed to compare the means of single continuous quantitative variables across three or more experimental conditions and repeats. Statistical analyses for these experiments were conducted using generalized linear mixed-effect models (GLMM) treating each experimental condition and repeat as fixed and random effects, respectively. Model fitting was tested through assessment of both independence and homoscedasticity of residuals using a residual plot, and adjusted when necessary using other distributions (e.g., Gaussian, Gamma, Binomial, Poisson, etc.), and/ or data transformations with logarithmic, square root, or inverse functions. ANOVAs were then used to determine statistical significance of any fixed or random effect in the fitted models. When applicable, Tukey's HSD post-hoc tests were conducted to further assess statistical significance among individual experimental conditions using the Kenward-Roger degrees-of-freedom method to correct for small sample bias. Comparison of means across two experimental conditions were carried out using paired t-tests. In some experiments, proportions were compared across various experimental conditions (e.g., proportion of ciliated cells). Statistical analyses for these experiments were conducted using contingency tables with the Fisher's exact test. Asterisks used to indicate statistical significance on plotted graphs (* $p<0.05$, ** $p<0.01$, *** $p<0.001$) highlight differences between control and experimental samples, unless otherwise indicated.

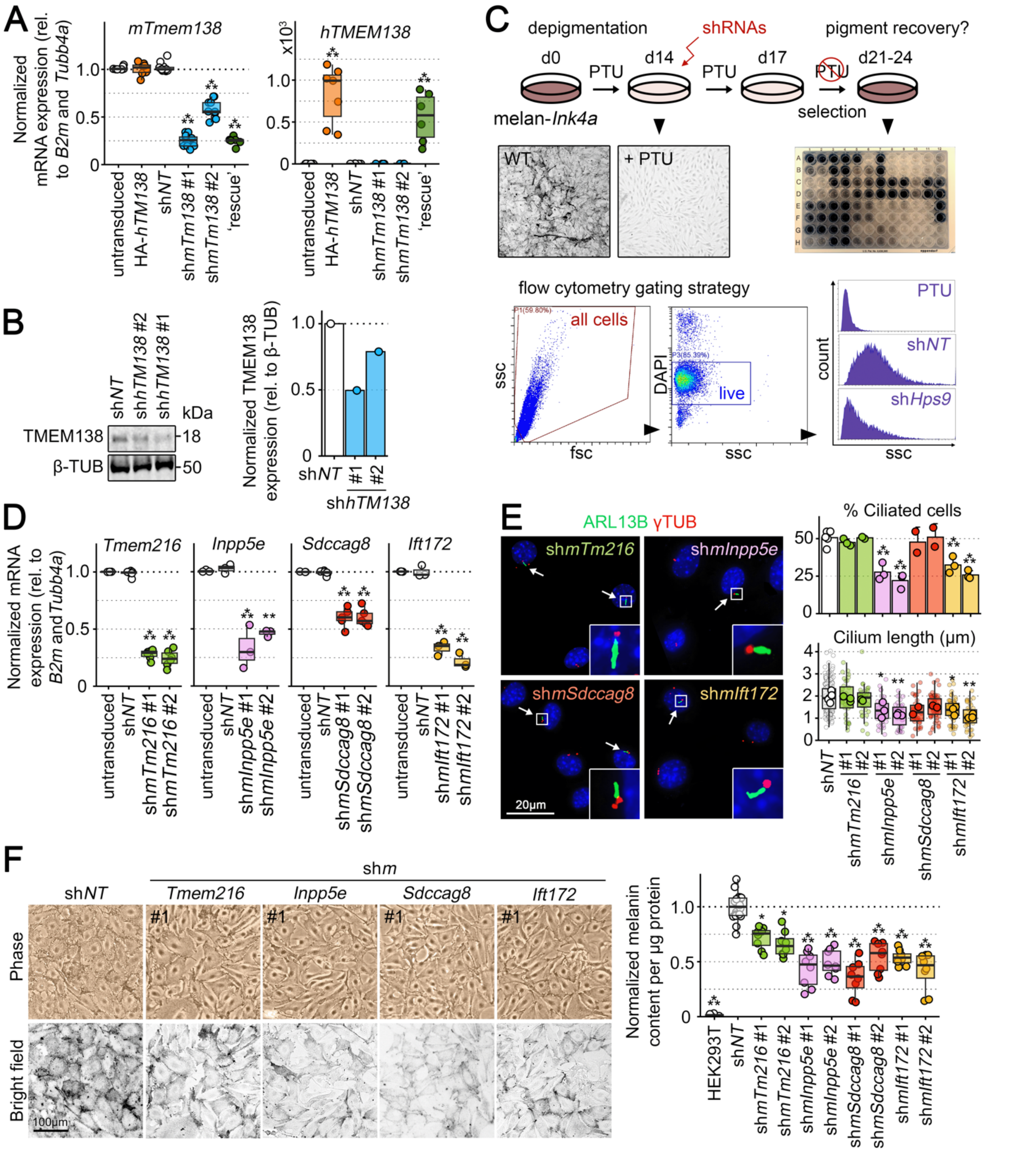


**Figure S1. Validation of target knockdowns and effects on cilium length and pigmentation.** **(A)** RT-qPCR assessment of mouse *Tmem138* (*mTm138*) and human *TMEM138* (*hTM138*) mRNA expression levels relative to *B2m* and *Tubb4a* housekeeping genes across 7-12 experiments in untransduced or stably transduced melan-*Ink4a* cells expressing HA-tagged human TMEM138, sh*NT*, sh*mTm38* #1 or #2, or sh*mTm38* #1 + HA-TMEM138 (rescue). Data are normalized to levels in untransduced cells. **(B)** Left, immunoblotting of TMEM138 relative to β-Tubulin (β-TUB) in stably transduced primary human keratinocytes expressing sh*NT* or sh*hTM38* #1 or #2. Positions of nearby molecular weight markers are indicated to the right in kDa. Right, quantification of TMEM138 band intensity relative to sh*NT* cells. Note that these shRNAs to *hTM138* are distinct from the ones targeting *mTm138*. **(C)** Melan-*Ink4a* cell preparation and gating strategy for shRNA screen by flow cytometry. Top, experimental strategy. Middle, examples of (left) fields of untransduced cells that were untreated (WT) or treated with phenylthiourea for 14 days (PTU) and (right) a 96-well plate from the screen. Bottom left, gating strategy for flow cytometry analysis of side scatter (ssc; fsc, forward scatter; DAPI, labeling with nuclear stain 4′,6-diamidino-2-phenylindole to exclude dead cells). Bottom right, examples of gated ssc profiles for cells treated throughout the 24-day period with phenylthiourea (PTU), and negative control (sh*NT*) and positive control (sh*Hps9*) cells after 7-day recovery without PTU. **(D)** Target mRNA expression by RT-qPCR relative to *B2m* and *Tubb4a* in melan-*Ink4a* cells stably expressing sh*NT* or either of two shRNAs to mouse *Tmem216* (sh*mTm216*), *Inpp5e* (sh*mInpp5e*), *Sdccag8* (sh*mSdccag8*), or *Ift172* (sh*mIft172*). Data are from 3-6 independent experiments and normalized to levels in untransduced cells. **(E)** Left, IFM imaging of primary cilia using antibodies against ARL13B (green) and γ-Tubulin (red). Arrows indicate ciliated cells. Insets represent 4-fold magnification of boxed region. Right, proportion of ciliated cells and cilium length among ciliated cells were quantified in 71-300 cells per condition across 2-3 experiments. **(F, G)** Representative images of phase and bright-field contrast microscopy (**F**) and quantitative melanin content assay across 6-18 experiments (**G**) of stable melan-Ink4a cells expressing sh*NT*, sh*mTm216*, sh*mInpp5e*, sh*mSdccag8*, or sh*mIft172*. Values shown in G are normalized to melanin content of sh*NT* cells. Statistics: Data were analyzed using Fisher's exact test (E, proportion of ciliated cells), or one-way ANOVA and Tukey's multiple comparison test (all other datasets). *, <0.05; **, p<0.01; ***, p<0.001.

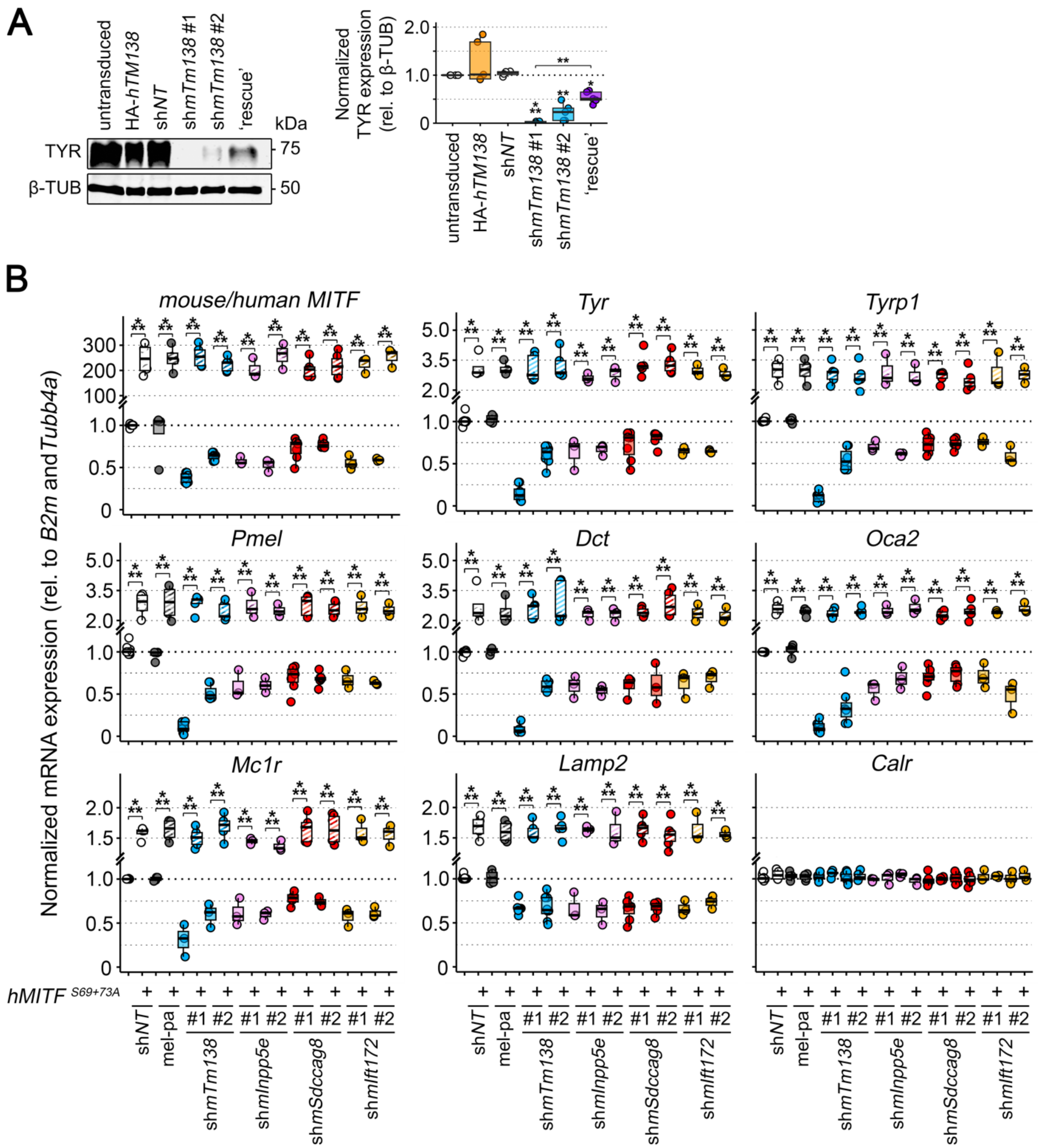


**Fig. S2. Effect of ciliary knockdown and/ or *MITF* expression on melanogenic protein or gene expression.** (**A**) Immunoblotting of TYR expression relative to β-Tubulin (β-TUB) in stable cell lines of melan-*Ink4a* melanocytes expressing nothing (untransduced), HA-tagged human TMEM138 (HA-hTM138), sh*NT*, sh*mTm138* #1 or #2, or sh*mTm138* #1 and HA-TMEM138 (rescue). Left, representative immunoblot, with positions of nearby molecular weight markers indicated to the right in kDa. Right, quantification of TYR band intensity over 4-5 experiments normalized to levels in untransduced cells. (**B**) RT-qPCR assessment of mRNA expression levels for indicated melanogenic genes, the lysosomal protein *Lamp2*, and the ER protein Calreticulin (*Calr*) relative to *B2m* and *Tubb4a* housekeeping genes across 3-8 experiments in melan-*Ink4a*

melanocytes expressing sh*NT* or either of two shRNAs to *mTm138* (sh*mTm138*), *mTmem216* (sh*mTm216*), *mInpp5e* (sh*mInpp5e*), *mSdccag8* (sh*mSdccag8*) or *mIft172* (sh*mIft172*). +, cells were additionally stably transduced with a lentivirus to express a constitutively active form of EGFP-tagged human *MITF* ($hMITF^{S69+73A}$). Untransduced cells and BLOC-1-deficient melan-pa (mel-pa) melanocytes were used as controls. Data are normalized to levels in cells expressing sh*NT* only. Statistics: Data were analyzed using one-way ANOVA and Tukey's multiple comparison test. *, p<0.05; **, p<0.01; ***, p<0.001.

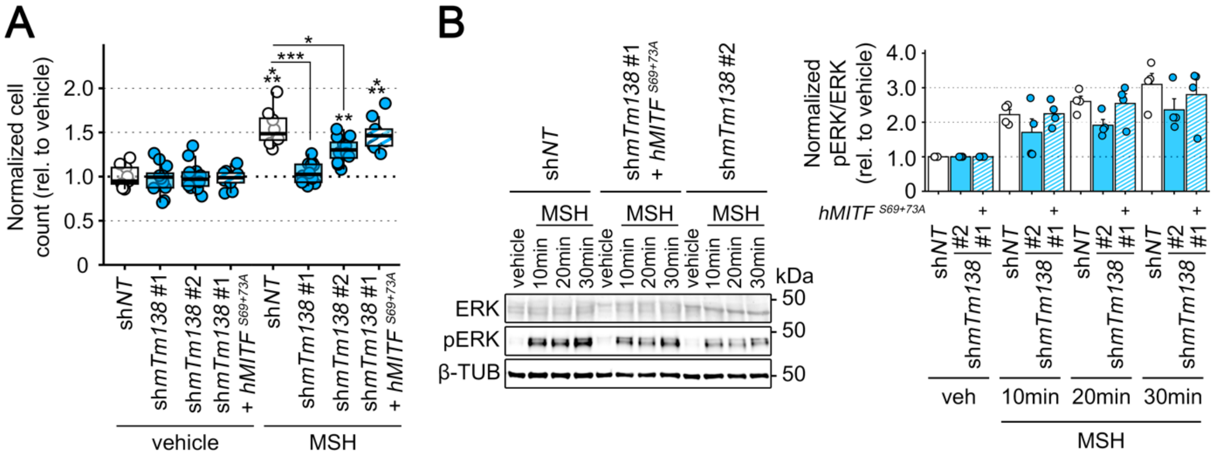


**Fig. S3. Effect of MSH stimulation on non-pigmentary functions of MC1R in ciliated and non-ciliated melanocytes.** (**A**) Melan-*Ink4a* melanocytes stably expressing sh*NT*, sh*mTm138* #1 or #2, or both sh*mTm138* #1 and constitutively active *hMITF*$^{S69+73A}$ were seeded at the same cell density and treated daily with vehicle or 100 pM [Nle$^4$, D-Phe$^7$]-α-MSH in serum-containing medium. After 6 days, cells were enzymatically detached and manually counted after Trypan blue exclusion. Data are presented as mean + SEM fold changes in cell counts after MSH stimulation relative to vehicle control across 7 independent experiments. (**B**) The same cell lines were stimulated with vehicle or MSH for 10, 20, or 30 min, then harvested for immunoblotting for total ERK, pERK, and β-Tubulin (β-TUB). Left, representative immunoblots, with the position of the 50 kDa molecular weight marker indicated. Right, quantification of pERK/ ERK ratio (normalized to vehicle control) over 4 experiments. Statistics: Data were analyzed using one-way ANOVA and Tukey's multiple comparison test. *, $p<0.05$; **, $p<0.01$; ***, $p<0.001$.

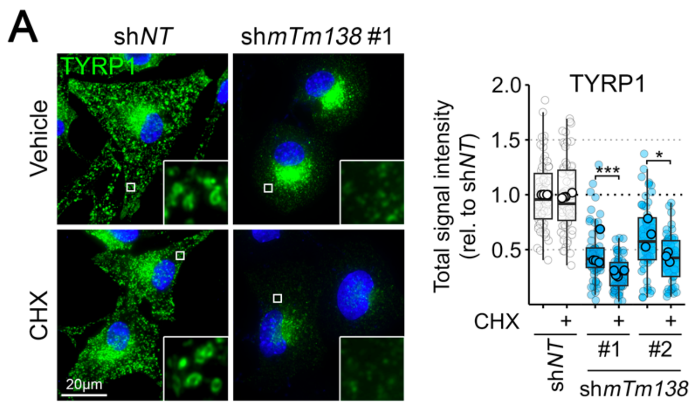


**Fig. S4. TYRP1 is unstable in melanocytes depleted of TMEM138.** (**A**) Melan-*Ink4a* cells expressing sh*NT*, or sh*mTm138* #1 or #2 were treated on coverslips for 3 h with vehicle or cycloheximide (CHX), then fixed, labeled for TYRP1, and analyzed by dIFM. Left, representative dIFM images; inset shows 7-fold magnification of boxed region. Right, TYRP1 signal intensity was quantified in 48-67 cells per condition across 3-4 experiments, and plotted relative to the mean intensity of sh*NT*-expressing cells. Statistics: Data were analyzed using one-way ANOVA and Tukey's multiple comparison test. *, $p<0.05$; ***, $p<0.001$.

### References and Notes


1. J. F. Berson, D. C. Harper, D. Tenza, G. Raposo, M. S. Marks, Pmel17 Initiates Premelanosome Morphogenesis within Multivesicular Bodies. *Mol. Biol. Cell* **12**, 3451–3464 (2001). doi:10.1091/mbc.12.11.3451.
2. J. F. Berson, D. W. Frank, P. A. Calvo, B. M. Bieler, M. S. Marks, "A Common Temperature-sensitive Allelic Form of Human Tyrosinase Is Retained in the Endoplasmic Reticulum at the Nonpermissive Temperature*" (2000); http://www.jbc.org.
3. Y. Feng, N. Xie, F. Inoue, S. Fan, J. Saskin, C. Zhang, F. Zhang, M. E. B. Hansen, T. Nyambo, S. W. Mpoloka, G. G. Mokone, C. Fokunang, G. Belay, A. K. Njamnshi, M. S. Marks, E. Oancea, N. Ahituv, S. A. Tishkoff, Integrative functional genomic analyses identify genetic variants influencing skin pigmentation in Africans. *Nat. Genet.* **56**, 258–272 (2024). doi:10.1038/s41588-023-01626-1; pmid:38200130.
4. S. R. G. Setty, D. Tenza, S. T. Truschel, E. Chou, E. V. Sviderskaya, A. C. Theos, M. L. Lamoreux, S. M. Di Pietro, M. Starcevic, D. C. Bennett, E. C. Dell'Angelica, G. Raposo, M. S. Marks, BLOC-1 is required for cargo-specific sorting from vacuolar early endosomes toward lysosome-related organelles. *Mol. Biol. Cell* **18**, 768–780 (2007). doi:10.1091/mbc.E06-12-1066; pmid:17182842.
5. M. Kovářová, P. Tolar, R. Arudchandran, L. Dráberová, J. Rivera, P. Dráber, Structure-Function Analysis of Lyn Kinase Association with Lipid Rafts and Initiation of Early Signaling Events after Fcε Receptor I Aggregation. *Mol. Cell. Biol.* **21**, 8318–8328 (2001). doi:10.1128/mcb.21.24.8318-8328.2001; pmid:11713268.
6. K. C. Ngeow, H. J. Friedrichsen, L. Li, Z. Zeng, S. Andrews, L. Volpon, H. Brunsdon, G. Berridge, S. Picaud, R. Fischer, R. Lisle, S. Knapp, P. Filippakopoulos, H. Knowles, E. Steingrímsson, K. L. B. Borden, E. E. Patton, C. R. Goding, BRAF/MAPK and GSK3 signaling converges to control MITF nuclear export. *Proc. Natl. Acad. Sci. U. S. A.* **115**, E8668–E8677 (2018). doi:10.1073/pnas.1810498115; pmid:30150413.
7. J. Moffat, D. A. Grueneberg, X. Yang, S. Y. Kim, A. M. Kloepfer, G. Hinkle, B. Piqani, T. M. Eisenhaure, B. Luo, J. K. Grenier, A. E. Carpenter, S. Y. Foo, S. A. Stewart, B. R. Stockwell, N. Hacohen, W. C. Hahn, E. S. Lander, D. M. Sabatini, D. E. Root, A Lentiviral RNAi Library for Human and Mouse Genes Applied to an Arrayed Viral High-Content Screen. *Cell* **124**, 1283–1298 (2006). doi:10.1016/j.cell.2006.01.040; pmid:16564017.
8. S. B. Frank, V. V. Schulz, C. K. Miranti, A streamlined method for the design and cloning of shRNAs into an optimized Dox-inducible lentiviral vector. *BMC Biotechnol.* **17**, 1–10 (2017). doi:10.1186/s12896-017-0341-x; pmid:28245848.
9. E. V. Sviderskaya, S. P. Hill, T. J. Evans-Whipp, L. Chin, S. J. Orlow, D. J. Easty, S. C. Cheong, D. Beach, R. A. DePinho, D. C. Bennett, p16Ink4a in melanocyte senescence and differentiation. *J. Natl. Cancer Inst.* **94**, 446–454 (2002). doi:10.1093/jnci/94.6.446; pmid:11904317.
10. S. Morita, T. Kojima, T. Kitamura, "Plat-E: an efficient and stable system for transient packaging of retroviruses" (2000); www.nature.com/gt.
11. P. Salmon, D. Trono, Production and Titration of Lentiviral Vectors. *Curr. Protoc. Neurosci.* **37** (2006). doi:10.1002/0471142301.ns0421s37.
12. B. Goyer, U. Pereira, B. Magne, D. Larouche, S. Kearns-Turcotte, P. J. Rochette, L. Martin, L. Germain, Impact of ultraviolet radiation on dermal and epidermal DNA damage in a human pigmented bilayered skin substitute. *J. Tissue Eng. Regen. Med.* **13**, 2300–2311 (2019). doi:10.1002/term.2959.

13. D. Larouche, L. Cantin-Warren, M. Desgagné, R. Guignard, I. Martel, A. Ayoub, A. Lavoie, R. Gauvin, F. A. Auger, V. J. Moulin, L. Germain, Improved Methods to Produce Tissue-Engineered Skin Substitutes Suitable for the Permanent Closure of Full-Thickness Skin Injuries. *Biores. Open Access* **5**, 320–329 (2016). doi:10.1089/biores.2016.0036.
14. R. Pouliot, D. Larouche, F. A. Auger, J. Juhasz, W. Xu, H. Li, L. Germain, Reconstructed human skin produced in vitro and grafted on athymic mice. *Transplantation* **73**, 1751–1757 (2002). doi:10.1097/00007890-200206150-00010.
15. F. A. Auger, M. Rémy-Zolghadri, G. Grenier, L. Germain, A truly new approach for tissue engineering: the LOEX self-assembly technique. *Ernst Schering Res. Found. Workshop*, 73–88 (2002). doi:10.1007/978-3-662-04816-0_6; pmid:11816275.
16. C. Delevoye, X. Heiligenstein, L. Ripoll, F. Gilles-Marsens, M. K. Dennis, R. A. Linares, L. Derman, A. Gokhale, E. Morel, V. Faundez, M. S. Marks, G. Raposo, BLOC-1 Brings Together the Actin and Microtubule Cytoskeletons to Generate Recycling Endosomes. *Current Biology* **26**, 1–13 (2016). doi:10.1016/j.cub.2015.11.020; pmid:26725201.
17. C. A. Schneider, W. S. Rasband, K. W. Eliceiri, NIH Image to ImageJ: 25 years of image analysis. [Preprint] (2012). https://doi.org/10.1038/nmeth.2089.
18. F. Giordano, C. Bonetti, E. M. Surace, V. Marigo, G. Raposo, The ocular albinism type 1 (OA1) G-protein-coupled receptor functions with MART-1 at early stages of melanogenesis to control melanosome identity and composition. *Hum. Mol. Genet.* **18**, 4530–4545 (2009). doi:10.1093/hmg/ddp415; pmid:19717472.
19. M. S. Marks, P. A. Roche, E. van Donselaar, L. Woodruff, P. J. Peters, J. S. Bonifacino, A lysosomal targeting signal in the cytoplasmic tail of the beta chain directs HLA-DM to MHC class II compartments. *J. Cell Biol.* **131**, 351–69 (1995). doi:10.1083/jcb.131.2.351; pmid:7593164.
20. Y. Zhu, S. Li, A. Jaume, R. A. Jani, C. Delevoye, G. Raposo, M. S. Marks, Type II phosphatidylinositol 4-kinases function sequentially in cargo delivery from early endosomes to melanosomes. *Journal of Cell Biology* **221** (2022). doi:10.1083/jcb.202110114; pmid:36169639.
21. S. J. Lord, K. B. Velle, R. Dyche Mullins, L. K. Fritz-Laylin, SuperPlots: Communicating reproducibility and variability in cell biology. Rockefeller University Press [Preprint] (2020). https://doi.org/10.1083/JCB.202001064.
22. Hadley. Wickham, *Ggplot2 : Elegant Graphics for Data Analysis* (Springer, New York, 2016; https://ggplot2.tidyverse.org).
23. S. Xu, M. Chen, T. Feng, L. Zhan, L. Zhou, G. Yu, Use ggbreak to Effectively Utilize Plotting Space to Deal With Large Datasets and Outliers. *Front. Genet.* **12** (2021). doi:10.3389/fgene.2021.774846.
24. M. FC, T. L. Davis, ggpattern: "ggplot2" Pattern Geoms. [Preprint] (2022). https://doi.org/10.32614/CRAN.package.ggpattern.
25. R. V. Lenth, J. Piaskowski, emmeans: Estimated Marginal Means, aka Least-Squares Means. [Preprint] (2017). https://doi.org/10.32614/CRAN.package.emmeans.
26. M. Hervé, RVAideMemoire: Testing and Plotting Procedures for Biostatistics. [Preprint] (2011). https://doi.org/10.32614/CRAN.package.RVAideMemoire.
