## Supplementary Tables for "Ciliary regulation of melanogenesis unveils moonlighting roles for Joubert Syndrome proteins"

#### Table S1

| List of primers for cloning | Forward/ Reverse | Sequence (5' to 3') |
|---|---|---|
| Primer 1 | Forward | GAGTCGACCCGGGCGGCCGCCACCATGTACCCATACGATGTTCCAGATTACGCTATGCTCCAGACCAG |
| Primer 2 | Reverse | GGGCGGAATTTACGTAGCGGCCGCTTTACCTTCGAACTTGCATGAACTCCTTG |
| Primer 3 | Forward | TTGTTTAATATTATTATCATTTTCCTCATGTTCTTCAACAC |
| Primer 4 | Reverse | GACTGCAATATCCTGGATGATGAA |
| Primer 5 | Forward | GATGATAATATGGCCACAACCATGGATGACCGAGTACAAGCCCACGGTG |
| Primer 6 | Reverse | ATTTTATCGTCGACCACTGTGCTGGTCAGGCACCGGGCTTGCG |
| Primer 7 | Forward | GAATTCCTGCAGGCCTGCCACCATGTTCAGTCTGATGGC |
| Primer 8 | Reverse | GGCGGAATTTACGTAGCGGCCTCACTTGTACAGCTCGT |
| Primer 9 | Forward | TACGGGAATTCCTGCAGGCCTCACTATAGGGAGACCCAAG |
| Primer 10 | Reverse | GGAATTTACGTAGCGGCCGTTAACCTCGAGGGAATTCGGCT |
| Primer 11 | Forward | CAAGCGCAAGGACAAGGATCCCATGGCGGCCGCAATGGTGAGCAAGGG |
| Primer 12 | Reverse | CTCTTGATGCAGCCCATGGTGGCGGCAAGCTTGGGTCTCCCTATA |
| Primer 13 | Forward | GATCTAGCTAGTTAATGCCACCATGTTCAGTCTGATGGCCAG |
| Primer 14 | Reverse | TATAGTGAGGCCGGAGCTTCCTGAGATCACATCATGAGCATCAC |
| Primer 15 | Forward | AAAAGCAGGCTGCCACCTTTGCCACCATGGTGTCTACAGGCGAGGA |
| Primer 16 | Reverse | CAACGGCAAAAAGAATGGCCATGAATTCGAACATAGACCCACCGCCTCCG |
| Primer 17 | Forward | ATTCGAGCTCGGTACCTTTAAGAC |
| Primer 18 | Reverse | GGTGGCGACCGGTGGATC |
| Primer 19 | Forward | ATATAGCGGCCGCTACGTAAATTCCG |
| Primer 20 | Reverse | GTTGGTTAATTAAGCTATTCCTTTGCCCTCGG |
| Primer 21 | Forward | TCTCTCCCCAGGGGGATCCACCTGCAGGCCACCATGGTGAGCAAGGGCGAGG |
| Primer 22 | Reverse | GCTCGAATGGTGGCGACCGGTCTAGACTAACACGCATGCTCCGTTTCTTCTG |

#### Table S2

| WT sequence for human TMEM138 | Modified sequence for human TMEM138 (sh*mTmem138* #1 resistant) |
|---|---|
| ATGCTCCAGACCAGTAACTACAGCCTGGTGCTCTCTCTGCAGTTCCT<br>GCTGCTGTCCTATGACCTCTTTGTCAATTCCTTCTCAGAACTGCTCCA<br>AAAGACTCCTGTCATCCAGCTTGTGCTCTTCATCATCCAGGATATTGC<br>AGTCCTCTTCAACATCATCATCATTTTCCTCATGTTCTTCAACACCTTC<br>GTCTTCCAGGCTGGCCTGGTCAACCTCCTATTCCATAAGTTCAAAGG<br>GACCATCATCCTGACAGCTGTGTACTTTGCCCTCAGCATCTCCCTTCA<br>TGTCTGGGTCATGAACTTACGCTGGAAAAACTCCAACAGCTTCATAT<br>GGACAGATGGACTTCAAATGCTGTTTGTATTCCAGAGACTAGCAGCA<br>GTGTTGTACTGCTACTTCTATAAACGGACAGCCGTAAGACTAGGCGA<br>TCCTCACTTCTACCAGGACTCTTTGTGGCTGCGCAAGGAGTTCATGC<br>AAGTTCGAAGG | ATGCTCCAGACCAGTAACTACAGCCTGGTGCTCTCTCTGCAGTTCCT<br>GCTGCTGTCCTATGACCTCTTTGTCAATTCCTTCTCAGAACTGCTCCA<br>AAAGACTCCTGTCATCCAGCTTGTGCTCTTCATCATCCAGGATATTGC<br>AGTCTTGTTTAATATTATTATCATTTTCCTCATGTTCTTCAACACCTTCG<br>TCTTCCAGGCTGGCCTGGTCAACCTCCTATTCCATAAGTTCAAAGGG<br>ACCATCATCCTGACAGCTGTGTACTTTGCCCTCAGCATCTCCCTTCAT<br>GTCTGGGTCATGAACTTACGCTGGAAAAACTCCAACAGCTTCATATG<br>GACAGATGGACTTCAAATGCTGTTTGTATTCCAGAGACTAGCAGCAG<br>TGTTGTACTGCTACTTCTATAAACGGACAGCCGTAAGACTAGGCGAT<br>CCTCACTTCTACCAGGACTCTTTGTGGCTGCGCAAGGAGTTCATGCA<br>AGTTCGAAGG |

#### Table S3

| species | GeneID | Symbol | RefSeq | TRC_ID | SenseSeq | Region | shRNA_ID |
|---|---|---|---|---|---|---|---|
| human | 51524 | TMEM138 | NM_016464 | TRCN0000148855 | CCATAAGTTCAAAGGGACCAT | CDS | |
| human | 51524 | TMEM138 | NM_016464 | TRCN0000149031 | GCTGTTTGTATTCCAGAGACT | CDS | |
| human | 51524 | TMEM138 | NM_016464 | TRCN0000146693 | CCTCCTATTCCATAAGTTCAA | CDS | sh*hTMEM138* #1 |
| human | 51524 | TMEM138 | NM_016464 | TRCN0000128381 | GATGGACTTCAAATGCTGTTT | CDS | sh*hTMEM138* #2 |
| human | 51524 | TMEM138 | NM_016464 | TRCN0000128999 | GCTGTGTTCAGCATTCAAGAA | 3UTR | |
| mouse | 12010 | B2M | NM_009735 | TRCN0000066423 | CGCCTCACATTGAAATCCAAA | CDS | |
| mouse | 12010 | B2M | NM_009735 | TRCN0000066424 | GCCGAACATACTGAACTGCTA | CDS | |
| mouse | 12010 | B2M | NM_009735 | TRCN0000066425 | GCAGAGTTAAGCATGCCAGTA | CDS | |
| mouse | 12010 | B2M | NM_009735 | TRCN0000066426 | CCAAATGCTGAAGAACGGGAA | CDS | |
| mouse | 12010 | B2M | NM_009735 | TRCN0000066427 | CGGCCTGTATGCTATCCAGAA | CDS | |
| mouse | 18457 | BLOC1S6/ HPS9 | NM_019788 | TRCN0000125574 | GCTTAAAGAAAGGCATAACTT | 3UTR | |
| mouse | 18457 | BLOC1S6/ HPS9 | NM_019788 | TRCN0000125575 | CAGAACCAAGTTGTGTTACTA | CDS | |
| mouse | 18457 | BLOC1S6/ HPS9 | NM_019788 | TRCN0000125576 | CATGTTGGACATCAATGCTTT | CDS | |
| mouse | 18457 | BLOC1S6/ HPS9 | NM_019788 | TRCN0000125577 | ACACAGAACCAAGTTGTGTTA | CDS | |
| mouse | 18457 | BLOC1S6/ HPS9 | NM_019788 | TRCN0000125578 | CTGTCCCACTACTTACCAGAT | CDS | |
| mouse | 18582 | PDE6D | NM_008801 | TRCN0000114881 | GCTTTCTTTAGCCTCTGGTTT | 3UTR | |
| mouse | 18582 | PDE6D | NM_008801 | TRCN0000114882 | CAATGTCATCATAGAGACAAA | CDS | |
| mouse | 18582 | PDE6D | NM_008801 | TRCN0000114883 | GAAGAGTGGTTCTTCGAGTTT | CDS | |
| mouse | 18582 | PDE6D | NM_008801 | TRCN0000114884 | TGTCATCATAGAGACAAAGTT | CDS | |
| mouse | 18582 | PDE6D | NM_008801 | TRCN0000114885 | GCTTCAAACTAAATTGGATGA | CDS | |
| mouse | 140859 | NPHP9/ NEK8 | NM_080849 | TRCN0000026956 | GCTGGATGAGACTCATCCTTA | CDS | |
| mouse | 140859 | NPHP9/ NEK8 | NM_080849 | TRCN0000026979 | CCCAACGTCATCGAGTACTAT | CDS | |
| mouse | 140859 | NPHP9/ NEK8 | NM_080849 | TRCN0000026978 | CCGAGGCATTATCATGACGTT | CDS | |
| mouse | 140859 | NPHP9/ NEK8 | NM_080849 | TRCN0000027000 | CCTTCTTGACAAACACCGCAT | CDS | |
| mouse | 140859 | NPHP9/ NEK8 | NM_080849 | TRCN0000027033 | CAGAGAGTTGTCTGTGGTATT | CDS | |
| mouse | 64436 | INPP5E | NM_033134 | TRCN0000080703 | GCCCAAGATCAAGATCATAAA | 3UTR | sh*mInpp5e* #1 |
| mouse | 64436 | INPP5E | NM_033134 | TRCN0000080704 | CCATCCTACAAGTTTGACATT | CDS | |
| mouse | 64436 | INPP5E | NM_033134 | TRCN0000080705 | CGGCAAGTTTGACCGAGAATT | CDS | |
| mouse | 64436 | INPP5E | NM_033134 | TRCN0000080706 | GAACCCACAAACTAAGGCAAA | CDS | |

| | | | | | | | |
|---|---|---|---|---|---|---|---|
| mouse | 64436 | INPP5E | NM_033134 | TRCN0000080707 | CCTTGTGGAGTGACTGTCTTT | CDS | sh*mInpp5e* #2 |
| mouse | 72507 | DZIP1L | NM_028258 | TRCN0000173899 | GCCAGGCATAGTAATCGAGTA | 3UTR | |
| mouse | 72507 | DZIP1L | NM_028258 | TRCN0000174038 | CAGCGGGAAATGGAAGCTAAA | CDS | |
| mouse | 72507 | DZIP1L | NM_028258 | TRCN0000173436 | CCACGTTAGAAGAGAAACTGA | CDS | |
| mouse | 72507 | DZIP1L | NM_028258 | TRCN0000176437 | CAATATGTTCTTAAGGCCCAA | CDS | |
| mouse | 72507 | DZIP1L | NM_028258 | TRCN0000173781 | CAAGAGGGACACAAAGGGAAT | CDS | |
| mouse | 93730 | BBS17/ LZTFL1 | NM_033322 | TRCN0000084323 | CCATGCAAGTACAGTGAGATT | 3UTR | |
| mouse | 93730 | BBS17/ LZTFL1 | NM_033322 | TRCN0000084324 | CCATTGAAATACAGGCTGTAA | CDS | |
| mouse | 93730 | BBS17/ LZTFL1 | NM_033322 | TRCN0000084325 | GCCTAAACTTGTTCCAATTAA | CDS | |
| mouse | 93730 | BBS17/ LZTFL1 | NM_033322 | TRCN0000084326 | GCAACTCTTAGGAGTGAATTT | CDS | |
| mouse | 93730 | BBS17/ LZTFL1 | NM_033322 | TRCN0000084327 | CATTGAAATACAGGCTGTAAA | CDS | |
| mouse | 67694 | IFT74/ BBS22 | NM_026319 | TRCN0000113225 | GCCGACTGAGTCTACAAGCAT | 3UTR | |
| mouse | 67694 | IFT74/ BBS22 | NM_026319 | TRCN0000113226 | CCTCAGATTATGATACCCTTA | CDS | |
| mouse | 67694 | IFT74/ BBS22 | NM_026319 | TRCN0000113227 | GCCAGCATCATCACACTGTTA | CDS | |
| mouse | 67694 | IFT74/ BBS22 | NM_026319 | TRCN0000113228 | CGAGATCAAATGATTGCAGAA | CDS | |
| mouse | 67694 | IFT74/ BBS22 | NM_026319 | TRCN0000113229 | GAGGCTGTATTGCTGTATGAA | CDS | |
| mouse | 67661 | IFT172/ BBS20/ NPHP17 | NM_026298 | TRCN0000079813 | GCGGCCATCAACCACTATATT | CDS | |
| mouse | 67661 | IFT172/ BBS20/ NPHP17 | NM_026298 | TRCN0000079814 | GCTGCTGATCTCTCATTACTA | CDS | sh*mIft172* #1 |
| mouse | 67661 | IFT172/ BBS20/ NPHP17 | NM_026298 | TRCN0000079815 | GCTTATGTGTATGGTACAATA | CDS | sh*mIft172* #2 |
| mouse | 67661 | IFT172/ BBS20/ NPHP17 | NM_026298 | TRCN0000079816 | CCAGTGGAAGAAGGCAATTTA | CDS | |
| mouse | 67661 | IFT172/ BBS20/ NPHP17 | NM_026298 | TRCN0000079817 | CGGTTCTTGGATCTGACTGAT | CDS | |
| mouse | 24069 | SUFU/ BCNS2 | NM_016169 | TRCN0000089053 | CCTGAACTTTATTGTCCTTTA | 3UTR | |
| mouse | 24069 | SUFU/ BCNS2 | NM_016169 | TRCN0000089054 | GCACTTCACCTACAAGAGTAT | CDS | |
| mouse | 24069 | SUFU/ BCNS2 | NM_016169 | TRCN0000089055 | GCATTGAGACAGACGGTTCTA | CDS | |
| mouse | 24069 | SUFU/ BCNS2 | NM_016169 | TRCN0000089056 | CGTTTCGTCTGAAGAGAGAAA | CDS | |
| mouse | 24069 | SUFU/ BCNS2 | NM_016169 | TRCN0000089057 | GAGGAATTTAAACTTCCCAAA | CDS | |
| mouse | 69807 | BBS11/ TRIM32 | NM_053084 | TRCN0000040828 | GCCGCAAGGAAATTCTCCATT | CDS | |
| mouse | 69807 | BBS11/ TRIM32 | NM_053084 | TRCN0000040829 | CCCAAGTTTGTCACCTGTGAT | CDS | |
| mouse | 69807 | BBS11/ TRIM32 | NM_053084 | TRCN0000040830 | GCATTGATAGCTTCGTGCTAA | CDS | |
| mouse | 69807 | BBS11/ TRIM32 | NM_053084 | TRCN0000040831 | GCTAGAATGTCCCATCTGCAT | CDS | |
| mouse | 69807 | BBS11/ TRIM32 | NM_053084 | TRCN0000040832 | GCTATCATCTGAGAAGATATT | CDS | |
| mouse | 16569 | KIF3B | NM_008444 | TRCN0000091378 | CCTCCCTTCTTGTTAAACAAT | 3UTR | |

| | | | | | | | |
|---|---|---|---|---|---|---|---|
| mouse | 16569 | KIF3B | NM_008444 | TRCN0000091379 | GCAGGGTTTCAATGGCACAAT | CDS | |
| mouse | 16569 | KIF3B | NM_008444 | TRCN0000091380 | CCATTGGAAATTACATCCTAT | CDS | |
| mouse | 16569 | KIF3B | NM_008444 | TRCN0000091381 | CGGGTAGGAAAGCTGAATCTT | CDS | |
| mouse | 16569 | KIF3B | NM_008444 | TRCN0000091382 | CGGTGCTACAAACATGAATGA | CDS | |
| mouse | 76816 | SDCCAG8/ BBS16 | NM_006642 | TRCN0000250548 | TCATCTGATTTGTACTTAATG | 3UTR | |
| mouse | 76816 | SDCCAG8/ BBS16 | NM_006642 | TRCN0000250551 | ATGCGCTTTCAGTTGAATAAA | CDS | sh*mSdccag8* #1 |
| mouse | 76816 | SDCCAG8/ BBS16 | NM_006642 | TRCN0000177945 | GCAGTTGCTTAACAAGCAGAA | CDS | sh*mSdccag8* #2 |
| mouse | 76816 | SDCCAG8/ BBS16 | NM_006642 | TRCN0000250547 | ACAACCTCGTTCCTATCATTA | CDS | |
| mouse | 19726 | RFX3 | NM_011265 | TRCN0000084613 | CTGAAGGTTGACTGTTGTCAA | 3UTR | |
| mouse | 19726 | RFX3 | NM_011265 | TRCN0000084614 | GCGAAGATACACATCGCTCAA | CDS | |
| mouse | 19726 | RFX3 | NM_011265 | TRCN0000084615 | CCGAACAACAACTTATCCCTA | CDS | |
| mouse | 19726 | RFX3 | NM_011265 | TRCN0000084616 | CGTCACAAGTACACAGACCAT | CDS | |
| mouse | 19726 | RFX3 | NM_011265 | TRCN0000084617 | CGTTGTTGTGAATCTTCAGTT | CDS | |
| mouse | 264134 | IFT56/ TTC26 | NM_153600 | TRCN0000192659 | GCTTGTTTGTATGCTCTGTAA | 3UTR | |
| mouse | 264134 | IFT56/ TTC26 | NM_153600 | TRCN0000192948 | GACGACACTAACTTATGGATT | CDS | |
| mouse | 264134 | IFT56/ TTC26 | NM_153600 | TRCN0000190598 | GCTTCCTGAAAGACACTCATT | 3UTR | |
| mouse | 264134 | IFT56/ TTC26 | NM_153600 | TRCN0000192807 | GCTCGCTGCTATATTATGAAT | CDS | |
| mouse | 94187 | NPHP14/ ZNF423 | NM_033327 | TRCN0000084708 | CCCTGAATGTAACGTGAAGTT | CDS | |
| mouse | 94187 | NPHP14/ ZNF423 | NM_033327 | TRCN0000084709 | CGGTGCATTACATGACTACAT | CDS | |
| mouse | 94187 | NPHP14/ ZNF423 | NM_033327 | TRCN0000084710 | CGCAGATGATAGGAGATGGTT | CDS | |
| mouse | 94187 | NPHP14/ ZNF423 | NM_033327 | TRCN0000084711 | CGACCTCAAGTTCTCCAACTT | CDS | |
| mouse | 94187 | NPHP14/ ZNF423 | NM_033327 | TRCN0000084712 | CGTGGAAGATGAGTCAATTTA | CDS | |
| mouse | 231214 | MKS06/ CC2D2A | NM_172274 | TRCN0000181757 | GACCCGAAACTAGATGAGGAT | CDS | |
| mouse | 231214 | MKS06/ CC2D2A | NM_172274 | TRCN0000197491 | CCCATTTAGCACCATCTATTT | CDS | |
| mouse | 231214 | MKS06/ CC2D2A | NM_172274 | TRCN0000197895 | GCTATGCAATTACTTTCTGTT | CDS | |
| mouse | 231214 | MKS06/ CC2D2A | NM_172274 | TRCN0000178278 | GTTCCGTGATTCTGTACGAAA | CDS | |
| mouse | 231214 | MKS06/ CC2D2A | NM_172274 | TRCN0000177905 | CCAAGATTCCTGGAAGATGAA | CDS | |
| mouse | 74682 | IFT121/ WDR35 | NM_172470 | TRCN0000103510 | GCCAAGGTTACCAGTTGGTTA | 3UTR | |
| mouse | 74682 | IFT121/ WDR35 | NM_172470 | TRCN0000103511 | CCCAATAATGTGAAGCTGAAA | CDS | |
| mouse | 74682 | IFT121/ WDR35 | NM_172470 | TRCN0000103512 | CCTGGCAATTTGCTTTGACAA | CDS | |
| mouse | 74682 | IFT121/ WDR35 | NM_172470 | TRCN0000103513 | GCTGAAGTTATTGCCTACTTT | CDS | |
| mouse | 74682 | IFT121/ WDR35 | NM_172470 | TRCN0000103514 | GCTGGAGTCTTGACTTTCTTT | CDS | |

| | | | | | | |
|---|---|---|---|---|---|---|
| mouse | 66279 | TMEM218 | NM_001310096 | TRCN0000124829 | GCGGTTATTGACCATTCCTTA | 3UTR |
| mouse | 66279 | TMEM218 | NM_001310096 | TRCN0000124830 | CCTGCTTCTTACTCATCATTT | CDS |
| mouse | 66279 | TMEM218 | NM_001310096 | TRCN0000124832 | GCTAGGTTCTCCATCGTCTTT | CDS |
| mouse | 66279 | TMEM218 | NM_001310096 | TRCN0000124833 | CTGGAGCCAATCTATGCCAAA | CDS |
| mouse | 106021 | TOPORS | NM_134097 | TRCN0000099110 | CCATTTCTTAGACTGAAGTAA | 3UTR |
| mouse | 106021 | TOPORS | NM_134097 | TRCN0000099111 | GCGGGTATTGAATGTAGCAAT | CDS |
| mouse | 106021 | TOPORS | NM_134097 | TRCN0000099112 | GCTGAGTTCTTCCGTAGAAAT | CDS |
| mouse | 106021 | TOPORS | NM_134097 | TRCN0000099113 | CGGAACTTGTTGAACTGTCTT | CDS |
| mouse | 106021 | TOPORS | NM_134097 | TRCN0000099114 | GCTTGCCTTCACAGATTAGTT | CDS |
| mouse | 52906 | AHI1 | NM_026203 | TRCN0000192100 | CGAGCCGATTCTTCTTTATAT | CDS |
| mouse | 52906 | AHI1 | NM_026203 | TRCN0000201363 | GCTGAAATGTTGAAACGCTAT | CDS |
| mouse | 52906 | AHI1 | NM_026203 | TRCN0000190390 | CCCTATTTGCTTCGAGAGTTT | CDS |
| mouse | 52906 | AHI1 | NM_026203 | TRCN0000202060 | CCAGCCGATCAGATGAACTAA | CDS |
| mouse | 52906 | AHI1 | NM_026203 | TRCN0000191207 | CCGAGTGTATTTCAAAGATAA | CDS |
| mouse | 16348 | NPHP2/ inversin | NM_010569 | TRCN0000086439 | GCCATTAAACTACTGCTAGAT | CDS |
| mouse | 16348 | NPHP2/ inversin | NM_010569 | TRCN0000086440 | CCTAATCAGATGGAGAACAAT | CDS |
| mouse | 16348 | NPHP2/ inversin | NM_010569 | TRCN0000086441 | CGGAGGGTACATCAACTGTAT | CDS |
| mouse | 16348 | NPHP2/ inversin | NM_010569 | TRCN0000086442 | GCAGTCTATACATCTTGACAA | CDS |
| mouse | 56297 | BBS03/ ARL6 | NM_019665 | TRCN0000100840 | GCCATCTCAATATCCTATCAT | 3UTR |
| mouse | 56297 | BBS03/ ARL6 | NM_019665 | TRCN0000100841 | CCAACGCTCAATCTCAAGATA | CDS |
| mouse | 56297 | BBS03/ ARL6 | NM_019665 | TRCN0000100842 | CAACGCTCAATCTCAAGATAT | CDS |
| mouse | 56297 | BBS03/ ARL6 | NM_019665 | TRCN0000100844 | CCAGATATTAAGCACCGTCGA | CDS |
| mouse | 234740 | TMEM231 | NM_001033321 | TRCN0000179219 | GCTTCTAACAACTGGTTCTTT | 3UTR |
| mouse | 234740 | TMEM231 | NM_001033321 | TRCN0000180945 | GAGATACGGAAAGAGCACTTA | CDS |
| mouse | 234740 | TMEM231 | NM_001033321 | TRCN0000179255 | GCCCATTCAGAACTTTGATTT | 3UTR |
| mouse | 234740 | TMEM231 | NM_001033321 | TRCN0000181082 | CAGGATGTCAACATTCGAGAT | CDS |
| mouse | 83396 | NPHP7/ GLIS2 | NM_031184 | TRCN0000082233 | GCCAAGTGTAACCAGCTCTTT | CDS |
| mouse | 83396 | NPHP7/ GLIS2 | NM_031184 | TRCN0000082234 | GCCAGGTACAAGATGCTCATT | CDS |
| mouse | 83396 | NPHP7/ GLIS2 | NM_031184 | TRCN0000082235 | CCTAAAGCTGAGCATCACCAA | CDS |
| mouse | 83396 | NPHP7/ GLIS2 | NM_031184 | TRCN0000082236 | GACCATCATGTCAAGCCTGAA | CDS |
| mouse | 26370 | CETN2 | NM_019405 | TRCN0000090949 | GCTTTCAAGTTGTTCGATGAT | CDS |
| mouse | 26370 | CETN2 | NM_019405 | TRCN0000090950 | GCCCAAGAAAGAAGAAATTAA | CDS |

| | | | | | | | |
|---|---|---|---|---|---|---|---|
| mouse | 26370 | CETN2 | NM_019405 | TRCN0000090951 | ACTGGAACTATAGACATCAAA | CDS | |
| mouse | 26370 | CETN2 | NM_019405 | TRCN0000090952 | GAACTTACTGAAGACCAGAAA | CDS | |
| mouse | 225523 | CEP120/ CCDC100 | NM_178686 | TRCN0000178907 | CGATGTTACAAACGAGCCTTT | CDS | |
| mouse | 225523 | CEP120/ CCDC100 | NM_178686 | TRCN0000179007 | CGAAGGACATTGTTGCAGTAT | CDS | |
| mouse | 225523 | CEP120/ CCDC100 | NM_178686 | TRCN0000179082 | CCAGAATTTGCTACTGAGCTA | CDS | |
| mouse | 225523 | CEP120/ CCDC100 | NM_178686 | TRCN0000179546 | GAAGGGCAGTACAGAGATTAA | CDS | |
| mouse | 245866 | IFT52 | NM_001356522 | TRCN0000175165 | GCTCATCTGGAACATGATATT | 3UTR | |
| mouse | 245866 | IFT52 | NM_001356522 | TRCN0000175987 | GCTGTTTGTGTCAGAACCTAT | 3UTR | |
| mouse | 245866 | IFT52 | NM_001356522 | TRCN0000174785 | GAAATCACTTCTGAGAAGTTA | CDS | |
| mouse | 245866 | IFT52 | NM_001356522 | TRCN0000175444 | CTTCCAAAGGATCAACAGGAT | CDS | |
| mouse | 245866 | IFT52 | NM_001356522 | TRCN0000173408 | GTTTACCACCAACACTGGCTA | CDS | |
| mouse | 16559 | KIF17 | NM_010623 | TRCN0000090873 | CCGAAGCGACAGTGAGAATAT | CDS | |
| mouse | 16559 | KIF17 | NM_010623 | TRCN0000090874 | CGCCTACTACATAGAACACTT | CDS | |
| mouse | 16559 | KIF17 | NM_010623 | TRCN0000090875 | CCCGCAGACAACAATTACGAT | CDS | |
| mouse | 16559 | KIF17 | NM_010623 | TRCN0000090876 | CCCAGATCCCATCATACTGAA | CDS | |
| mouse | 16559 | KIF17 | NM_010623 | TRCN0000090877 | GCCTGATGTAAACCTGAGAGT | CDS | |
| mouse | 241950 | BBS12 | NM_001008502 | TRCN0000178910 | CCTTTCCGAGTGATTCTCATT | CDS | |
| mouse | 241950 | BBS12 | NM_001008502 | TRCN0000179579 | GCTGTATAAGAACAGCAAGCA | 3UTR | |
| mouse | 241950 | BBS12 | NM_001008502 | TRCN0000179253 | GCATTGGATGTAGTGCTCTTA | CDS | |
| mouse | 241950 | BBS12 | NM_001008502 | TRCN0000183292 | GCTAGGACATTCATTCAACAA | CDS | |
| mouse | 68146 | ARL13B | NM_026577 | TRCN0000100500 | GCAATGAACATACACTGGTTT | 3UTR | |
| mouse | 68146 | ARL13B | NM_026577 | TRCN0000100501 | CGGCCTTGATAATGCTGGTAA | CDS | |
| mouse | 68146 | ARL13B | NM_026577 | TRCN0000100502 | GCAAAGGACTTTGATGCCTTA | CDS | |
| mouse | 68146 | ARL13B | NM_026577 | TRCN0000100503 | CAGTCAATACAGACGAGTCTA | CDS | |
| mouse | 68146 | ARL13B | NM_026577 | TRCN0000100504 | CCTGTCAGAAAGGTGACACTT | CDS | |
| mouse | 68642 | TMEM216 | NM_001277860 | TRCN0000249269 | GAAACCTCTGCCAGCGAAAGA | CDS | sh*mTmem216* #1 |
| mouse | 68642 | TMEM216 | NM_001277860 | TRCN0000249270 | CTCAGAGCTGCTGCTTGAGAT | CDS | |
| mouse | 68642 | TMEM216 | NM_001277860 | TRCN0000249268 | CGCTATGATGGCTTCCTATTA | CDS | sh*mTmem216* #2 |
| mouse | 68642 | TMEM216 | NM_001277860 | TRCN0000249272 | GAAGCTATCATGAACAGTATC | CDS | |
| mouse | 329795 | MKS03/ TMEM67 | NM_177861 | TRCN0000125984 | CAGAACTTAAGGACTCAGTAT | 3UTR | |
| mouse | 329795 | MKS03/ TMEM67 | NM_177861 | TRCN0000125985 | CCAGTGTTAAACCTAAATCTT | CDS | |
| mouse | 329795 | MKS03/ TMEM67 | NM_177861 | TRCN0000125986 | GCGAACCAACATTTGTGAATA | CDS | |

| | | | | | | |
|---|---|---|---|---|---|---|
| mouse | 329795 | MKS03/ TMEM67 | NM_177861 | TRCN0000125987 | CCTGTGTATGTCTACCAGGAT | CDS |
| mouse | 329795 | MKS03/ TMEM67 | NM_177861 | TRCN0000125988 | CGTGATGAACATGAACTCTTA | CDS |
| mouse | 236266 | ALMS1 | NM_145223 | TRCN0000180565 | GCCAGGACAAACTAACTTCTA | CDS |
| mouse | 236266 | ALMS1 | NM_145223 | TRCN0000183730 | CCAATGTCAGTACAAATGTTA | CDS |
| mouse | 236266 | ALMS1 | NM_145223 | TRCN0000183842 | CCAGACACTAAATCCATTAAA | CDS |
| mouse | 236266 | ALMS1 | NM_145223 | TRCN0000183191 | GCTATGGTAGTACAGATTCAT | CDS |
| mouse | 76411 | IFT43 | NM_001199843 | TRCN0000202015 | CAGGAGCATCACTGAGCATTT | 3UTR |
| mouse | 76411 | IFT43 | NM_001199843 | TRCN0000190841 | GAGCATCACTGAGCATTTGAA | 3UTR |
| mouse | 76411 | IFT43 | NM_001199843 | TRCN0000201136 | CTCAGGCTGAGAATTACTTAA | CDS |
| mouse | 76411 | IFT43 | NM_001199843 | TRCN0000189622 | CGACGACGATATTCCTGTGAT | CDS |
| mouse | 244585 | MKS05/ NPHP8/ RPGRIP1L | NM_173431 | TRCN0000106075 | CGCCTCTGATTAGCCTCATTA | 3UTR |
| mouse | 244585 | MKS05/ NPHP8/ RPGRIP1L | NM_173431 | TRCN0000106076 | CGGGACAATGTAGAAACGATT | CDS |
| mouse | 244585 | MKS05/ NPHP8/ RPGRIP1L | NM_173431 | TRCN0000106077 | CGGTGAAAGATACAGGTCTAA | CDS |
| mouse | 244585 | MKS05/ NPHP8/ RPGRIP1L | NM_173431 | TRCN0000106078 | GCAGGAACTATCCAAGTTATA | CDS |
| mouse | 244585 | MKS05/ NPHP8/ RPGRIP1L | NM_173431 | TRCN0000106079 | CGGCTAGTTAATGACAAGAAA | CDS |
| mouse | 237222 | OFD1 | NM_177429 | TRCN0000191532 | GCTAGAATCTTTAGAGACAAA | CDS |
| mouse | 237222 | OFD1 | NM_177429 | TRCN0000192215 | CGCTCAATTTGCAGATGGTTT | CDS |
| mouse | 237222 | OFD1 | NM_177429 | TRCN0000191531 | GAACTTACTATCAAAGAGCTA | CDS |
| mouse | 237222 | OFD1 | NM_177429 | TRCN0000191237 | CTTTCAAATGTGGACAAGCAA | CDS |
| mouse | 237222 | OFD1 | NM_177429 | TRCN0000191091 | CTACTTAAACTGCAACTGGAA | CDS |
| mouse | 21821 | IFT88/ TTC10 | NM_009376 | TRCN0000182620 | GCCCTCAGATAGAAAGACCAA | CDS |
| mouse | 21821 | IFT88/ TTC10 | NM_009376 | TRCN0000177272 | GCCAAATAAGTCATTTACCAA | 3UTR |
| mouse | 21821 | IFT88/ TTC10 | NM_009376 | TRCN0000176589 | CCATAAAGAAATGAGGATCAA | CDS |
| mouse | 21821 | IFT88/ TTC10 | NM_009376 | TRCN0000178064 | GCAGGAAGACTGAAAGTGAAT | CDS |
| mouse | 16576 | KIF7 | NM_001291222 | TRCN0000090438 | GCCCAGAGTTCTGGAGATAAT | 3UTR |
| mouse | 16576 | KIF7 | NM_001291222 | TRCN0000090439 | CGAGTACAAGAATGAGGCTAT | CDS |
| mouse | 16576 | KIF7 | NM_001291222 | TRCN0000090440 | CCTGCGTGCATTCTCATCAAA | CDS |
| mouse | 16576 | KIF7 | NM_001291222 | TRCN0000090441 | CGTGCTCAGAACATCCGCAAT | CDS |
| mouse | 27078 | MKS09/ B9d1 | NM_013717 | TRCN0000198821 | CCGGATGTGTTTGGGAATGAT | CDS |
| mouse | 27078 | MKS09/ B9d1 | NM_013717 | TRCN0000198787 | CCTCTTCAATGTGGTGACCAA | CDS |
| mouse | 27078 | MKS09/ B9d1 | NM_013717 | TRCN0000198329 | GCATCTAAGAGCCAAGATGTA | CDS |
| mouse | 27078 | MKS09/ B9d1 | NM_013717 | TRCN0000182008 | CCATGTTTGTGCCAGAGTCTA | CDS |

| | | | | | | |
|---|---|---|---|---|---|---|
| mouse | 27078 | MKS09/ B9d1 | NM_013717 | TRCN0000177277 | GATCTCACAGATAGCATCTAA | CDS |
| mouse | 68259 | IFT80/ WDR56 | NM_026641 | TRCN0000191159 | CCTCAGACTTAGAGTCTAAAT | 3UTR |
| mouse | 68259 | IFT80/ WDR56 | NM_026641 | TRCN0000190186 | CGGCAGTGATTAGTCTCTCAT | 3UTR |
| mouse | 68259 | IFT80/ WDR56 | NM_026641 | TRCN0000191678 | GCTCTTAGAATTGGTGCATTA | 3UTR |
| mouse | 68259 | IFT80/ WDR56 | NM_026641 | TRCN0000190083 | GCTCGCTTATCGTCAGAAGTT | CDS |
| mouse | 68259 | IFT80/ WDR56 | NM_026641 | TRCN0000193055 | CCAAGTTAGGAAGAGTGGAAA | CDS |
| mouse | 319845 | BBS09/ PTHB1 | NM_178415 | TRCN0000182387 | GCCTTTAGTCACATGCGCTTA | 3UTR |
| mouse | 319845 | BBS09/ PTHB1 | NM_178415 | TRCN0000182069 | CCTTTAGTCACATGCGCTTAT | 3UTR |
| mouse | 319845 | BBS09/ PTHB1 | NM_178415 | TRCN0000178683 | CTTTAGTCACATGCGCTTATA | 3UTR |
| mouse | 319845 | BBS09/ PTHB1 | NM_178415 | TRCN0000182647 | GCTGTGTGATAGGTTAGCCAA | CDS |
| mouse | 319845 | BBS09/ PTHB1 | NM_178415 | TRCN0000181485 | CCACTACAGTTGACTTGTGAT | CDS |
| mouse | 16568 | KIF3A | NM_008443 | TRCN0000090403 | CCCTGGACATACAGCTTACAA | 3UTR |
| mouse | 16568 | KIF3A | NM_008443 | TRCN0000090404 | CGCCATCTTTACAATTACTAT | CDS |
| mouse | 16568 | KIF3A | NM_008443 | TRCN0000090405 | GCGAAGAAAGCGTTCTGCAAA | CDS |
| mouse | 16568 | KIF3A | NM_008443 | TRCN0000090406 | CCAAAGACATTTACTTTCGAT | CDS |
| mouse | 16568 | KIF3A | NM_008443 | TRCN0000090407 | TCCGCCAGTTTCAGAAAGAAA | CDS |
| mouse | 67590 | TCTN3 | NM_026260 | TRCN0000180953 | GCTGCCATTCGATTTCTTCTT | CDS |
| mouse | 67590 | TCTN3 | NM_026260 | TRCN0000180137 | CCAAACAGTCAATGCAACCAA | CDS |
| mouse | 67590 | TCTN3 | NM_026260 | TRCN0000179139 | GCCTACTTATCTTGGCTAAAT | 3UTR |
| mouse | 67590 | TCTN3 | NM_026260 | TRCN0000184479 | GCAGAAAGACTCTCTCGTTCT | CDS |
| mouse | 67042 | IFT27/ BBS19 | NM_025931 | TRCN0000102721 | GCCTGGAATTCTTTGAGACAT | CDS |
| mouse | 67042 | IFT27/ BBS19 | NM_025931 | TRCN0000102722 | CCTTGTCTATGATGTGACCAA | CDS |
| mouse | 67042 | IFT27/ BBS19 | NM_025931 | TRCN0000102723 | CTGGATAAGTTGTGGGAGAAT | CDS |
| mouse | 67042 | IFT27/ BBS19 | NM_025931 | TRCN0000102724 | CAGTGCCAGTTCTTGACACAA | CDS |
| mouse | 230967 | CEP104 | NM_177673 | TRCN0000194490 | GCGCAAGACACATGTTCTCTA | CDS |
| mouse | 230967 | CEP104 | NM_177673 | TRCN0000174353 | CATGCCAACAAATACAACATA | CDS |
| mouse | 230967 | CEP104 | NM_177673 | TRCN0000174878 | GCTTCATTGAAATTGCTGAAA | CDS |
| mouse | 230967 | CEP104 | NM_177673 | TRCN0000175806 | GTTATGACCAAAGAGCCCTTT | 3UTR |
| mouse | 226075 | GLIS3/ ZNF515 | NM_175459 | TRCN0000095874 | CGCACGGTATAAACTGCTGAT | CDS |
| mouse | 226075 | GLIS3/ ZNF515 | NM_175459 | TRCN0000095875 | CGGGATCGACTTCAACACTAT | CDS |
| mouse | 226075 | GLIS3/ ZNF515 | NM_175459 | TRCN0000095876 | CCAGCAATCATTCAACTCGAA | CDS |
| mouse | 226075 | GLIS3/ ZNF515 | NM_175459 | TRCN0000095877 | ACTAAACCTTATGCTTGTCAA | CDS |

| | | | | | | |
|---|---|---|---|---|---|---|
| mouse | 226075 | GLIS3/ ZNF515 | NM_175459 | TRCN0000095878 | CTAAACCTTATGCTTGTCAAA | CDS |
| mouse | 53885 | NPHP1 | NM_016902 | TRCN0000190911 | CCCGAAACAAGCAGAAACAGA | CDS |
| mouse | 53885 | NPHP1 | NM_016902 | TRCN0000201440 | CAGGAAGACTTGCCTCAAACA | 3UTR |
| mouse | 53885 | NPHP1 | NM_016902 | TRCN0000200670 | GTTGATAGTTTGGTGACTGAA | CDS |
| mouse | 53885 | NPHP1 | NM_016902 | TRCN0000192281 | CCTGCAAAGTGCCGATTTAAT | CDS |
| mouse | 217935 | DYNC2I1/WDR60 | NM_146039 | TRCN0000191802 | GAGAAAGAAGATACCGAGAAA | CDS |
| mouse | 217935 | DYNC2I1/WDR60 | NM_146039 | TRCN0000191741 | GCAGGCTTTCAAATGATTAAA | 3UTR |
| mouse | 217935 | DYNC2I1/WDR60 | NM_146039 | TRCN0000200755 | GTTGGCGAGTTATCTTTGAAA | CDS |
| mouse | 217935 | DYNC2I1/WDR60 | NM_146039 | TRCN0000192207 | CCGTGAGAAAGACAAGCTAAA | CDS |

### Table S4

| GeneID | Symbol | RefSeq | species | TRC_ID | SenseSeq | Region | shRNA_ID |
|---|---|---|---|---|---|---|---|
| 72982 | TMEM138 | NM_028411 | mouse | TRCN0000249784 | ACGGCCTGCAAACACTGTTTG | CDS | |
| 72982 | TMEM138 | NM_028411 | mouse | TRCN0000249785 | ACTCCGAATGGCTCCTGTTAT | CDS | sh*mTmem138* #1 |
| 72982 | TMEM138 | NM_028411 | mouse | TRCN0000249787 | CCTCTTCAACATCATCATAAT | CDS | sh*mTmem138* #2 |
| 72982 | TMEM138 | NM_028411 | mouse | TRCN0000257914 | GGACTTCCTAGGACACATTTA | 3UTR | |
| 211535 | ODAD1/ CCDC114 | NM_001364168 | mouse | TRCN0000253368 | AGATCTTGCTGCGACAGATAT | CDS | |
| 211535 | ODAD1/ CCDC114 | NM_001364168 | mouse | TRCN0000253369 | CCAAGCTCCAGACGCAGATTA | CDS | |
| 211535 | ODAD1/ CCDC114 | NM_001364168 | mouse | TRCN0000253371 | CCCTAGAGACTCCTTTCTAAT | 3UTR | |
| 100503884 | CCDC149 | NM_001379231 | mouse | TRCN0000183739 | CAATGTCCTGTTTATCTCTAT | 3UTR | |
| 100503884 | CCDC149 | NM_001379231 | mouse | TRCN0000184721 | CCCATTCATCGTAAACCCTGT | 3UTR | |
| 100503884 | CCDC149 | NM_001379231 | mouse | TRCN0000136802 | CGAGAGCTGATTGATGGAGAT | CDS | |

| GeneID | Symbol | Sense_oligo (5' to 3') |
|---|---|---|
| 72982 | TMEM138 | CCGGTACGGCCTGCAAACACTGTTTG**TACTAGT**CAAACAGTGTTTGCAGGCCGTTTTTTG |
| 72982 | TMEM138 | CCGGTACTCCGAATGGCTCCTGTTAT**TACTAGT**ATAACAGGAGCCATTCGGAGTTTTTTG |
| 72982 | TMEM138 | CCGGTCCTCTTCAACATCATCATAAT**TACTAGT**ATTATGATGATGTTGAAGAGGTTTTTG |
| 72982 | TMEM138 | CCGGTGGACTTCCTAGGACACATTTA**TACTAGT**TAAATGTGTCCTAGGAAGTCCTTTTTG |
| 211535 | ODAD1/ CCDC114 | CCGGTAGATCTTGCTGCGACAGATAT**TACTAGT**ATATCTGTCGCAGCAAGATCTTTTTTG |
| 211535 | ODAD1/ CCDC114 | CCGGTCCAAGCTCCAGACGCAGATTA**TACTAGT**TAATCTGCGTCTGGAGCTTGGTTTTTG |
| 211535 | ODAD1/ CCDC114 | CCGGTCCCTAGAGACTCCTTTCTAAT**TACTAGT**ATTAGAAAGGAGTCTCTAGGGTTTTTG |
| 100503884 | CCDC149 | CCGGTCAATGTCCTGTTTATCTCTAT**TACTAGT**ATAGAGATAAACAGGACATTGTTTTTG |
| 100503884 | CCDC149 | CCGGTCCCATTCATCGTAAACCCTGT**TACTAGT**ACAGGGTTTACGATGAATGGGTTTTTG |
| 100503884 | CCDC149 | CCGGTCGAGAGCTGATTGATGGAGAT**TACTAGT**ATCTCCATCAATCAGCTCTCGTTTTTG |

| GeneID | Symbol | Anti-sense_oligo (5' to 3') |
|---|---|---|
| 72982 | TMEM138 | AATTCAAAAAACGGCCTGCAAACACTGTTTG**ACTAGTA**CAAACAGTGTTTGCAGGCCGTA |
| 72982 | TMEM138 | AATTCAAAAAACTCCGAATGGCTCCTGTTAT**ACTAGTA**ATAACAGGAGCCATTCGGAGTA |
| 72982 | TMEM138 | AATTCAAAAACCTCTTCAACATCATCATAAT**ACTAGTA**ATTATGATGATGTTGAAGAGGA |
| 72982 | TMEM138 | AATTCAAAAAGGACTTCCTAGGACACATTTA**ACTAGTA**TAAATGTGTCCTAGGAAGTCCA |
| 211535 | ODAD1/ CCDC114 | AATTCAAAAAAGATCTTGCTGCGACAGATAT**ACTAGTA**ATATCTGTCGCAGCAAGATCTA |
| 211535 | ODAD1/ CCDC114 | AATTCAAAAACCAAGCTCCAGACGCAGATTA**ACTAGTA**TAATCTGCGTCTGGAGCTTGGA |
| 211535 | ODAD1/ CCDC114 | AATTCAAAAACCCTAGAGACTCCTTTCTAAT**ACTAGTA**ATTAGAAAGGAGTCTCTAGGGA |
| 100503884 | CCDC149 | AATTCAAAAACAATGTCCTGTTTATCTCTAT**ACTAGTA**ATAGAGATAAACAGGACATTGA |
| 100503884 | CCDC149 | AATTCAAAAACCCATTCATCGTAAACCCTGT**ACTAGTA**ACAGGGTTTACGATGAATGGGA |
| 100503884 | CCDC149 | AATTCAAAAACGAGAGCTGATTGATGGAGAT**ACTAGTA**ATCTCCATCAATCAGCTCTCGA |

#### Table S5

| Species | Gene | Forward qPCR primer (5' to 3') | Reverse qPCR primer (5' to 3') |
|---|---|---|---|
| mouse | TUBB4A | ATGAGGCCACAGGTGGAAAC | CCTGCTCCGGATTGACCAAA |
| mouse | B2M | ATGGGAAGCCGAACATACTG | CAGTCTCAGTGGGGGTGAAT |
| mouse | TMEM138 | CATGAACGTGCGATGGAAAA | TGTTTGCAGGCCGTTTGTC |
| mouse | PMEL | GCACCCAACTTGTTGTTCCT | GTGCTACCATGTGGCATTTG |
| mouse | TYRP1 | AAGTTCAATGGCCAGGTCAG | TCAGTGAGGAGAGGCTGGTT |
| mouse | MITF | GCAAGAGGGAGTCATGCAGT | AGAACTGCTGCTCTTCAGAGGT |
| mouse | MC1R | CTCCACAGACCGCTTCCTAC | ACATACAGGCACCAAGGCTC |
| mouse | OCA2 | CACAGAGTAGAGTGGGCGAC | GCAAAGCGCTGATCTTCTGG |
| mouse | TYR | CCAGAAGCCAATGCACCTAT | ATAACAGCTCCCACCAGTGC |
| mouse | LAMP2 | CAAAAGGACAGTATTCTACAGCTCA | TGATGGCGCTTGAGACCAAT |
| mouse | HPS9 | CTCCAGACGGGGTCCTTAC | CACGTCTCCAGATGAAGGACT |
| mouse | CALR | GAATACAAGGGCGAGTGGAAACC | GACAACCCTGAATACTCCCCCG |
| mouse | DCT | AGAGAAACAACCCTTCCACAGA | CCAATGACCACTGAGAGAGTTG |
| mouse | TMEM216 | CGTGCTCCGCCTAGAAGCTA | AGCAGCTCTGAGCCACAAAAG |
| mouse | BBS16 | GCTGTGGAGAAGGAGATGATAAA | GCCTCCAGCTTGGCAATA |
| mouse | IFT172 | CAGTTGAAGCACCTGAGGAC | TCCACTGTGCAGACAGCAAA |
| mouse | INPP5E | ACTGGACTACAGCAGAACCAT | ACCACTCAGGCGGAAGTTG |
| mouse and human | MITF | AGGACCTTGAAAACCGACAG | GTGGATGGGATAAGGGAAAG |
| human | TUBB | ATGAAGCCACAGGTGGCAAA | CTGCCCCAGACTGACCAAAT |
| human | GAPDH | AATCCCATCACCATCTTCCA | TGGACTCCACGACGTACTCA |
| human | TMEM138 | CATCCTGACAGCTGTGTACTTTGC | CGTAAGTTCATGACCCAGACATG |
| human | TYR | GGCTGTTTTGTACTGCCTGCT | AGGAGACACAGGCTCTAGGGAA |
| human | TYRP1 | GTGCCACTGTTGAGGCTTTG | ATGGGGATACTGAGGGCTGT |
| human | MC1R | ACTTCTCACCAGCAGTCGTG | CATTGGAGCAGACGGAGTGT |
| human | PMEL | AACTCCAGAGGCTACAGGTATG | GTCTCCACCCACTCTGTAGTT |
